# Molecular responses of chicken embryos to maternal heat stress through DNA methylation and gene expression

**DOI:** 10.1101/2024.04.12.589068

**Authors:** Keyvan Karami, Jules Sabban, Chloé Cerutti, Guillaume Devailly, Sylvain Foissac, David Gourichon, Alexandre Hubert, Jean-Noël Hubert, Sophie Leroux, Tatiana Zerjal, Sandrine Lagarrigue, Frédérique Pitel

## Abstract

Climate change, with its repercussions on agriculture, is one of the most important adaptation challenges for livestock production. Poultry production is a major source of proteins for human consumption all over the world. With a growing human population, improving poultry’s adaptation to environmental constraints becomes critical. Extensive evidence highlights the influence of environmental variations on epigenetic modifications. The aim of this paper is therefore to explore chickens’ molecular response to maternal heat stress. We employed Reduced Representation Bisulfite Sequencing (RRBS) to generate genome-wide single-base resolution DNA methylation profiling and RNA sequencing (RNA-seq) to profile the transcriptome of the brains of embryos hatched from dams reared under either heat stress (32 °C) or thermoneutrality (22°C). We detected 289 significant differentially methylated CpG sites (DMCs) and one differentially methylated region (DMR) between heat stressed and control groups. These DMCs were associated with 357 genes involved in processes such as cellular response to stimulus, developmental processes and immune function. In addition, we identified 11 genes differentially expressed between the two groups of embryos, and identified ATP9A as a target gene of maternal heat stress on offspring. This study provides a body of fundamental knowledge on adaptive mechanisms concerning heat tolerance in chickens.

## Introduction

Climate change and its direct and indirect consequences represent one of the most important adaptation challenges for livestock production, as unpredictable and rapid environmental changes are a source of stress. Chicken meat and eggs are major sources of proteins for human food worldwide, but their production is affected by global warming. Rising temperatures have adverse effects on poultry growth, production and survival. It has been shown that heat stress causes a decrease in productivity in many species^1–3^ . Heat stress in chickens, as in other species, leads to reduced feed consumption, resulting in decreased energy and nutrient intake. This ultimately leads to compromised growth and reduced quality of broiler products, as well as decreased egg quantity and quality in layers^4–9^. The increased demand for animal products worldwide combined with a growing human population urges the need to improve the ability of animals to respond to heat stress^10^. Research has demonstrated that the environment exerts influence on gene expression in both plants and animals, resulting in phenotypic plasticity; this phenomenon leads to the emergence of different phenotypes from the same genotype in response to different environmental conditions, and can even affects the phenotype of future generations through transgenerational plasticity^11–13^. Some of these effects are mediated by epigenetics phenomena: in response to the environment, epigenetic mechanisms can induce changes in gene expression, linking environmental changes to the physiology and health of animals^14,15^. These mechanisms may act as catalysts and trigger the adaptation of organisms to their environment.

Epigenetics covers all mechanisms that modify gene expression in a reversible and transmissible way through mitosis or meiosis, without modifying the DNA sequence^16^. These phenomena include DNA methylation, histone modification, remodeling of chromatin, and regulation of gene expression by non-coding RNAs (ncRNAs). Numerous studies, particularly in humans and mammals, showed that maternal stress can lead to epigenetic alterations in offspring, which ultimately may affect their phenotype^17,18^.

In avian species, Tzschentke and Basta (2002) reported that, in ducks, prenatal temperature experience has a clear influence on postnatal neural hypothalamic thermosensitivity and could be the result of epigenetic temperature adaptation^19^. In chickens, research focused on the effect of thermal manipulations during embryogenesis on post-hatch heat tolerance and showed an increased heat tolerance in broilers within the first 5 weeks of life, when exposed to an acute heat stress^20,21^. In Japanese quails, a study by Vitorino Carvalho et al. (2020) reported that thermal manipulation during embryogenesis significantly reduced the hatching rate and increased mortality during the first four weeks of life^22^. Subsequent research (Vitorino Carvalho et al., 2021) reported that thermal manipulation during embryogenesis had little to no effect on gene expression regulation in the hypothalamus of 35-day-old quails^23^. On the contrary, exposure to a heat challenge before this sampling resulted in an increase in the number of differentially expressed genes, reinforcing the hypothesis that embryonic thermal conditioning has a beneficial effect and increases thermotolerance later in life^10,21,24^.

The response to heat stress can also be triggered by heat exposure in the previous generation. For example, Ahmed et al., (2017) reported that maternal heat stress during late gestation increased acute thermal tolerance of the calf at maturity^25^. In birds several studies have also tried to elucidate the effect of the environmental experience of mothers on their offspring. In Japanese quails, it has been reported that maternal stress may affect and prepare future generations to cope with later environmental difficulties^26,26^. Santana et al. (2021) reported that maternal stress led to lower laying rate, egg mass and higher chick mortality rate at the 1–15 days of age. They observed that the performance and oxidative metabolism of offspring raised in thermoneutral conditions were unaffected by maternal heat stress, while offspring subjected to heat stress during growth showed increased levels of protein oxidation^18^. In a recent study^27^, it was shown that thermal manipulation repeated during 4 generations in Japanese quail had a transgenerational effect on body weight and egg weight, suggesting non-genetic inheritance mechanisms. The hypothesis made to justify the improved resistance was that heat stress-induced epigenetic modifications were occurring as a consequence of the embryonic thermal manipulation, leading to increased thermal tolerance and adaptability in adults. A recent study confirmed the epigenetic nature of the transmission of heat-induced effects between generations through epigenetic mechanisms in chicken^28^.

Unlike mammals, birds have not been extensively studied for the effect of maternal heat stress on offspring heat tolerance. In this study, we explored this aspect by analyzing the genome-wide methylation and transcriptomic profiling of embryos whose mothers were reared under high ambient temperatures or under thermoneutral conditions. The underlying hypothesis is that maternal heat stress induces changes in DNA methylation in chicken embryos leading to changes in gene expression.

## Results

In order to assess the epigenomic response to maternal heat stress on the DNA methylation levels in 13-day-old embryos, 22 embryos (10 controls and 12 stressed) were analysed. The results showed that heat stress of hens can mediate changes in the methylation patterns and also differential expression of some genes in offspring.

### DNA methylation changes

Some general statistics of RRBS sequencing results are summarized in Table S2. An average of 20 million reads per sample were obtained.. The average mapping efficiency was 64.84%, in accordance with what is expected from this type of data^29^. We have assessed 1,075,291 CpG sites (after preprocessing; Fig 2A) with an average depth of 18.34. The distribution of methylation level around the transcription start site (TSS) showed a decreased value in this region (Fig 2B). Among the analysed CpGs, we detected a total of 289 DMCs between HS and CT groups, of which 138 were hypermethylated and 151 were hypomethylated in the HS group (Fig 3). The DMCs were present along most chromosomes (Fig 4 and Fig S2). Their distribution was not constant along the genome and some regions had a high density of DMCs. Notably, one region on chromosome 4 (Chr4:2858109,2858165) was identified as a DMR. This region harbored two lncRNA genes (LOC121110553, LOC121110554) with unknown functions. As shown in Fig 5 these two genes have contrasted expression patterns across 47 tissuesl^30^, and only LOC121110553 was expressed in embryo.

**Fig 1:**
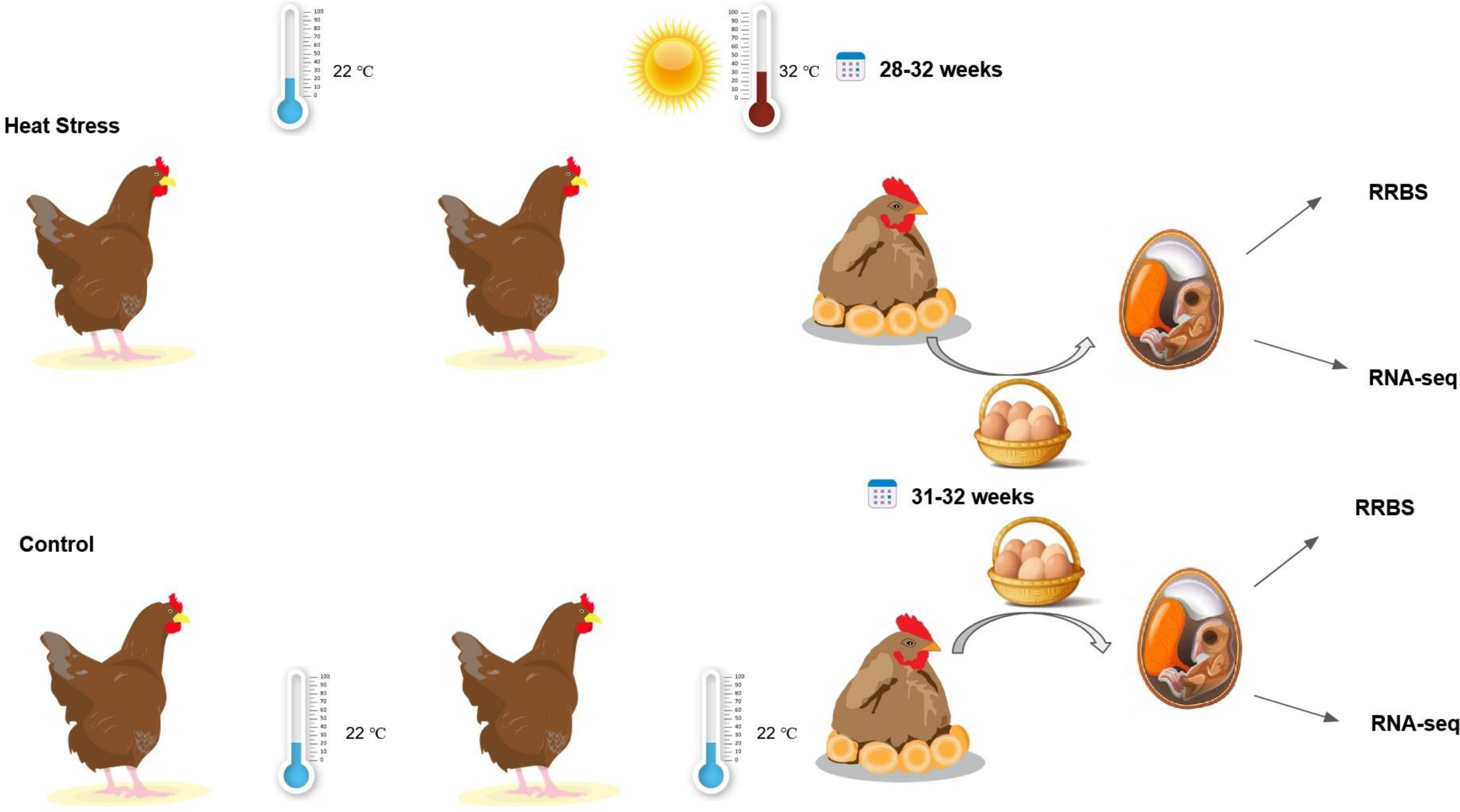
Experimental design

**Fig 2:**
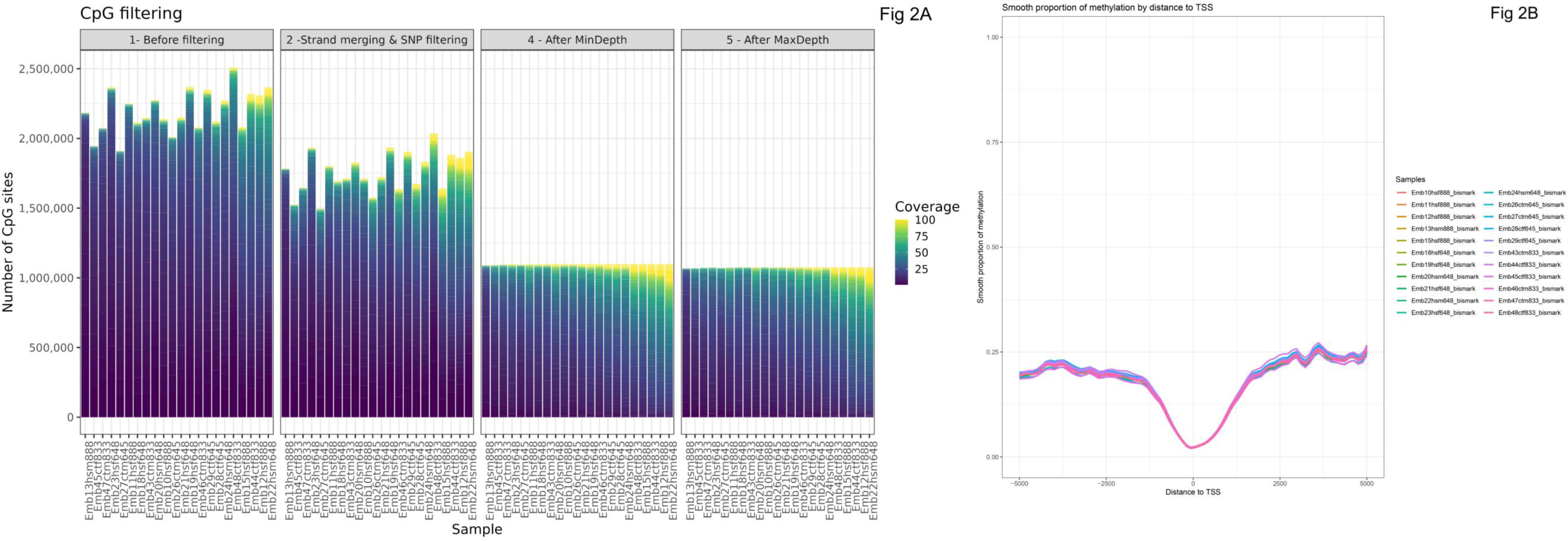
Preprocessed data A) Number of CpGs kept after each step of the pre-processing workflow. B) Average methylation level around TSS regions.

**Fig 3:**
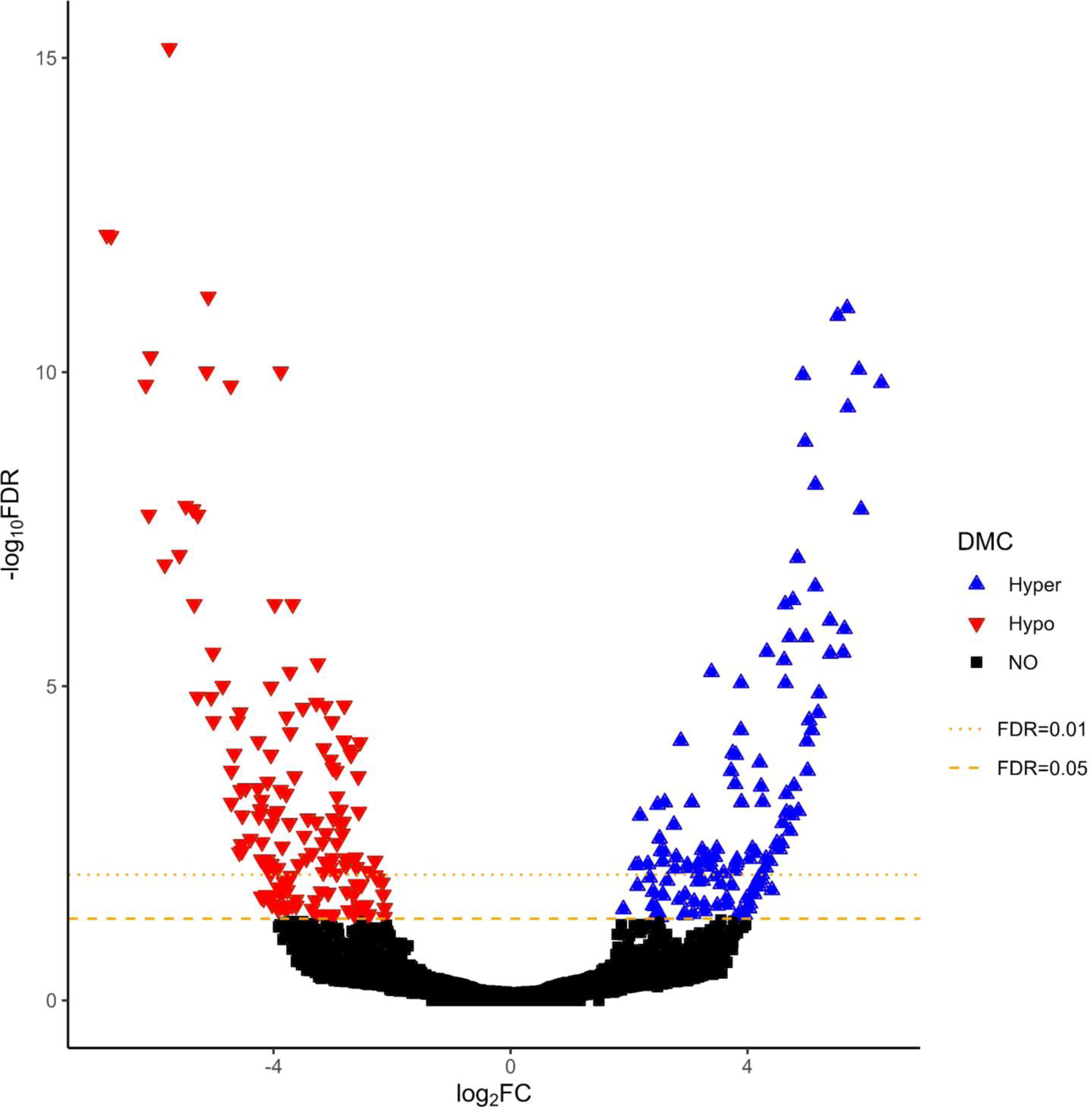
Volcano plot of CpG methylation and DMCs between HS and CT. Hyper=hypermethylated; Hypo=hypomethylated, FDR=False Discovery Rate, DMC=differentially methylated CpG site

**Fig 4:**
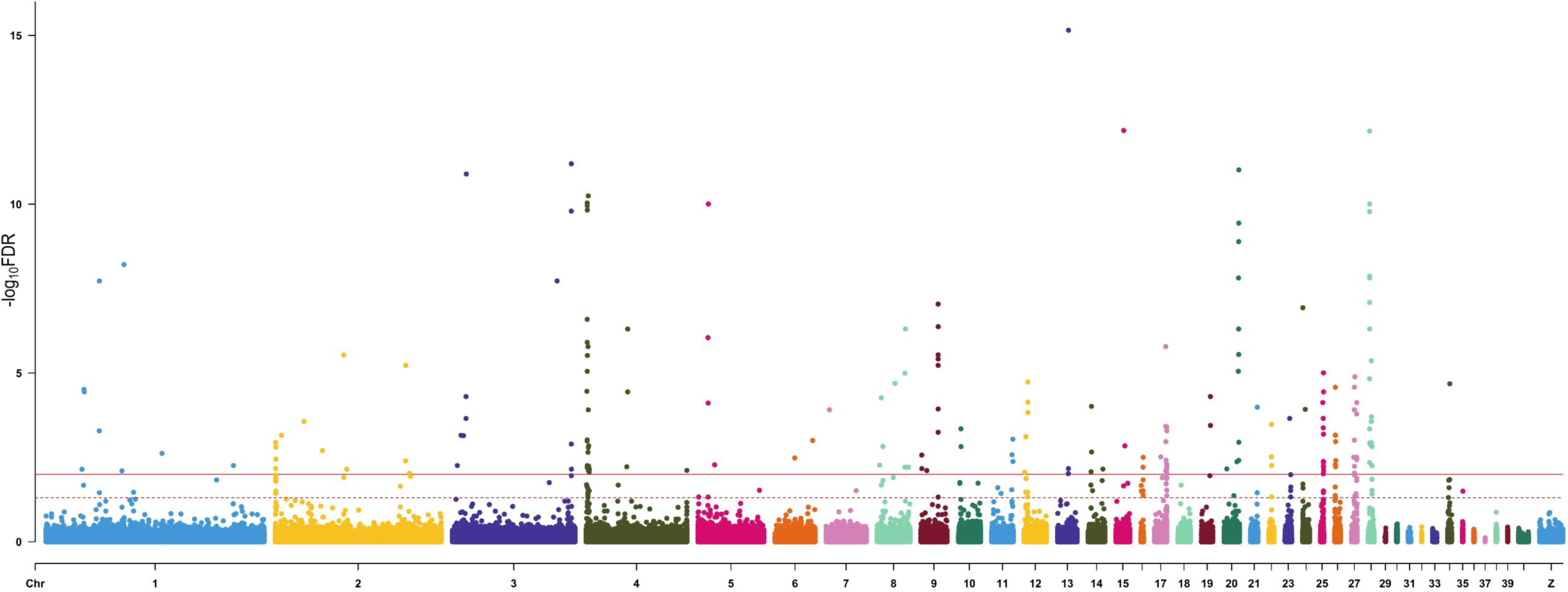
Manhattan plot of differential methylation analysis between HS and CT groups. The above dashed line represents FDR ≤ 0.05 and solid line represents FDR ≤ 0.01. *FDR=False Discovery Rate*

**Fig 5:**
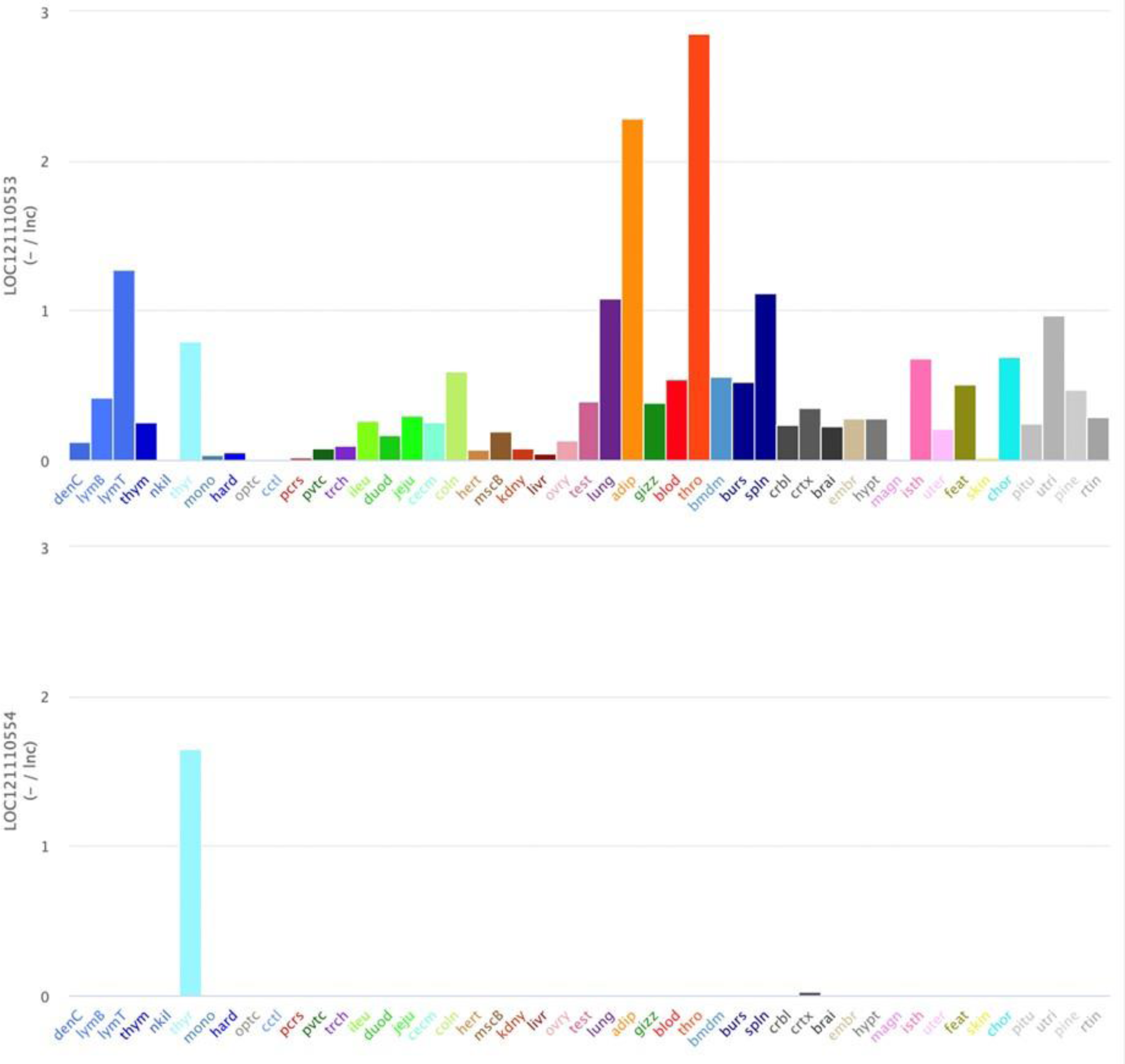
Expression pattern of two lncRNA genes (LOC121110553, LOC121110554) across 47 tissues (https://gega.sigenae.org/)

### Annotation of differentially methylated cytosine

DMCs were annotated according to gene features. From the detected DMCs, 28.85% were located in promoter regions, 40.28% in introns and 18.42% in exons (Fig 6). Chi2 test showed that these distributions among CpGs and DMCs (p-value < 2.2e-16) and among hyper and hypo DMCs (p-value < 2.2e-16) were significantly different. The fraction of the DMCs located in the promoter region was more frequently hypermethylated (37.25%) than hypomethylated (19.98%), while hypomethylation was more frequent in exons and introns.

**Fig 6:**
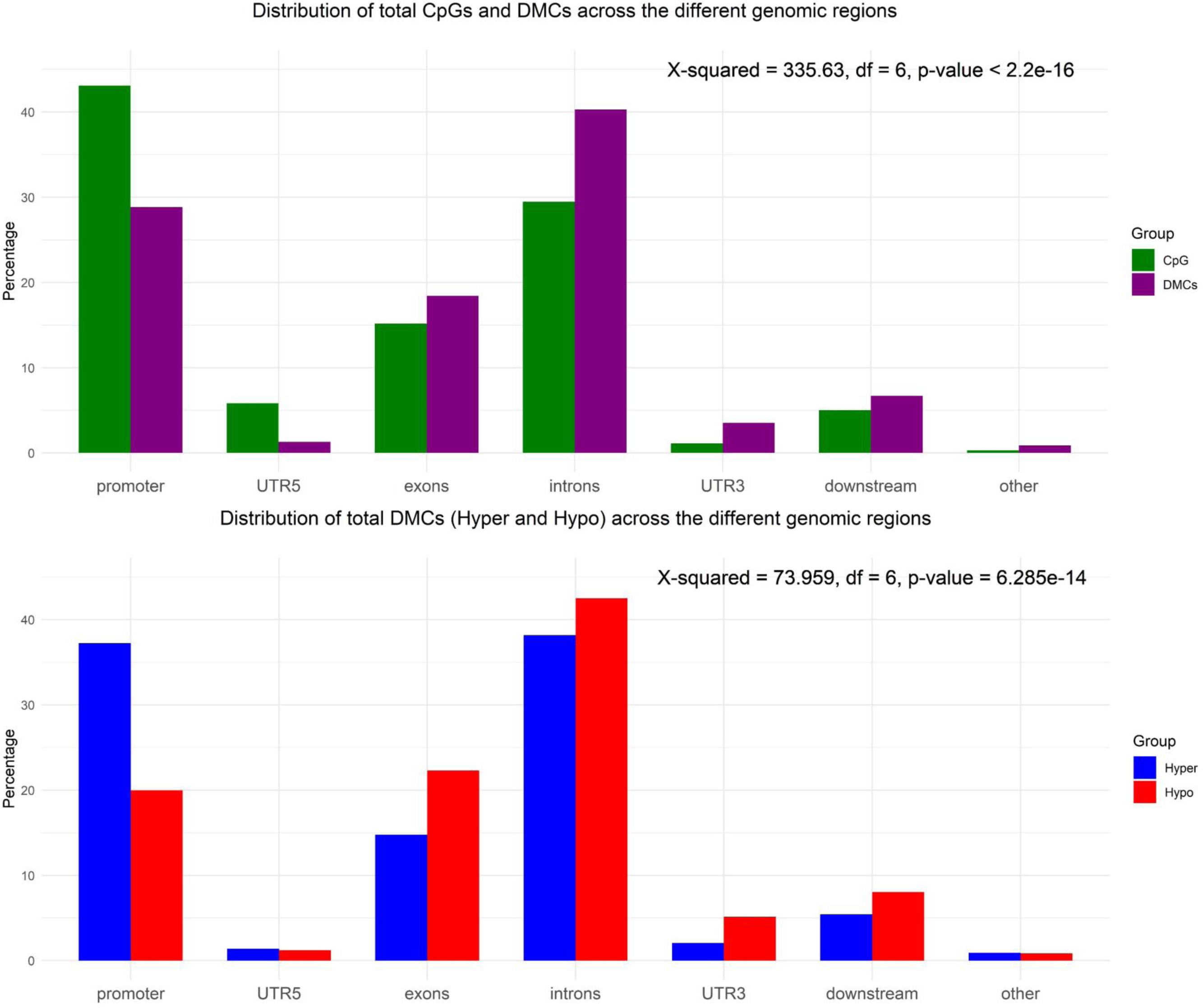
Distribution of total CpGs and DMCs (hypermethylated and hypomethylated) across the different genomic regions.

### Gene ontology functional analysis

Based on the DMCs location, we identified 357 differentially methylated genes (DMGs) that harbored at least one DMC in one of the gene features considered (Table S3) out of 35,995 genes with at least one CpG. The functional analysis of these genes has enabled us to identify as enriched several biological processes (BP) linked to the development stage. The gene ontology ViSEAGO output showed also the significance of embryo development, metabolic process, cellular response to stimulus, immune function (Fig 7).

**Fig 7.**
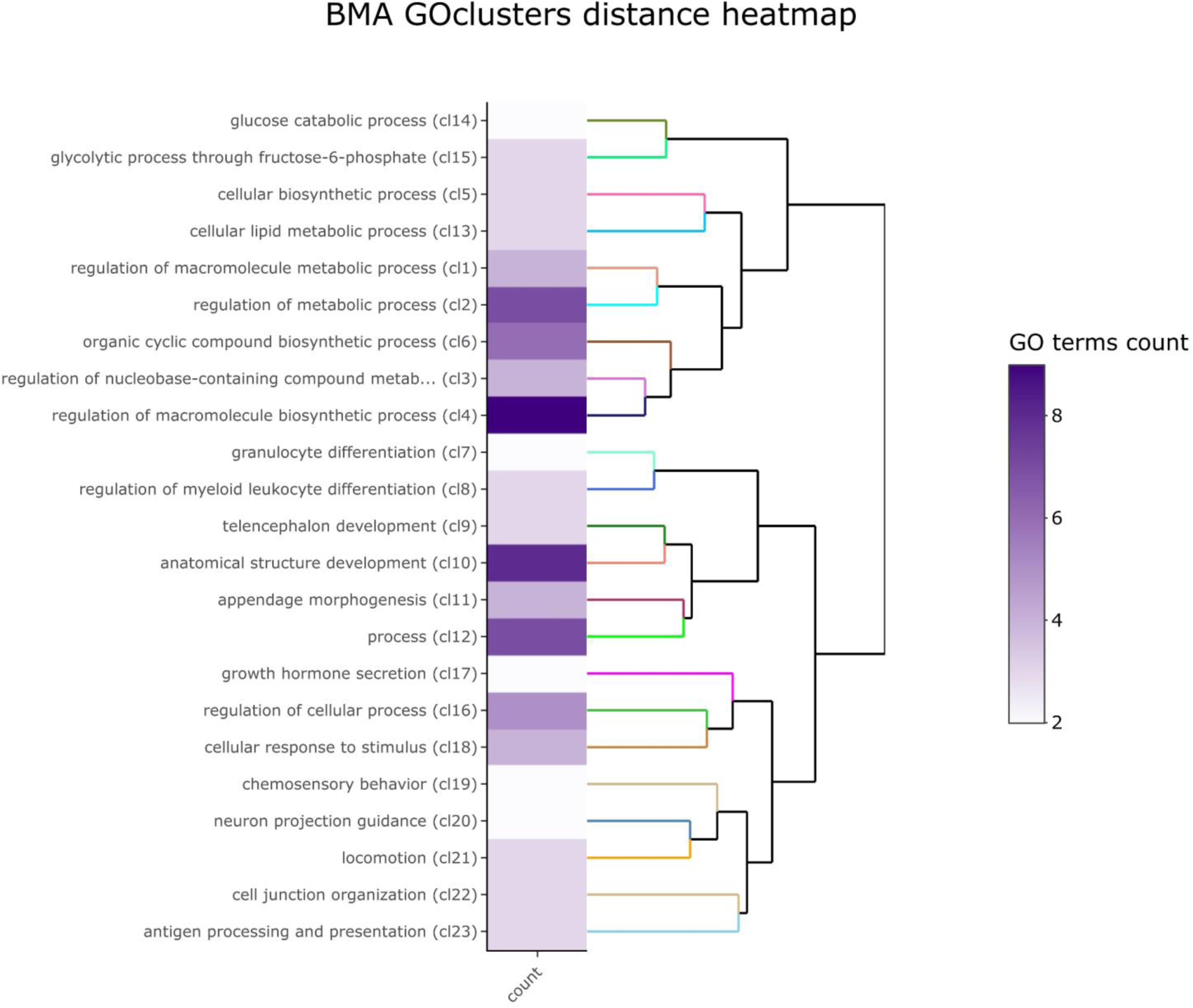
: Gene Ontology functional analysis of the genes related to DMCs. The clustering heat map plot of the functional sets of gene ontology (GO) terms was obtained using ViSEAGO. Gene Ontology functional analysis with count showing information content and a dendrogram on enriched GO terms based on BMA semantic similarity distance and Ward’s clustering criterion.

### Gene expression analysis

RNA sequencing analysis was performed to investigate the impact of heat stress on embryo gene expression. Among the 17,939 genes identified as expressed in embryos, eleven DEGs were detected between HS and CT groups as listed in Table 1, all being protein coding genes. Among these, four genes were upregulated and seven genes were down regulated. ATP9A (ATPase phospholipid transporting 9A), one of the upregulated genes in the HS embryos, was also in the list of DMGs, with 4 DMCs in the introns and exon regions, all of them being hypermethylated (Fig 8).

**Fig 8:**
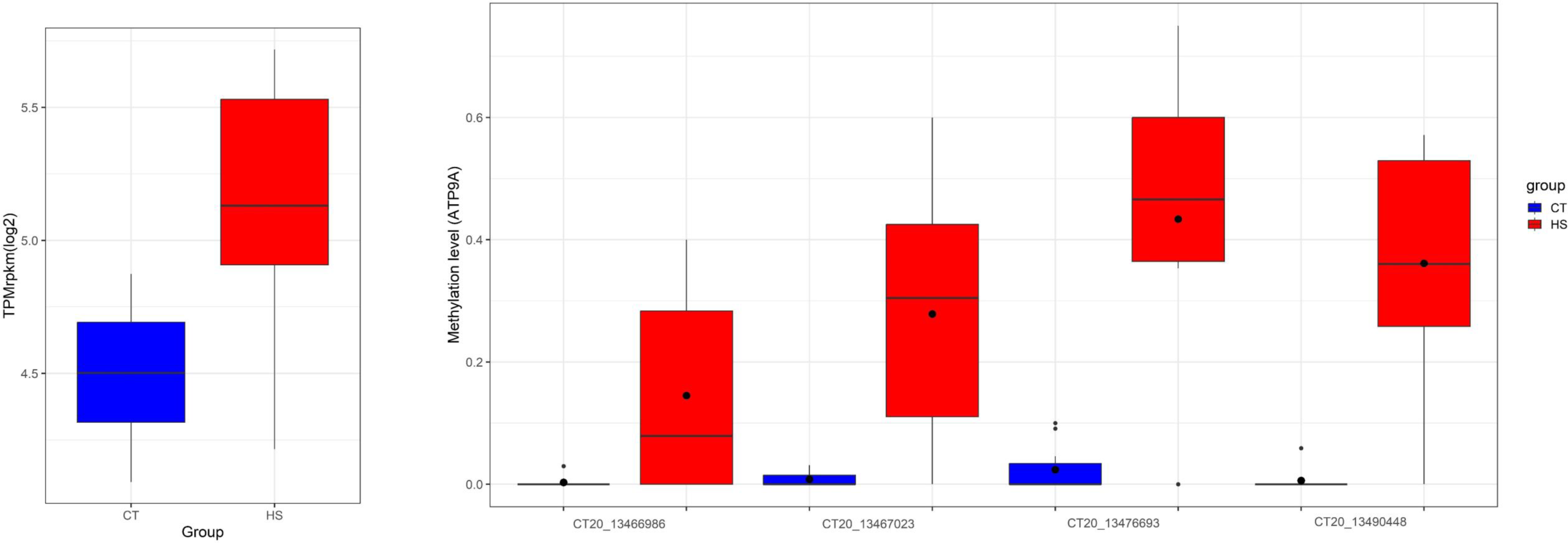
Expression and methylation level of the 4 DMCs per group (CT and HS) for ATP9A.

**Table 1.** Differentially expressed genes between Heat Stress and Control groups.

| Gene id | Gene Name | Chr | start | end | strand | Expression* | padj | fc | lfc |
| --- | --- | --- | --- | --- | --- | --- | --- | --- | --- |
| LOC100858942 | LOC100858942 | 34 | 2198709 | 2201885 | + | UP | 0.01 | 242.44 | 7.92 |
| LOC112531412 | LOC112531412 | JAENSK010000420.1 | 8859 | 18598 | - | UP | 0.03 | 98.82 | 6.63 |
| LOC396217 | MBP | 2 | 90091375 | 90199666 | + | UP | 0.02 | 12.92 | 3.69 |
| LOC419345 | ATP9A | 20 | 13450938 | 13503307 | + | UP | 0.01 | 2.57 | 1.36 |
| LOC107054346 | LOC107054346 | 12 | 1193659 | 1196336 | + | DOWN | 0.04 | 0.04 | -4.48 |
| LOC121108245 | LOC121108245 | Z | 169136 | 201956 | - | DOWN | 0.01 | 0.08 | -3.69 |
| LOC100857335 | LOC100857335 | 34 | 1513723 | 1516547 | - | DOWN | 0.01 | 0.1 | -3.34 |
| LOC107057116 | ZNFY4 | 16 | 1583937 | 1595533 | + | DOWN | 0.01 | 0 | -12.35 |
| LOC121108653 | LOC121108653 | MU179258.1 | 33562 | 38085 | + | DOWN | 0.01 | 0.33 | -1.62 |
| LOC100502566 | TMSB15B | 4 | 1940045 | 1942512 | + | DOWN | 0.00 | 0.34 | -1.57 |
| LOC417488 | CLIP2 | 19 | 3258667 | 3318184 | - | DOWN | 0.00 | 0.41 | -1.28 |

\*Up: more expressed in HS than in CT, Down: less expressed in HS than in CT. fc: fold change, lfc: log<sub>2</sub>(fold change)

### Pyromark validation

Pyrosequencing validation of seven DMCs with PyroMark confirmed all the positions as DMCs. Fig 9 shows the methylation level obtained with RRBS and PyroMark.

## Discussion

The livestock industry faces a growing number of challenges due to climate change and global warming, which have a direct impact on animal growth, reproduction, health, and welfare. The exposure of animals to climate changes and other associated stressors has both short- and long-term effects over the course of the animal’s life. There is growing evidence that epigenetics, in interaction with the environment, may also contribute to the phenotypic diversity of animals^31^. In addition, these effects can be passed across generations with multigenerational inheritance and perhaps provide the ability to adapt to climate change for the subsequent generations^32,33^.

Our study aimed to elucidate the effect of maternal heat exposure on DNA methylation and gene expression in chicken embryos. The results revealed a slight influence of maternal heat stress on embryo transcriptomic levels, with eleven differentially expressed genes. We detected a total of 289 DMCs between HS and CT groups, consistent with findings from previous studies in chicken^28^, cow^34^ or guinea pig^35^, which have demonstrated changes in DNA methylation linked to parental heat exposure.

We observed that promoter DMCs were more frequently hypermethylated than hypomethylated in contrast with what was observed in exon and intron regions. This suggests that the promoter region may be more prone to hypermethylation in response to the mother heat stress than the other parts of the genes. A slight similar trend was observed in rainbow trout sperm after heat exposure of males during spermatogenesis^36^.

We identified 357 DMGs containing at least one DMC in various gene features, with a number of 6 DMCs per gene on average. In contrast, only 11 genes exhibited significant differential expression. This highlighted the observation that the majority of differential methylation sites are not simultaneously associated with changes in gene expression. Such finding is consistent with the well-established knowledge that gene expression is highly context dependent, presenting a very fine tissue and stage specificity^37^. The lack of association at this developmental stage does not exclude a potential functional impact of methylation marks on gene expression later in life, which could facilitate responses to heat stress exposures. It is indeed expected that during embryogenesis, some epigenetic marks are programmed and largely maintained throughout development, contributing to better cope with environmental stressors later in life^38^ (Skinner, 2011).

Among the identified DMGs, ERBB4 (Erb-B2 Receptor Tyrosine Kinase 4), NFATC2 (Nuclear Factor Of Activated T-Cells 2) and ATP9A (ATPase Phospholipid Transporting 9A) have been linked to GWAS signals associated with thermotolerance in pigs, as reported by Kim et al., (2018)^39^. Another study by Ramírez-Ayala et al., (2021) linked the ATP9A gene to thermogenesis in cattle^40^. Interestingly, in our study, ATP9A emerged as both DMG and DEG, and harbored numerous DMCs in both its intronic and exonic regions. This observation suggests the existence of temperature regulation pathways potentially shared between mammals and birds.

The DMR on chromosome 4 is associated with two long non-coding RNAs whose function has yet to be characterized: LOC121110553 is weakly expressed but not differentially expressed between the two groups, while LOC121110554 does not appear to be expressed.

The gene ontology analysis of DMGs identified important biological processes including cellular response to stimulus, embryo development, and telencephalon development. Cellular response to stimulus encompasses any process that alters the state or activity of a cell, such as movement, secretion, enzyme production, or gene expression. Indeed, cellular reaction to stress is diverse, ranging from activation of pathways involved in survival strategies to programmed cell death, which eliminate damaged cells^41^. Cellular apoptosis was reported as upregulated after a longer period of heat stress in highland and lowland chicken^10^. The cell’s initial reaction to a stressful stimulus tends to support its defense and recover from injury. However, if distressing stimuli persist without resolution, cells activate signaling pathways leading to programmed cell death^41^ .

Adaptive immune response is another pathway that was associated with DMGs. Heat stress in commercial laying hens has been shown to reduce production performance and inhibit immune function leading to an increase in mortality^42^. Similarly, a study showed that HS causes immune abnormalities in broiler chickens by impairing T and B cell development and maturation in primary and secondary lymphoid tissues^43^. In another study, transcriptome analysis revealed the genes and pathways involved in bursal responses to heat stress and lipopolysaccharide, showing that the combined treatments had the greatest effect^44^. The negative link between heat stress and immune function was also observed in cattle. For example, Dahl et al., (2020) reported that lactating cows often exhibit higher disease incidence in summer (metritis, mastitis, respiratory disease), possibly linked to compromised immune cell activity due to heat stress^45^. Additionally, calves born to mothers experiencing heat stress and dry period during the late gestation had lower weight at birth and through puberty^46–48^.

Epigenetics has the capability of conveying information to next generations without DNA sequence alteration. Epigenetic marks may represent the signature of environment stresses and specific physiological states acquired by the parental generation that could enhance adaptability of next generations to new situations. The outcome of the current study illustrates that maternal exposure to heat stress has an effect on the DNA methylation pattern of offspring. However, even with the exclusion of observed SNPs at CpG sites, we cannot rule out the hypothesis that some of the identified DMCs may be caused by genetic polymorphisms. Although these methylome changes were not associated with extensive transcriptional changes at the embryonic level, the affected genes and pathways identified from differentially methylated genes suggest a potential foundation for adaptive responses in progeny. This aligns with the studies of McGuigan et al. (2021) and Weyrich et al. (2016), indicating that under conditions of climate change and stressful environments, epigenetic factors, through intergenerational and transgenerational effects, play a role in promoting adaptability of exposed populations^33,35^. This has been observed in chicken, with an intergenerational inheritance of heat resilience after fathers’ embryonic heat conditioning, associated with DNA methylation changes in anterior preoptic hypothalamus^28^.

This study shows that maternal exposure to heat stress can induce hundreds of changes in methylation level and minor changes in transcriptome level in offspring. These DNA methylation modifications during the embryonic development as a consequence of their mother’s heat stress may provide the capability of an adaptive response to subsequent heat stress exposure.

## Materials and Methods

### Sample preparation and experimental design

A total of 4 hens (2 controls and 2 heat-stressed) from an experimental layer population (R-) issued from selection for feed efficiency^49^ were used. All birds were reared under standard conditions (22°C, *ad libitum* feeding) at the INRAE UE 1295 PEAT Poultry Experimental Unit (Nouzilly). In the heat stress group (Fig 1), hens were reared at 22°C until 28 weeks of age. Between 28 and 32 weeks of age, the hens were kept at 32°C (increasing by 2°C per hour for 5 hours). The four hens were inseminated by the same male at week 30. Their eggs were collected between 31 and 32 weeks and incubated for 13 days.

The experiments were carried out at the PEAT experimental unit under license number C37-175- 1 for animal experimentation, in compliance with European Union legislation, and were approved by the local ethics committee for animal experimentation (Val de Loire) and by the French Ministries of Higher Education and Scientific Research, and Agriculture and Fisheries (n°2873- 2015112512076871), complying with the ARRIVE guidelines.

### DNA and RNA extraction

DNA and RNA from brain of 13-day-old embryos were extracted, according to the manufacturer’s instructions, with AllPrep DNA/RNA Mini Kit (Qiagen catalog No. / ID: 80204). Total RNA and DNA were quantified with a NanoDrop ND-1000 spectrophotometer (Thermo Scientific). The dsDNA concentration was measured using the Quant-iT PicoGreen dsDNA (Invitrogen) assay according to the manufacturer instructions. The fluorometric measurements were performed using ABI7900HT (Applied Biosystem).

The RNA quality was controlled using an Agilent 2100 bioanalyzer (Agilent Technologies France) with the Eukaryote Total RNA Nano Assay. Results were analysed with the 2100 Expert Software. RNA integrity (RIN) was 9.9 on average.

### Reduced representation bisulfite sequencing

We obtained Reduced Representation Bisulfite Sequencing (RRBS) data from whole brains of 22 embryos of unknown sex (10 controls and 12 stressed) at 13 days of age, derived from R- hens with or without heat stress. RRBS libraries were prepared using the Premium RRBS Kit (Diagenode, #C02030033), according to the manufacturer’s instructions. Briefly, the protocol consisted in the digestion of 100 ng of genomic DNA by the *Msp*I enzyme followed by fragment end repair, and addition of adaptors. A size selection step was performed with AMPure XP Beads (Beckman Coulter). Next, samples were quantified by qPCR and the Ct values were used to pool samples by equimolarity. Then the bisulfite conversion was realized on the pool and the final libraries were amplified using MethylTaq Plus Master Mix (Diagenode kit). After a clean-up with AMPure XP Beads, the RRBS library pools were analysed with the Qubit dsDNA HS Assay Kit (Thermo Fisher Scientific), and the profile of the pools was verified using the High Sensitivity DNA chip for 2100 Bioanalyzer (Agilent) or DNF-474 NGS fragment kit on a Fragment Analyzer (Agilent). Libraries were sequenced in single-end mode of 50 bp on an Illumina HiSeq 4000 on the GenomEast platform (https://www.igbmc.fr/en/plateforms-and-services/platforms/genomeast).

### Bioinformatics analyses

The nf-core/methylseq pipeline^50^ version 2.1.0 was used for analysing methylation bisulfite sequencing data. Bismark version 0.24.2 with Bowtie2 as an alignment tool was used for mapping on the *Gallus gallus* genome GRCg7b obtained from Ensembl (bGalGal1.mat.broiler.GRCg7b, https://ftp.ensembl.org/pub/release-109/fasta/gallus_gallus/dna/Gallus_gallus.bGalGal1.mat.broiler.GRCg7b.dna.toplevel.fa.gz).

Pipeline’s default parameters were used, with the option --clip_r1 3 for adapter trimming (trimming 3 bases from the 5’ end of each read).

### Differential methylation analyses

The Bioconductor package edgeR v3.28.1^51^ was used to detect differentially methylated CpGs sites (DMCs), The callDMR function from the DSS package v2.38.0^52^ was used to call DMRs (differentially methylated regions) from the edgeR outputs. A DMR was defined as a region with a minimum number of 3 CpGs and a percentage of CpG sites with significant p-values (less than 0.05) greater than 50% between Heat Stress (HS) and Control (CT) groups. Here a two-step process has been implemented: preprocessing and differential methylation analysis. During the preprocessing step, CpGs that overlapped with C-T single nucleotide polymorphisms (SNPs) were filtered out to avoid erroneous identification of C-T polymorphisms as methylation changes. SNPs were detected by gemBS v4.053 with option “bs_call”, CpGs were further filtered using other criteria (maximum Coverage: 200, minimum coverage: 5 and minimum fraction of samples present per position: 0.8). Differential methylation analysis was performed with edgeR using a multifactor model (HS/CT and Sex) with False Discovery Rate (FDR) ≤ 0.05. Identification of the sex of embryos was performed through the average of read mapped on sex chromosomes (Fig S1).

Genomic features annotation was done with the GenomeFeatures package version 1.3 (https://forgemia.inra.fr/aurelien.brionne/GenomeFeatures) with default defined promoter region upstream:3000 bp and downstream:500 bp. An in-house enriched annotation file was used in this study^54^.

### Functional enrichment analysis

We analyzed all the genes that had at least one DMC in their genomic features (promoter, UTR5, introns, UTR3, downstream). Functional enrichment analysis was done with the R package ViSEAGO v1.14.0^55^, and the full list of genes having at least one CpG in genomic features was used as background.

### RNA-seq data acquisition

Paired-end sequencing was performed using an Illumina HiSeq3000 (Illumina, California, USA) system, with 2 × 150 bp, as in Jehl et al, 2019^56^. FASTQ files were mapped on the GRCg7b reference genome (GCF_016699485.2) and the nf-co.re/rnaseq^50^ pipeline version 3.8.1 was used for providing raw count and transcript per kilobase million (TPM) normalized expression per gene and sample.

### RNA-seq analysis

The normalized expression level was obtained using the trimmed mean of M-values (TMM) scaling factor method, implemented in Bioconductor package edgeR version 3.32.1, with the functions of “calcNormFactors” and “rpkm” used to scale the raw library sizes and scale of gene model size respectively. In situations where TPM and TMM normalized expressions were ≥ 0.1 and read counts ≥ 6 in at least 80% of the samples, the gene was considered as expressed. For differential expression analysis we used the raw counts from the expressed genes previously selected and normalized by the TMM method. The Bioconductor package edgeR was used to perform the differential expression analysis, which is based on a generalized negative binomial model for model fitting. The method of “edgeR-Robust” was used to account for potential outliers when estimating per gene dispersion parameters. P-values were corrected for multiple testing using the Benjamini-Hochberg approach to control the false discovery rate (FDR), and FDR < 0.05 was used to identify significant DEG (Differentially Expressed Gene).

### Pyromark validation

For the DMC validation, the Pyrosequencing method was used to perform a quantitative methylation analysis of bisulfite-converted DNA for each individual. The pyrosequencing was performed using PyroMark Q24 (QIAGEN). All the primers (forward, reverse and sequencing primers) were designed with the PyroMark Assay Design software (Version 2.0.1.15, Qiagen) using the assay type "Methylation Analysis" (CpG) (Table S1).

The PCR reaction contained 2 µl of bisulfite treated DNA sample (EZ DNA Methylation-Gold kit, Zymo Research), 2.5 µl of buffer + 0.05 µl of Taq Polymerase (PCRBIO Classic Taq, Eurobio), 2.5 μl of dNTP (2mM, Promega), 1 μl of each primer (10 μM), and 5.95 μl of water. The program on the thermal cycler (Thermocycleur ABI2720, Applied Bisystem) was: 95 °C for 5 min; followed by 35 cycles of: 95 °C for 30 sec, hybridization temperature for 30 sec, and 72 °C for 30 sec; and a final extension at 72 °C for 5 min.

Ten μl of PCR product were then mixed with 1 μl of Streptavidin sepharose™ high performance (GE Healthcare) and 40 μl of PyroMark binding buffer (Qiagen). The mix was shaken at 1400 rpm on a microplate mixer for at least 10 min. The immobilized PCR products were purified using PyroMark Q24 vacuum workstation (manufacturer instructions, QIAGEN), mixed with 1 μl of a sequencing primer (5µM) and 24 μl of Pyromark annealing buffer, and heated at 80 °C for 5 min to anneal the sequencing primer before analysis on the PyroMark Q24. Results were analysed with the PyroMark Q24 software (version 2.0.8, build 3, Qiagen). DNA methylation values obtained via pyrosequencing were compared between the HS and Control groups using a Wilcoxon test.

## Acknowledgments

We are grateful to the entire staff of the UE1295 PEAT (Nouzilly, France, doi: 10.15454/1.5572326250887292E12) for their excellent animal care. We are grateful to the genotoul bioinformatics platform Toulouse Occitanie (Bioinfo Genotoul, https://doi.org/10.15454/1.5572369328961167E12) for providing help, computing, and storage resources. RRBS sequencing was performed by the GenomEast platform, a member of the “France Génomique” consortium (ANR-10-INBS-0009). This work was funded by the French National Agency of Research (ChickStress project, ANR-13-ADAP). KK also thanks the French National Program MOPGA (Make Our Planet Great Again) for funding and support.

## Author contribution

FP, TZ and SLa conceived the experimental design and secured the funding. KK, JS and SLa performed the analyses. KK, CC, GD, SF, AH and JNH participated in the bioinformatic and statistical analyses. DG performed animal breeding. SLe performed molecular experiments. KK drafted the manuscript, FP, SLa and TZ revised the manuscript draft. All authors read and approved the final version.

## Data availability

The DNA methylation and RNA-seq datasets analyzed in the current study are available at ENA (https://www.ebi.ac.uk/ena/browser/home) with accession numbers PRJEB70935 and PRJEB28745, respectively.

## Competing interest**s**

The authors declare no competing interests.

## Notes

### Competing Interest Statement

The authors have declared no competing interest.

