## Supplementary Information for "Molecular responses of chicken embryos to maternal heat stress through DNA methylation and gene expression"

Fig S1: Ratio of read mapped on sex chromosomes to identify the sex of the embryos

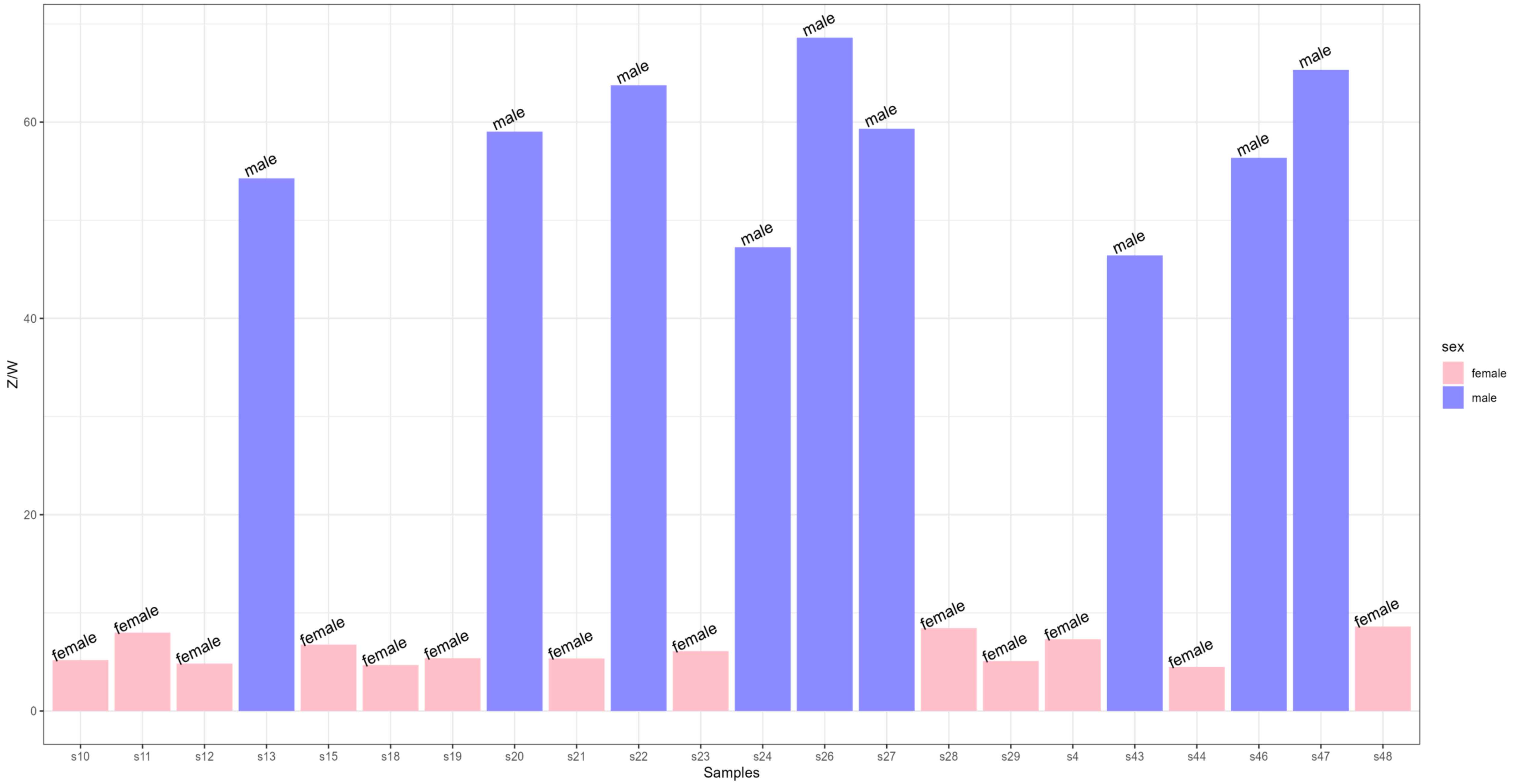

Fig S2: Distribution of DMCs on the chicken genome. A) Location of DMCs along chicken chromosomes. B) Number of hypo/hyper DMCs per chromosome

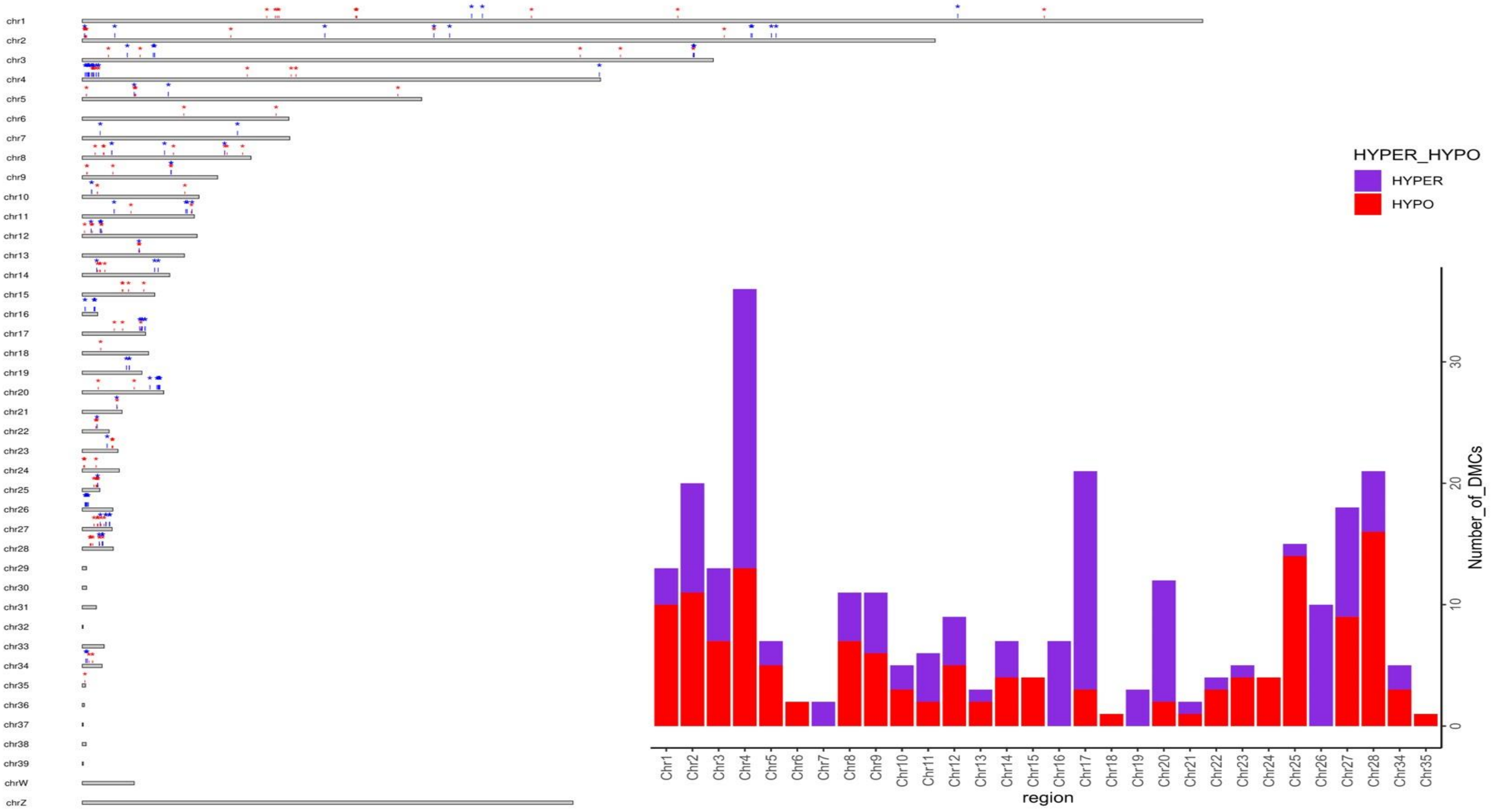

Fig S3: Methylation level obtained with RRBS and PyroMark results.

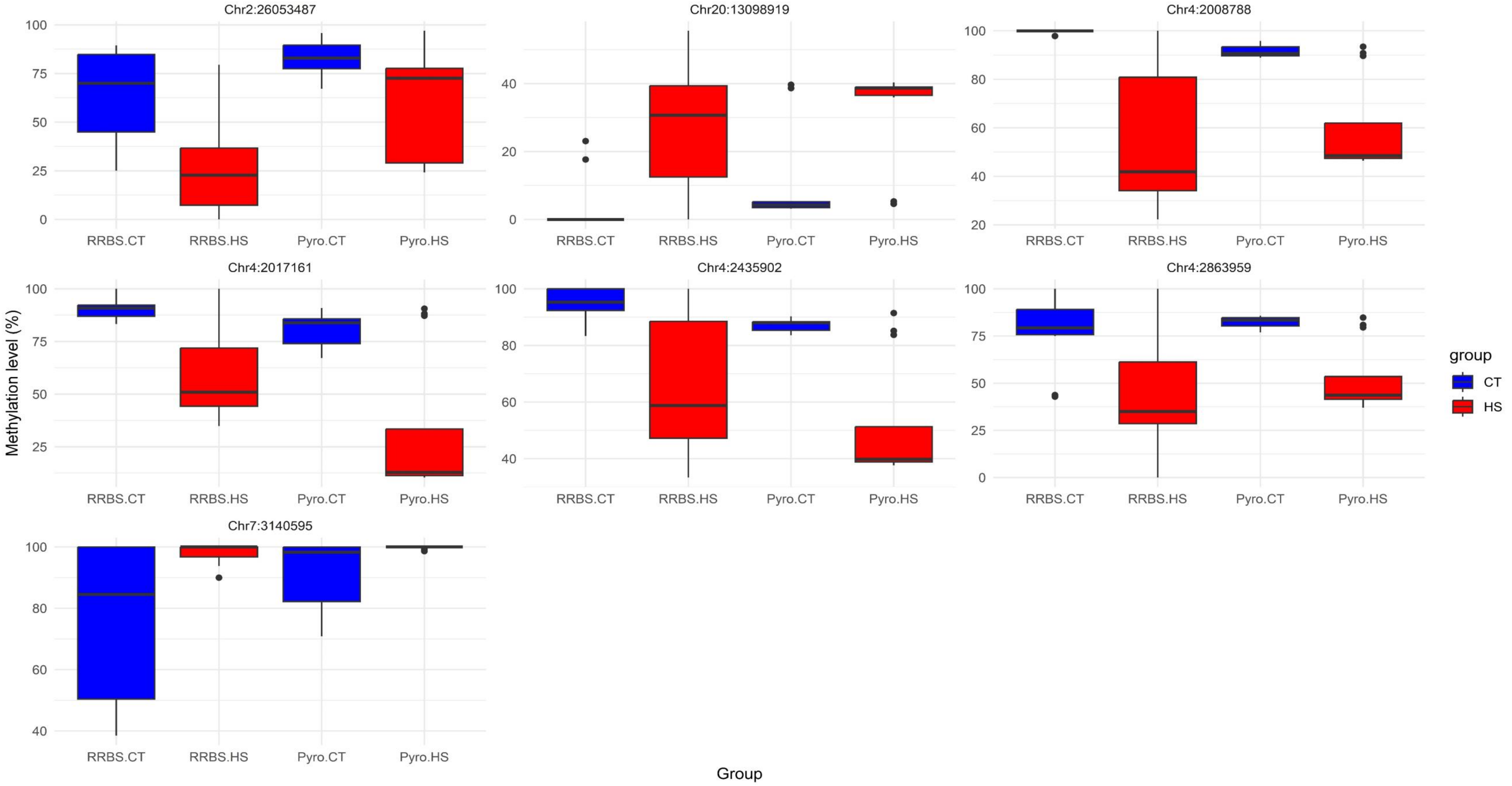

| position | Foward primer | Reverse primer | sequencing primer | PCR product length |
|---|---|---|---|---|
| Chr2:26053487 | TGGTGTTGAGAGTATTGAGTTATTT | [btn]ATCTTTCCCATACTAAATCCCCTATATCT | TGTTGTTTTGGTATATTAGATAATT | 171 |
| Chr4:2008788 | [btn]GTAGTTGGAGAGTTGTTAATTTAGTTT | CACAACAATATCTCCCAAATCTCAAT | CTTACCCACCATAATCTCCATA | 279 |
| Chr4:2017161 | GTTTTGTTTAAATTAAGGGGAAAGGA | [btn]CAATAAAACACCCAAAAACCATAATCACCA | AGTTTTGAGATTTTTTGAGTTAT | 143 |
| Chr4:2435902 | GGATAGGAAGAGATGGGAGTGTATA | [btn]CTCCCCACCAATACCAAAATACCTT | AGTAATTAGAGAGATTATTTTTAG | 246 |
| Chr4:2863959 | GGTAGGAGGTAGTAGATTAAGAATAGGA | [btn]CCATACCTTACAAATTCCTCTACATC | GAATTTGGAGTATGGGTT | 94 |
| Chr7:3140595 | TAAGGTGTTTGTAAAAGTGTTGAGGTATA | [btn]ATCCAATAAAAAAAAACCCATTTATCT | GTTGTTTATAGTTTGGAGATT | 158 |
| Chr20:13098919 | GAGTTGTTTTTTGGGAGTGATGTTT | [btn]AAATCCCTAAAACTCCTACACAACACAAT | GTGGAGGGAAAGGAT | 100 |

Table S1: Primers used for pyrosequencing analyses

Table S2: General statistics of RRBS sequencing results

| Sample name | Treatment | Number of reads | Aligned (%) | Number of mapped reads (million) | Number of CpGs* | coverage |
|---|---|---|---|---|---|---|
| s10 | HS | 17,731,302 | 64.2 | 11.4 | 1074875 | 18.15 |
| s11 | HS | 14,336,327 | 64.9 | 9.3 | 1073619 | 17.03 |
| s12 | HS | 37,767,056 | 65.2 | 24.6 | 1075224 | 21.80 |
| s13 | HS | 9,883,774 | 66.5 | 6.6 | 1066724 | 16.09 |
| s15 | HS | 22,808,286 | 64.2 | 14.6 | 1075083 | 19.63 |
| s18 | HS | 17,248,082 | 62.4 | 10.8 | 1074491 | 17.97 |
| s19 | HS | 23,054,605 | 64.3 | 14.8 | 1075024 | 18.15 |
| s20 | HS | 16,725,484 | 64.6 | 10.8 | 1074671 | 17.25 |
| s21 | HS | 19,725,892 | 65.4 | 12.9 | 1074940 | 18.54 |
| s22 | HS | 33,580,847 | 65.8 | 22.1 | 1075205 | 20.61 |
| s23 | HS | 16,816,217 | 65.3 | 11.0 | 1074585 | 17.36 |
| s24 | HS | 25,200,611 | 66.1 | 16.7 | 1075154 | 18.65 |
| s26 | CT | 13,323,097 | 65 | 8.6 | 1073549 | 17.22 |
| s27 | CT | 11,256,658 | 63.4 | 7.1 | 1071096 | 17.53 |
| s28 | CT | 17,761,202 | 66.6 | 11.8 | 1074713 | 17.59 |
| s29 | CT | 25,505,549 | 64.9 | 16.5 | 1075039 | 19.28 |
| s43 | CT | 15,721,809 | 65.7 | 10.3 | 1074843 | 17.77 |
| s44 | CT | 39,906,195 | 64 | 25.5 | 1075115 | 23.71 |
| s45 | CT | 10,052,977 | 63.9 | 6.4 | 1068642 | 15.97 |
| s46 | CT | 17,412,282 | 64.1 | 11.1 | 1074883 | 18.05 |
| s47 | CT | 11,240,033 | 63.7 | 7.2 | 1071413 | 16.52 |
| s48 | CT | 23,928,746 | 66.2 | 15.8 | 1075052 | 18.60 |
| Average | - | 20,044,865 | 64.84 | 13.00 | 1073815 | 18.34 |

*number of CpGs obtained after pre-procesing (with read depth > 5 and < 200)

Table S3: List of identified DMC and DMGs

| condition | seqnames | position | distance | feature_type | feature | tx_name | gene_name | gene_id |
|---|---|---|---|---|---|---|---|---|
| DMC | 1 | 32347715 | -55649 | other | NA | NA | NA | NA |
| DMC | 1 | 33831751 | 425 | introns | '828:XM_015281587:in | XM_015281587 | RASSF3 | LOC417828 |
| DMC | 1 | 33831751 | 425 | introns | '828:XM_015281591:in | XM_015281591 | RASSF3 | LOC417828 |
| DMC | 1 | 33831751 | 425 | introns | '828:XM_046938125:in | XM_046938125 | RASSF3 | LOC417828 |
| DMC | 1 | 33831751 | 425 | introns | '828:XM_040695649:in | XM_040695649 | RASSF3 | LOC417828 |
| DMC | 1 | 33831751 | 425 | introns | 09404:XR_005847925:i | XR_005847925 | LOC121109404 | LOC121109404 |
| DMC | 1 | 33831751 | 425 | intron1 | '828:XM_015281587:in | XM_015281587 | RASSF3 | LOC417828 |
| DMC | 1 | 33831751 | 425 | intron1 | 09404:XR_005847925:i | XR_005847925 | LOC121109404 | LOC121109404 |
| DMC | 1 | 34188568 | 1954 | introns | '833:XM_040695677:in | XM_040695677 | MSRB3 | LOC417833 |
| DMC | 1 | 34188568 | 1954 | introns | '833:XM_040695680:in | XM_040695680 | MSRB3 | LOC417833 |
| DMC | 1 | 34188568 | 1954 | introns | '833:XM_015281645:in | XM_015281645 | MSRB3 | LOC417833 |
| DMC | 1 | 34188568 | 1954 | introns | '833:XM_015281611:in | XM_015281611 | MSRB3 | LOC417833 |
| DMC | 1 | 34188568 | 1954 | introns | '833:XM_015281618:in | XM_015281618 | MSRB3 | LOC417833 |
| DMC | 1 | 34188568 | 1954 | introns | '833:XM_046938164:in | XM_046938164 | MSRB3 | LOC417833 |
| DMC | 1 | 34188568 | 1954 | introns | '833:XM_046938163:in | XM_046938163 | MSRB3 | LOC417833 |
| DMC | 1 | 34188568 | 1954 | introns | '833:NM_001199578:in | NM_001199578 | MSRB3 | LOC417833 |
| DMC | 1 | 34188568 | 1954 | introns | '833:XM_015281636:in | XM_015281636 | MSRB3 | LOC417833 |
| DMC | 1 | 34188568 | 1954 | introns | '833:XM_046938165:in | XM_046938165 | MSRB3 | LOC417833 |
| DMC | 1 | 34188568 | 1954 | introns | '833:XM_046938175:in | XM_046938175 | MSRB3 | LOC417833 |
| DMC | 1 | 34188568 | 1954 | introns | '833:XM_040695684:in | XM_040695684 | MSRB3 | LOC417833 |
| DMC | 1 | 34188568 | 1954 | introns | '833:XM_046938151:in | XM_046938151 | MSRB3 | LOC417833 |
| DMC | 1 | 34188568 | 1954 | introns | '833:XM_046938166:in | XM_046938166 | MSRB3 | LOC417833 |
| DMC | 1 | 34481401 | 1314 | introns | 00541:INRAGALT00000 | IRAGALT0000000109 | RAGALG0000000005 | IRAGALG0000000054 |
| DMC | 1 | 34481401 | 1314 | intron1 | 00541:INRAGALT00000 | IRAGALT0000000109 | RAGALG0000000005 | IRAGALG0000000054 |
| DMC | 1 | 47963035 | -124 | exons | 57684:XM_003640366 | XM_003640366 | C7orf49 | LOC100857684 |
| DMC | 1 | 47963035 | -124 | cds | 57684:XM_003640366 | XM_003640366 | C7orf49 | LOC100857684 |
| DMC | 1 | 48088881 | -651 | exons | 7879:NM_001142255:e | NM_001142255 | ART4 | LOC427879 |
| DMC | 1 | 48088881 | -651 | cds | 7879:NM_001142255:e | NM_001142255 | ART4 | LOC427879 |
| DMC | 1 | 48132522 | 1476 | promoter | 00773:INRAGALT00000 | IRAGALT0000000158 | RAGALG0000000007 | IRAGALG0000000077 |
| DMC | 1 | 48132522 | 1476 | introns | 00773:INRAGALT00000 | IRAGALT0000000158 | RAGALG0000000007 | IRAGALG0000000077 |
| DMC | 1 | 48132522 | 1476 | intron1 | 00773:INRAGALT00000 | IRAGALT0000000158 | RAGALG0000000007 | IRAGALG0000000077 |
| DMC | 1 | 68276036 | -2 | promoter | 3219:XM_040697354:pr | XM_040697354 | PPFIBP1 | LOC418219 |
| DMC | 1 | 68276036 | -2 | UTR5 | 19:XM_040697354:5'UT | XM_040697354 | PPFIBP1 | LOC418219 |
| DMC | 1 | 68276036 | -2 | exons | 3219:XM_040697354:ex | XM_040697354 | PPFIBP1 | LOC418219 |
| DMC | 1 | 68276036 | -2 | exon1 | 3219:XM_040697354:ex | XM_040697354 | PPFIBP1 | LOC418219 |
| DMC | 1 | 68276036 | -2 | introns | 219:XM_015290411:int | XM_015290411 | PPFIBP1 | LOC418219 |
| DMC | 1 | 68276036 | -2 | introns | 219:XM_015290418:int | XM_015290418 | PPFIBP1 | LOC418219 |
| DMC | 1 | 68276036 | -2 | introns | 219:XM_025148974:int | XM_025148974 | PPFIBP1 | LOC418219 |
| DMC | 1 | 68276036 | -2 | introns | 219:XM_025148981:int | XM_025148981 | PPFIBP1 | LOC418219 |
| DMC | 1 | 68276036 | -2 | introns | 219:XM_025148984:int | XM_025148984 | PPFIBP1 | LOC418219 |
| DMC | 1 | 68276036 | -2 | introns | 219:XM_025148986:int | XM_025148986 | PPFIBP1 | LOC418219 |
| DMC | 1 | 68276036 | -2 | introns | 219:XM_025148991:int | XM_025148991 | PPFIBP1 | LOC418219 |
| DMC | 1 | 68276036 | -2 | introns | 219:XM_025148992:int | XM_025148992 | PPFIBP1 | LOC418219 |
| DMC | 1 | 68276036 | -2 | introns | 219:XM_025148994:int | XM_025148994 | PPFIBP1 | LOC418219 |
| DMC | 1 | 68276036 | -2 | introns | 219:XM_025148995:int | XM_025148995 | PPFIBP1 | LOC418219 |
| DMC | 1 | 68276036 | -2 | introns | 219:XM_025148997:int | XM_025148997 | PPFIBP1 | LOC418219 |
| DMC | 1 | 68276036 | -2 | introns | 219:XM_025148998:int | XM_025148998 | PPFIBP1 | LOC418219 |
| DMC | 1 | 68276036 | -2 | introns | 219:XM_025149012:int | XM_025149012 | PPFIBP1 | LOC418219 |
| DMC | 1 | 68276036 | -2 | introns | 219:XM_025149025:int | XM_025149025 | PPFIBP1 | LOC418219 |
| DMC | 1 | 68276036 | -2 | introns | 219:XM_040697345:int | XM_040697345 | PPFIBP1 | LOC418219 |
| DMC | 1 | 68276036 | -2 | introns | 219:XM_040697364:int | XM_040697364 | PPFIBP1 | LOC418219 |
| DMC | 1 | 68276036 | -2 | introns | 219:XM_040697375:int | XM_040697375 | PPFIBP1 | LOC418219 |
| DMC | 1 | 68276036 | -2 | introns | 219:XM_040697389:int | XM_040697389 | PPFIBP1 | LOC418219 |
| DMC | 1 | 68276036 | -2 | introns | 219:XM_040697415:int | XM_040697415 | PPFIBP1 | LOC418219 |
| DMC | 1 | 68276036 | -2 | introns | 219:XM_040697425:int | XM_040697425 | PPFIBP1 | LOC418219 |
| DMC | 1 | 68276036 | -2 | introns | 219:XM_040697432:int | XM_040697432 | PPFIBP1 | LOC418219 |
| DMC | 1 | 68276036 | -2 | introns | 219:XM_040697442:int | XM_040697442 | PPFIBP1 | LOC418219 |
| DMC | 1 | 68276036 | -2 | introns | 219:XM_040697450:int | XM_040697450 | PPFIBP1 | LOC418219 |
| DMC | 1 | 68276036 | -2 | introns | 219:XM_040697451:int | XM_040697451 | PPFIBP1 | LOC418219 |
| DMC | 1 | 68276036 | -2 | introns | 219:XM_040697455:int | XM_040697455 | PPFIBP1 | LOC418219 |
| DMC | 1 | 68276036 | -2 | introns | 219:XM_040697466:int | XM_040697466 | PPFIBP1 | LOC418219 |
| DMC | 1 | 68276036 | -2 | introns | 219:XM_040697477:int | XM_040697477 | PPFIBP1 | LOC418219 |
| DMC | 1 | 68276036 | -2 | introns | 219:XM_040697487:int | XM_040697487 | PPFIBP1 | LOC418219 |
| DMC | 1 | 68276036 | -2 | introns | 219:XM_040697503:int | XM_040697503 | PPFIBP1 | LOC418219 |
| DMC | 1 | 68276036 | -2 | introns | 219:XM_040697516:int | XM_040697516 | PPFIBP1 | LOC418219 |
| DMC | 1 | 68276036 | -2 | introns | 219:XM_040697521:int | XM_040697521 | PPFIBP1 | LOC418219 |

| | | | | | | | | |
|---|---|---|---|---|---|---|---|---|
| DMC | 1 | 68276036 | -2 | introns | 219:XM_040697539:int | XM_040697539 | PPFIBP1 | LOC418219 |
| DMC | 1 | 68276036 | -2 | introns | 219:XM_046939437:int | XM_046939437 | PPFIBP1 | LOC418219 |
| DMC | 1 | 68276036 | -2 | introns | 219:XM_046939448:int | XM_046939448 | PPFIBP1 | LOC418219 |
| DMC | 1 | 68276036 | -2 | introns | 219:XM_046939459:int | XM_046939459 | PPFIBP1 | LOC418219 |
| DMC | 1 | 68276036 | -2 | introns | 219:XM_046939469:int | XM_046939469 | PPFIBP1 | LOC418219 |
| DMC | 1 | 68276036 | -2 | introns | 219:XM_046939470:int | XM_046939470 | PPFIBP1 | LOC418219 |
| DMC | 1 | 68276036 | -2 | introns | 219:XM_046939471:int | XM_046939471 | PPFIBP1 | LOC418219 |
| DMC | 1 | 68276036 | -2 | introns | 219:XM_046939495:int | XM_046939495 | PPFIBP1 | LOC418219 |
| DMC | 1 | 68276036 | -2 | intron1 | 219:XM_015290411:int | XM_015290411 | PPFIBP1 | LOC418219 |
| DMC | 1 | 68276036 | -2 | intron1 | 219:XM_015290418:int | XM_015290418 | PPFIBP1 | LOC418219 |
| DMC | 1 | 68276036 | -2 | intron1 | 219:XM_025148974:int | XM_025148974 | PPFIBP1 | LOC418219 |
| DMC | 1 | 68276036 | -2 | intron1 | 219:XM_025148981:int | XM_025148981 | PPFIBP1 | LOC418219 |
| DMC | 1 | 68276036 | -2 | intron1 | 219:XM_025148984:int | XM_025148984 | PPFIBP1 | LOC418219 |
| DMC | 1 | 68276036 | -2 | intron1 | 219:XM_025148986:int | XM_025148986 | PPFIBP1 | LOC418219 |
| DMC | 1 | 68276036 | -2 | intron1 | 219:XM_025148991:int | XM_025148991 | PPFIBP1 | LOC418219 |
| DMC | 1 | 68276036 | -2 | intron1 | 219:XM_025148992:int | XM_025148992 | PPFIBP1 | LOC418219 |
| DMC | 1 | 68276036 | -2 | intron1 | 219:XM_025148994:int | XM_025148994 | PPFIBP1 | LOC418219 |
| DMC | 1 | 68276036 | -2 | intron1 | 219:XM_025148995:int | XM_025148995 | PPFIBP1 | LOC418219 |
| DMC | 1 | 68276036 | -2 | intron1 | 219:XM_025148997:int | XM_025148997 | PPFIBP1 | LOC418219 |
| DMC | 1 | 68276036 | -2 | intron1 | 219:XM_025148998:int | XM_025148998 | PPFIBP1 | LOC418219 |
| DMC | 1 | 68276036 | -2 | intron1 | 219:XM_025149012:int | XM_025149012 | PPFIBP1 | LOC418219 |
| DMC | 1 | 68276036 | -2 | intron1 | 219:XM_025149025:int | XM_025149025 | PPFIBP1 | LOC418219 |
| DMC | 1 | 68276036 | -2 | intron1 | 219:XM_040697345:int | XM_040697345 | PPFIBP1 | LOC418219 |
| DMC | 1 | 68276036 | -2 | intron1 | 219:XM_040697364:int | XM_040697364 | PPFIBP1 | LOC418219 |
| DMC | 1 | 68276036 | -2 | intron1 | 219:XM_040697375:int | XM_040697375 | PPFIBP1 | LOC418219 |
| DMC | 1 | 68276036 | -2 | intron1 | 219:XM_040697389:int | XM_040697389 | PPFIBP1 | LOC418219 |
| DMC | 1 | 68276036 | -2 | intron1 | 219:XM_040697415:int | XM_040697415 | PPFIBP1 | LOC418219 |
| DMC | 1 | 68276036 | -2 | intron1 | 219:XM_040697425:int | XM_040697425 | PPFIBP1 | LOC418219 |
| DMC | 1 | 68276036 | -2 | intron1 | 219:XM_040697432:int | XM_040697432 | PPFIBP1 | LOC418219 |
| DMC | 1 | 68276036 | -2 | intron1 | 219:XM_040697442:int | XM_040697442 | PPFIBP1 | LOC418219 |
| DMC | 1 | 68276036 | -2 | intron1 | 219:XM_040697450:int | XM_040697450 | PPFIBP1 | LOC418219 |
| DMC | 1 | 68276036 | -2 | intron1 | 219:XM_040697451:int | XM_040697451 | PPFIBP1 | LOC418219 |
| DMC | 1 | 68276036 | -2 | intron1 | 219:XM_040697455:int | XM_040697455 | PPFIBP1 | LOC418219 |
| DMC | 1 | 68276036 | -2 | intron1 | 219:XM_040697466:int | XM_040697466 | PPFIBP1 | LOC418219 |
| DMC | 1 | 68276036 | -2 | intron1 | 219:XM_040697477:int | XM_040697477 | PPFIBP1 | LOC418219 |
| DMC | 1 | 68276036 | -2 | intron1 | 219:XM_040697487:int | XM_040697487 | PPFIBP1 | LOC418219 |
| DMC | 1 | 68276036 | -2 | intron1 | 219:XM_040697503:int | XM_040697503 | PPFIBP1 | LOC418219 |
| DMC | 1 | 68276036 | -2 | intron1 | 219:XM_040697516:int | XM_040697516 | PPFIBP1 | LOC418219 |
| DMC | 1 | 68276036 | -2 | intron1 | 219:XM_040697521:int | XM_040697521 | PPFIBP1 | LOC418219 |
| DMC | 1 | 68276036 | -2 | intron1 | 219:XM_040697539:int | XM_040697539 | PPFIBP1 | LOC418219 |
| DMC | 1 | 68276036 | -2 | intron1 | 219:XM_046939437:int | XM_046939437 | PPFIBP1 | LOC418219 |
| DMC | 1 | 68276036 | -2 | intron1 | 219:XM_046939448:int | XM_046939448 | PPFIBP1 | LOC418219 |
| DMC | 1 | 68276036 | -2 | intron1 | 219:XM_046939459:int | XM_046939459 | PPFIBP1 | LOC418219 |
| DMC | 1 | 68276036 | -2 | intron1 | 219:XM_046939469:int | XM_046939469 | PPFIBP1 | LOC418219 |
| DMC | 1 | 68276036 | -2 | intron1 | 219:XM_046939470:int | XM_046939470 | PPFIBP1 | LOC418219 |
| DMC | 1 | 68276036 | -2 | intron1 | 219:XM_046939471:int | XM_046939471 | PPFIBP1 | LOC418219 |
| DMC | 1 | 68276036 | -2 | intron1 | 219:XM_046939495:int | XM_046939495 | PPFIBP1 | LOC418219 |
| DMC | 1 | 68276036 | -2 | downstream | 0394:TAGAGALT00000( | GAGALT0000000222 | GAGALG000000060 | GAGALG0000000603 |
| DMC | 1 | 70154145 | 1092 | promoter | 07132:DAVISGALT0007 | AVISGALT00071320 | AVISGALG0000071 | AVISGALG00000713 |
| DMC | 1 | 70154145 | 1092 | promoter | 07133:DAVISGALT0007 | AVISGALT00071330 | AVISGALG0000071 | AVISGALG00000713 |
| DMC | 1 | 70154145 | 1092 | introns | 935:XM_015291043:int | XM_015291043 | PHF21B | LOC427935 |
| DMC | 1 | 70154145 | 1092 | introns | 935:XM_040703199:int | XM_040703199 | PHF21B | LOC427935 |
| DMC | 1 | 70154145 | 1092 | introns | 935:XM_046943894:int | XM_046943894 | PHF21B | LOC427935 |
| DMC | 1 | 78759889 | 3868 | introns | 07536:NONGGAT0127 | NONGGAT012750 | NONGGAG007536 | NONGGAG007536 |
| DMC | 1 | 104414426 | 267 | promoter | 05086:TAGAGALT0000( | GAGALT000000347 | GAGALG000000005 | GAGALG0000000050 |
| DMC | 1 | 104414426 | 267 | promoter | 05086:TAGAGALT0000( | GAGALT000000347 | GAGALG000000005 | GAGALG0000000050 |
| DMC | 1 | 104414426 | 267 | promoter | 08807:DAVISGALT0008 | AVISGALT00088070 | AVISGALG0000088 | AVISGALG00000880 |
| DMC | 1 | 104414426 | 267 | exons | 008807:DAVISGALT000 | AVISGALT00088070 | AVISGALG0000088 | AVISGALG00000880 |
| DMC | 1 | 104414426 | 267 | exon1 | 008807:DAVISGALT000 | AVISGALT00088070 | AVISGALG0000088 | AVISGALG00000880 |
| DMC | 1 | 104414426 | 267 | introns | 489:XM_015299614:int | XM_015299614 | GRIK1 | LOC418489 |
| DMC | 1 | 104414426 | 267 | introns | 489:XM_015299627:int | XM_015299627 | GRIK1 | LOC418489 |
| DMC | 1 | 104414426 | 267 | introns | 489:XM_040646819:int | XM_040646819 | GRIK1 | LOC418489 |
| DMC | 1 | 104414426 | 267 | introns | 489:XM_040646828:int | XM_040646828 | GRIK1 | LOC418489 |
| DMC | 1 | 104414426 | 267 | introns | 489:XM_040646808:int | XM_040646808 | GRIK1 | LOC418489 |
| DMC | 1 | 104414426 | 267 | introns | 489:XM_040646836:int | XM_040646836 | GRIK1 | LOC418489 |
| DMC | 1 | 104414426 | 267 | introns | 489:XM_040646841:int | XM_040646841 | GRIK1 | LOC418489 |
| DMC | 1 | 104414426 | 267 | introns | 489:XM_040646844:int | XM_040646844 | GRIK1 | LOC418489 |

| | | | | | | | | |
|---|---|---|---|---|---|---|---|---|
| DMC | 1 | 104414426 | 267 | intron1 | 489:XM_015299614:int | XM_015299614 | GRIK1 | LOC418489 |
| DMC | 1 | 104414426 | 267 | intron1 | 489:XM_015299627:int | XM_015299627 | GRIK1 | LOC418489 |
| DMC | 1 | 104414426 | 267 | intron1 | 489:XM_040646819:int | XM_040646819 | GRIK1 | LOC418489 |
| DMC | 1 | 104414426 | 267 | intron1 | 489:XM_040646828:int | XM_040646828 | GRIK1 | LOC418489 |
| DMC | 1 | 104414426 | 267 | intron1 | 489:XM_040646808:int | XM_040646808 | GRIK1 | LOC418489 |
| DMC | 1 | 104414426 | 267 | intron1 | 489:XM_040646836:int | XM_040646836 | GRIK1 | LOC418489 |
| DMC | 1 | 104414426 | 267 | intron1 | 489:XM_040646841:int | XM_040646841 | GRIK1 | LOC418489 |
| DMC | 1 | 104414426 | 267 | intron1 | 489:XM_040646844:int | XM_040646844 | GRIK1 | LOC418489 |
| DMC | 1 | 153503892 | 1889 | introns | 01062:ENSGALT00010( | NSGALT0001000233 | NSGALG000100010 | NSGALG0001000106 |
| DMC | 1 | 153503892 | 1889 | introns | 01062:ENSGALT00010( | NSGALT0001000233 | NSGALG000100010 | NSGALG0001000106 |
| DMC | 1 | 153503892 | 1889 | introns | 01062:ENSGALT00010( | NSGALT0001000233 | NSGALG000100010 | NSGALG0001000106 |
| DMC | 1 | 153503892 | 1889 | introns | 01064:ENSGALT00010( | NSGALT0001000234 | NSGALG000100010 | NSGALG0001000106 |
| DMC | 1 | 153503892 | 1889 | introns | 01064:ENSGALT00010( | NSGALT0001000234 | NSGALG000100010 | NSGALG0001000106 |
| DMC | 1 | 153503892 | 1889 | introns | 01064:ENSGALT00010( | NSGALT0001000234 | NSGALG000100010 | NSGALG0001000106 |
| DMC | 1 | 153503892 | 1889 | introns | 01064:ENSGALT00010( | NSGALT0001000234 | NSGALG000100010 | NSGALG0001000106 |
| DMC | 1 | 153503892 | 1889 | intron1 | 01062:ENSGALT00010( | NSGALT0001000233 | NSGALG000100010 | NSGALG0001000106 |
| DMC | 1 | 153503892 | 1889 | intron1 | 01064:ENSGALT00010( | NSGALT0001000234 | NSGALG000100010 | NSGALG0001000106 |
| DMC | 1 | 153503892 | 1889 | intron1 | 01064:ENSGALT00010( | NSGALT0001000234 | NSGALG000100010 | NSGALG0001000106 |
| DMC | 1 | 168677980 | -388 | promoter | 11610:DAVISGALT0011 | AVISGALT00116100 | AVISGALG0000116 | AVISGALG00001161 |
| DMC | 1 | 168677980 | -388 | exons | 96118:NM_205199:exc | NM_205199 | LPAR6 | LOC396118 |
| DMC | 1 | 168677980 | -388 | exon1 | 96118:NM_205199:exc | NM_205199 | LPAR6 | LOC396118 |
| DMC | 1 | 168677980 | -388 | introns | 6582:NM_204419:intro | NM_204419 | RB1 | LOC386582 |
| DMC | 1 | 168677980 | -388 | introns | 82:XM_046898854:intr | XM_046898854 | RB1 | LOC386582 |
| DMC | 1 | 168677980 | -388 | cds | 96118:NM_205199:exc | NM_205199 | LPAR6 | LOC396118 |
| DMC | 2 | 363130 | 1064 | introns | 8122:XM_015281150:i | XM_015281150 | KCNH2 | LOC100858122 |
| DMC | 2 | 383062 | -52 | exons | 8122:XM_015281150:e | XM_015281150 | KCNH2 | LOC100858122 |
| DMC | 2 | 383062 | -52 | UTR3 | 122:XM_015281150:3'U | XM_015281150 | KCNH2 | LOC100858122 |
| DMC | 2 | 383062 | -52 | downstream | 3159:XM_015281153:d | XM_015281153 | AOC1 | LOC100858159 |
| DMC | 2 | 398203 | -157 | promoter | 58196:XM_040696524: | XM_040696524 | LOC100858196 | LOC100858196 |
| DMC | 2 | 398203 | -157 | promoter | 58196:XM_025147395: | XM_025147395 | LOC100858196 | LOC100858196 |
| DMC | 2 | 398203 | -157 | exons | .13126:XM_040696529 | XM_040696529 | LOC121113126 | LOC121113126 |
| DMC | 2 | 398203 | -157 | exons | .13126:XM_040696531 | XM_040696531 | LOC121113126 | LOC121113126 |
| DMC | 2 | 398203 | -157 | exons | .13126:XM_046927919 | XM_046927919 | LOC121113126 | LOC121113126 |
| DMC | 2 | 398203 | -157 | exons | .13126:XM_046927921 | XM_046927921 | LOC121113126 | LOC121113126 |
| DMC | 2 | 398203 | -157 | exons | .13126:XM_040696528 | XM_040696528 | LOC121113126 | LOC121113126 |
| DMC | 2 | 398203 | -157 | exons | .13126:XM_040696530 | XM_040696530 | LOC121113126 | LOC121113126 |
| DMC | 2 | 398203 | -157 | introns | 06156:INRAGALT00000 | IRAGALT000000131 | RAGALG00000006 | RAGALG000000061 |
| DMC | 2 | 398203 | -157 | intron1 | 06156:INRAGALT00000 | IRAGALT000000131 | RAGALG00000006 | RAGALG000000061 |
| DMC | 2 | 398203 | -157 | cds | .13126:XM_040696529 | XM_040696529 | LOC121113126 | LOC121113126 |
| DMC | 2 | 398203 | -157 | cds | .13126:XM_040696531 | XM_040696531 | LOC121113126 | LOC121113126 |
| DMC | 2 | 398203 | -157 | cds | .13126:XM_046927919 | XM_046927919 | LOC121113126 | LOC121113126 |
| DMC | 2 | 398203 | -157 | cds | .13126:XM_046927921 | XM_046927921 | LOC121113126 | LOC121113126 |
| DMC | 2 | 398203 | -157 | cds | .13126:XM_040696528 | XM_040696528 | LOC121113126 | LOC121113126 |
| DMC | 2 | 398203 | -157 | cds | .13126:XM_040696530 | XM_040696530 | LOC121113126 | LOC121113126 |
| DMC | 2 | 398203 | -157 | downstream | 1142:TAGAGALT00000( | GAGALT000000153 | GAGALG00000003 | GAGALG0000000311 |
| DMC | 2 | 398252 | -206 | promoter | 58196:XM_040696524: | XM_040696524 | LOC100858196 | LOC100858196 |
| DMC | 2 | 398252 | -206 | promoter | 58196:XM_025147395: | XM_025147395 | LOC100858196 | LOC100858196 |
| DMC | 2 | 398252 | -206 | exons | .13126:XM_040696529 | XM_040696529 | LOC121113126 | LOC121113126 |
| DMC | 2 | 398252 | -206 | exons | .13126:XM_040696531 | XM_040696531 | LOC121113126 | LOC121113126 |
| DMC | 2 | 398252 | -206 | exons | .13126:XM_046927919 | XM_046927919 | LOC121113126 | LOC121113126 |
| DMC | 2 | 398252 | -206 | exons | .13126:XM_046927921 | XM_046927921 | LOC121113126 | LOC121113126 |
| DMC | 2 | 398252 | -206 | exons | .13126:XM_040696528 | XM_040696528 | LOC121113126 | LOC121113126 |
| DMC | 2 | 398252 | -206 | exons | .13126:XM_040696530 | XM_040696530 | LOC121113126 | LOC121113126 |
| DMC | 2 | 398252 | -206 | introns | 06156:INRAGALT00000 | IRAGALT000000131 | RAGALG00000006 | RAGALG000000061 |
| DMC | 2 | 398252 | -206 | intron1 | 06156:INRAGALT00000 | IRAGALT000000131 | RAGALG00000006 | RAGALG000000061 |
| DMC | 2 | 398252 | -206 | cds | .13126:XM_040696529 | XM_040696529 | LOC121113126 | LOC121113126 |
| DMC | 2 | 398252 | -206 | cds | .13126:XM_040696531 | XM_040696531 | LOC121113126 | LOC121113126 |
| DMC | 2 | 398252 | -206 | cds | .13126:XM_046927919 | XM_046927919 | LOC121113126 | LOC121113126 |
| DMC | 2 | 398252 | -206 | cds | .13126:XM_046927921 | XM_046927921 | LOC121113126 | LOC121113126 |
| DMC | 2 | 398252 | -206 | cds | .13126:XM_040696528 | XM_040696528 | LOC121113126 | LOC121113126 |
| DMC | 2 | 398252 | -206 | cds | .13126:XM_040696530 | XM_040696530 | LOC121113126 | LOC121113126 |
| DMC | 2 | 398252 | -206 | downstream | 1142:TAGAGALT00000( | GAGALT000000153 | GAGALG00000003 | GAGALG0000000311 |
| DMC | 2 | 529220 | 370 | introns | 51647:XM_025147396: | XM_025147396 | GIMAP7L5 | LOC101751647 |
| DMC | 2 | 529220 | 370 | downstream | 7409:ENSGALT0001006 | NSGALT0001006645 | NSGALG000100274 | NSGALG0001002740 |
| DMC | 2 | 529220 | 370 | downstream | 7409:ENSGALT0001006 | NSGALT0001006645 | NSGALG000100274 | NSGALG0001002740 |
| DMC | 2 | 529220 | 370 | downstream | 7409:ENSGALT0001006 | NSGALT0001006646 | NSGALG000100274 | NSGALG0001002740 |
| DMC | 2 | 529220 | 370 | downstream | 7409:ENSGALT0001006 | NSGALT0001006646 | NSGALG000100274 | NSGALG0001002740 |

| | | | | | | | | |
|---|---|---|---|---|---|---|---|---|
| DMC | 2 | 529220 | 370 | downstream | 7409:ENSGALT0001006 | NSGALT0001006646 | NSGALG000100274 | NSGALG0001002740 |
| DMC | 2 | 529220 | 370 | downstream | 7409:ENSGALT0001006 | NSGALT0001006646 | NSGALG000100274 | NSGALG0001002740 |
| DMC | 2 | 529220 | 370 | downstream | 7409:ENSGALT0001006 | NSGALT0001006646 | NSGALG000100274 | NSGALG0001002740 |
| DMC | 2 | 529220 | 370 | downstream | 3436:XM_004939058:d | XM_004939058 | LOC101748436 | LOC101748436 |
| DMC | 2 | 529220 | 370 | downstream | 3436:XM_025147387:d | XM_025147387 | LOC101748436 | LOC101748436 |
| DMC | 2 | 529220 | 370 | downstream | 3436:XM_040696522:d | XM_040696522 | LOC101748436 | LOC101748436 |
| DMC | 2 | 593744 | 542 | promoter | 52490:XM_015281180: | XM_015281180 | LOC107052490 | LOC107052490 |
| DMC | 2 | 593744 | 542 | introns | 51647:XM_025147396: | XM_025147396 | GIMAP7L5 | LOC101751647 |
| DMC | 2 | 593744 | 542 | introns | 47780:XM_046938094:i | XM_046938094 | LOC101747780 | LOC101747780 |
| DMC | 2 | 610965 | -909 | promoter | 0374:XM_001231409:pr | XM_001231409 | LOC420374 | LOC420374 |
| DMC | 2 | 610965 | -909 | introns | 51647:XM_025147396: | XM_025147396 | GIMAP7L5 | LOC101751647 |
| DMC | 2 | 610965 | -909 | downstream | 7780:XM_046938094:d | XM_046938094 | LOC101747780 | LOC101747780 |
| DMC | 2 | 636466 | 322 | exons | 027532:ENSGALT00010 | NSGALT0001006668 | NSGALG000100275 | NSGALG0001002753 |
| DMC | 2 | 636466 | 322 | introns | 51647:XM_025147396: | XM_025147396 | GIMAP7L5 | LOC101751647 |
| DMC | 2 | 636466 | 322 | downstream | 7532:ENSGALT0001006 | NSGALT0001006668 | NSGALG000100275 | NSGALG0001002753 |
| DMC | 2 | 759278 | -8 | exons | 097:XM_015281158:ex | XM_015281158 | MAP4 | LOC396097 |
| DMC | 2 | 759278 | -8 | exons | 097:XM_015281160:ex | XM_015281160 | MAP4 | LOC396097 |
| DMC | 2 | 759278 | -8 | exons | 097:XM_015281167:ex | XM_015281167 | MAP4 | LOC396097 |
| DMC | 2 | 759278 | -8 | exons | 097:XM_040696468:ex | XM_040696468 | MAP4 | LOC396097 |
| DMC | 2 | 759278 | -8 | exons | 097:XM_040696469:ex | XM_040696469 | MAP4 | LOC396097 |
| DMC | 2 | 759278 | -8 | exons | 097:XM_040696470:ex | XM_040696470 | MAP4 | LOC396097 |
| DMC | 2 | 759278 | -8 | exons | 097:XM_040696472:ex | XM_040696472 | MAP4 | LOC396097 |
| DMC | 2 | 759278 | -8 | exons | 097:XM_040696473:ex | XM_040696473 | MAP4 | LOC396097 |
| DMC | 2 | 759278 | -8 | exons | 097:XM_040696475:ex | XM_040696475 | MAP4 | LOC396097 |
| DMC | 2 | 759278 | -8 | exons | 097:XM_040696477:ex | XM_040696477 | MAP4 | LOC396097 |
| DMC | 2 | 759278 | -8 | exons | 097:XM_040696478:ex | XM_040696478 | MAP4 | LOC396097 |
| DMC | 2 | 759278 | -8 | exons | 097:XM_040696479:ex | XM_040696479 | MAP4 | LOC396097 |
| DMC | 2 | 759278 | -8 | exons | 097:XM_040696480:ex | XM_040696480 | MAP4 | LOC396097 |
| DMC | 2 | 759278 | -8 | exons | 097:XM_040696481:ex | XM_040696481 | MAP4 | LOC396097 |
| DMC | 2 | 759278 | -8 | exons | 097:XM_040696482:ex | XM_040696482 | MAP4 | LOC396097 |
| DMC | 2 | 759278 | -8 | exons | 097:XM_040696483:ex | XM_040696483 | MAP4 | LOC396097 |
| DMC | 2 | 759278 | -8 | exons | 097:XM_040696485:ex | XM_040696485 | MAP4 | LOC396097 |
| DMC | 2 | 759278 | -8 | exons | 097:XM_040696486:ex | XM_040696486 | MAP4 | LOC396097 |
| DMC | 2 | 759278 | -8 | exons | 097:XM_040696487:ex | XM_040696487 | MAP4 | LOC396097 |
| DMC | 2 | 759278 | -8 | introns | 51647:XM_025147396: | XM_025147396 | GIMAP7L5 | LOC101751647 |
| DMC | 2 | 759278 | -8 | UTR3 | 07:XM_015281158:3'UT | XM_015281158 | MAP4 | LOC396097 |
| DMC | 2 | 759278 | -8 | UTR3 | 07:XM_015281160:3'UT | XM_015281160 | MAP4 | LOC396097 |
| DMC | 2 | 759278 | -8 | UTR3 | 07:XM_015281167:3'UT | XM_015281167 | MAP4 | LOC396097 |
| DMC | 2 | 759278 | -8 | UTR3 | 07:XM_040696468:3'UT | XM_040696468 | MAP4 | LOC396097 |
| DMC | 2 | 759278 | -8 | UTR3 | 07:XM_040696469:3'UT | XM_040696469 | MAP4 | LOC396097 |
| DMC | 2 | 759278 | -8 | UTR3 | 07:XM_040696470:3'UT | XM_040696470 | MAP4 | LOC396097 |
| DMC | 2 | 759278 | -8 | UTR3 | 07:XM_040696472:3'UT | XM_040696472 | MAP4 | LOC396097 |
| DMC | 2 | 759278 | -8 | UTR3 | 07:XM_040696473:3'UT | XM_040696473 | MAP4 | LOC396097 |
| DMC | 2 | 759278 | -8 | UTR3 | 07:XM_040696475:3'UT | XM_040696475 | MAP4 | LOC396097 |
| DMC | 2 | 759278 | -8 | UTR3 | 07:XM_040696477:3'UT | XM_040696477 | MAP4 | LOC396097 |
| DMC | 2 | 759278 | -8 | UTR3 | 07:XM_040696478:3'UT | XM_040696478 | MAP4 | LOC396097 |
| DMC | 2 | 759278 | -8 | UTR3 | 07:XM_040696479:3'UT | XM_040696479 | MAP4 | LOC396097 |
| DMC | 2 | 759278 | -8 | UTR3 | 07:XM_040696480:3'UT | XM_040696480 | MAP4 | LOC396097 |
| DMC | 2 | 759278 | -8 | UTR3 | 07:XM_040696481:3'UT | XM_040696481 | MAP4 | LOC396097 |
| DMC | 2 | 759278 | -8 | UTR3 | 07:XM_040696482:3'UT | XM_040696482 | MAP4 | LOC396097 |
| DMC | 2 | 759278 | -8 | UTR3 | 07:XM_040696483:3'UT | XM_040696483 | MAP4 | LOC396097 |
| DMC | 2 | 759278 | -8 | UTR3 | 07:XM_040696485:3'UT | XM_040696485 | MAP4 | LOC396097 |
| DMC | 2 | 759278 | -8 | UTR3 | 07:XM_040696486:3'UT | XM_040696486 | MAP4 | LOC396097 |
| DMC | 2 | 759278 | -8 | UTR3 | 07:XM_040696487:3'UT | XM_040696487 | MAP4 | LOC396097 |
| DMC | 2 | 759278 | -8 | downstream | 97:XM_015281157:dov | XM_015281157 | MAP4 | LOC396097 |
| DMC | 2 | 759278 | -8 | downstream | 97:XM_015281162:dov | XM_015281162 | MAP4 | LOC396097 |
| DMC | 2 | 759278 | -8 | downstream | 97:XM_015281163:dov | XM_015281163 | MAP4 | LOC396097 |
| DMC | 2 | 759278 | -8 | downstream | 97:XM_015281164:dov | XM_015281164 | MAP4 | LOC396097 |
| DMC | 2 | 759278 | -8 | downstream | 97:XM_015281165:dov | XM_015281165 | MAP4 | LOC396097 |
| DMC | 2 | 759278 | -8 | downstream | 97:XM_015281166:dov | XM_015281166 | MAP4 | LOC396097 |
| DMC | 2 | 759278 | -8 | downstream | 97:XM_015281168:dov | XM_015281168 | MAP4 | LOC396097 |
| DMC | 2 | 759278 | -8 | downstream | 97:XM_025147368:dov | XM_025147368 | MAP4 | LOC396097 |
| DMC | 2 | 759278 | -8 | downstream | 97:XM_040696471:dov | XM_040696471 | MAP4 | LOC396097 |
| DMC | 2 | 759278 | -8 | downstream | 97:XM_040696476:dov | XM_040696476 | MAP4 | LOC396097 |
| DMC | 2 | 759278 | -8 | downstream | 97:XM_046927630:dov | XM_046927630 | MAP4 | LOC396097 |
| DMC | 2 | 759278 | -8 | downstream | 97:XM_046927637:dov | XM_046927637 | MAP4 | LOC396097 |
| DMC | 2 | 759278 | -8 | downstream | 97:XM_046927641:dov | XM_046927641 | MAP4 | LOC396097 |

| | | | | | | | | |
|---|---|---|---|---|---|---|---|---|
| DMC | 2 | 759278 | -8 | downstream | 97:XM_046927650:dov | XM_046927650 | MAP4 | LOC396097 |
| DMC | 2 | 759278 | -8 | downstream | 97:XM_046927654:dov | XM_046927654 | MAP4 | LOC396097 |
| DMC | 2 | 759278 | -8 | downstream | 97:XM_040696466:dov | XM_040696466 | MAP4 | LOC396097 |
| DMC | 2 | 759278 | -8 | downstream | 97:XM_040696467:dov | XM_040696467 | MAP4 | LOC396097 |
| DMC | 2 | 759278 | -8 | downstream | 97:XM_040696465:dov | XM_040696465 | MAP4 | LOC396097 |
| DMC | 2 | 5702062 | 8741 | other | NA | NA | NA | NA |
| DMC | 2 | 26053487 | -5404 | introns | ;83:XM_015281401:intr | XM_015281401 | PHF14 | LOC420583 |
| DMC | 2 | 26053487 | -5404 | introns | ;83:XM_015281402:intr | XM_015281402 | PHF14 | LOC420583 |
| DMC | 2 | 26053487 | -5404 | introns | ;83:NM_001012856:intı | NM_001012856 | PHF14 | LOC420583 |
| DMC | 2 | 42525595 | -119 | promoter | )96141:NM_205218:proı | NM_205218 | BFSP2 | LOC396141 |
| DMC | 2 | 42525595 | -119 | exons | )96141:NM_205218:exc | NM_205218 | BFSP2 | LOC396141 |
| DMC | 2 | 42525595 | -119 | exon1 | )96141:NM_205218:exc | NM_205218 | BFSP2 | LOC396141 |
| DMC | 2 | 42525595 | -119 | cds | )96141:NM_205218:exc | NM_205218 | BFSP2 | LOC396141 |
| DMC | 2 | 61654778 | -443 | introns | 20846:NR_131063:intr( | NR_131063 | RBM24 | LOC420846 |
| DMC | 2 | 61654778 | -443 | introns | )846:NM_001012863:in | NM_001012863 | RBM24 | LOC420846 |
| DMC | 2 | 61654778 | -443 | introns | )846:XM_015275877:in | XM_015275877 | RBM24 | LOC420846 |
| DMC | 2 | 61654778 | -443 | introns | 843:XR_005855807:intı | XR_005855807 | ATXN1 | LOC420843 |
| DMC | 2 | 61654778 | -443 | introns | 843:XR_005855809:intı | XR_005855809 | ATXN1 | LOC420843 |
| DMC | 2 | 61654778 | -443 | introns | 843:XR_005855813:intı | XR_005855813 | ATXN1 | LOC420843 |
| DMC | 2 | 61654778 | -443 | introns | 843:XR_005855821:intı | XR_005855821 | ATXN1 | LOC420843 |
| DMC | 2 | 61654778 | -443 | introns | 843:XR_005855822:intı | XR_005855822 | ATXN1 | LOC420843 |
| DMC | 2 | 61654778 | -443 | introns | 843:XR_005855824:intı | XR_005855824 | ATXN1 | LOC420843 |
| DMC | 2 | 61654778 | -443 | introns | 843:XR_006937206:intı | XR_006937206 | ATXN1 | LOC420843 |
| DMC | 2 | 61654778 | -443 | introns | 843:XR_006937207:intı | XR_006937207 | ATXN1 | LOC420843 |
| DMC | 2 | 61654833 | -498 | introns | 20846:NR_131063:intr( | NR_131063 | RBM24 | LOC420846 |
| DMC | 2 | 61654833 | -498 | introns | )846:NM_001012863:in | NM_001012863 | RBM24 | LOC420846 |
| DMC | 2 | 61654833 | -498 | introns | )846:XM_015275877:in | XM_015275877 | RBM24 | LOC420846 |
| DMC | 2 | 61654833 | -498 | introns | 843:XR_005855807:intı | XR_005855807 | ATXN1 | LOC420843 |
| DMC | 2 | 61654833 | -498 | introns | 843:XR_005855809:intı | XR_005855809 | ATXN1 | LOC420843 |
| DMC | 2 | 61654833 | -498 | introns | 843:XR_005855813:intı | XR_005855813 | ATXN1 | LOC420843 |
| DMC | 2 | 61654833 | -498 | introns | 843:XR_005855821:intı | XR_005855821 | ATXN1 | LOC420843 |
| DMC | 2 | 61654833 | -498 | introns | 843:XR_005855822:intı | XR_005855822 | ATXN1 | LOC420843 |
| DMC | 2 | 61654833 | -498 | introns | 843:XR_005855824:intı | XR_005855824 | ATXN1 | LOC420843 |
| DMC | 2 | 61654833 | -498 | introns | 843:XR_006937206:intı | XR_006937206 | ATXN1 | LOC420843 |
| DMC | 2 | 61654833 | -498 | introns | 843:XR_006937207:intı | XR_006937207 | ATXN1 | LOC420843 |
| DMC | 2 | 64404490 | 1374 | exons | 5920:XM_040675879:e: | XM_040675879 | RREB1 | LOC395920 |
| DMC | 2 | 64404490 | 1374 | exons | ;920:XM_046918752:ex | XM_046918752 | RREB1 | LOC395920 |
| DMC | 2 | 64404490 | 1374 | exons | )5920:NM_205049:exo | NM_205049 | RREB1 | LOC395920 |
| DMC | 2 | 64404490 | 1374 | exons | 920:XM_040675858:ex( | XM_040675858 | RREB1 | LOC395920 |
| DMC | 2 | 64404490 | 1374 | exons | ;920:XM_015275839:ex | XM_015275839 | RREB1 | LOC395920 |
| DMC | 2 | 64404490 | 1374 | exons | 920:XM_040675869:ex( | XM_040675869 | RREB1 | LOC395920 |
| DMC | 2 | 64404490 | 1374 | cds | 5920:XM_040675879:e: | XM_040675879 | RREB1 | LOC395920 |
| DMC | 2 | 64404490 | 1374 | cds | 5920:XM_046918752:e: | XM_046918752 | RREB1 | LOC395920 |
| DMC | 2 | 64404490 | 1374 | cds | )5920:NM_205049:exo | NM_205049 | RREB1 | LOC395920 |
| DMC | 2 | 64404490 | 1374 | cds | ;920:XM_040675858:ex | XM_040675858 | RREB1 | LOC395920 |
| DMC | 2 | 64404490 | 1374 | cds | ;920:XM_015275839:ex | XM_015275839 | RREB1 | LOC395920 |
| DMC | 2 | 64404490 | 1374 | cds | ;920:XM_040675869:ex | XM_040675869 | RREB1 | LOC395920 |
| DMC | 2 | 112581662 | 256 | promoter | .140:XM_046925616:pr | XM_046925616 | CHD7 | LOC421140 |
| DMC | 2 | 112581662 | 256 | promoter | 52732:XR_006938601:ı | XR_006938601 | LOC107052732 | LOC107052732 |
| DMC | 2 | 112581662 | 256 | promoter | 52732:XR_001465417:ı | XR_001465417 | LOC107052732 | LOC107052732 |
| DMC | 2 | 112581662 | 256 | introns | 140:XM_046925594:int | XM_046925594 | CHD7 | LOC421140 |
| DMC | 2 | 112581662 | 256 | introns | 140:XM_046925604:int | XM_046925604 | CHD7 | LOC421140 |
| DMC | 2 | 112581662 | 256 | introns | 140:XM_046925610:int | XM_046925610 | CHD7 | LOC421140 |
| DMC | 2 | 112581662 | 256 | introns | 140:NM_001077586:int | NM_001077586 | CHD7 | LOC421140 |
| DMC | 2 | 112581662 | 256 | introns | 140:XM_046925587:int | XM_046925587 | CHD7 | LOC421140 |
| DMC | 2 | 112581662 | 256 | introns | 140:XM_046925627:int | XM_046925627 | CHD7 | LOC421140 |
| DMC | 2 | 112581662 | 256 | introns | 140:XM_046925636:int | XM_046925636 | CHD7 | LOC421140 |
| DMC | 2 | 112581662 | 256 | introns | 140:XM_046925644:int | XM_046925644 | CHD7 | LOC421140 |
| DMC | 2 | 112581662 | 256 | intron1 | 140:XM_046925594:int | XM_046925594 | CHD7 | LOC421140 |
| DMC | 2 | 112581662 | 256 | intron1 | 140:XM_046925604:int | XM_046925604 | CHD7 | LOC421140 |
| DMC | 2 | 112581662 | 256 | intron1 | 140:XM_046925610:int | XM_046925610 | CHD7 | LOC421140 |
| DMC | 2 | 112581662 | 256 | intron1 | 140:NM_001077586:int | NM_001077586 | CHD7 | LOC421140 |
| DMC | 2 | 112581662 | 256 | intron1 | 140:XM_046925587:int | XM_046925587 | CHD7 | LOC421140 |
| DMC | 2 | 112581662 | 256 | intron1 | 140:XM_046925627:int | XM_046925627 | CHD7 | LOC421140 |
| DMC | 2 | 112581662 | 256 | intron1 | 140:XM_046925636:int | XM_046925636 | CHD7 | LOC421140 |
| DMC | 2 | 112581662 | 256 | intron1 | 140:XM_046925644:int | XM_046925644 | CHD7 | LOC421140 |
| DMC | 2 | 117258848 | 13248 | introns | .181:XM_025147998:in | XM_025147998 | KCNB2 | LOC431181 |

| | | | | | | | | |
|---|---|---|---|---|---|---|---|---|
| DMC | 2 | 117258848 | 13248 | introns | 181:XM_003640825:in | XM_003640825 | KCNB2 | LOC431181 |
| DMC | 2 | 117258848 | 13248 | introns | 181:XM_046935636:in | XM_046935636 | KCNB2 | LOC431181 |
| DMC | 2 | 117447407 | 231 | promoter | 0182:NM_001006345:pr | NM_001006345 | RPL7 | LOC420182 |
| DMC | 2 | 117447407 | 231 | promoter | 0182:XM_040677603:pr | XM_040677603 | RPL7 | LOC420182 |
| DMC | 2 | 117447407 | 231 | introns | 0183:NM_001199459:in | NM_001199459 | RDH10 | LOC420183 |
| DMC | 2 | 117447407 | 231 | introns | 0183:XM_015282780:in | XM_015282780 | RDH10 | LOC420183 |
| DMC | 2 | 117447407 | 231 | intron1 | 0183:NM_001199459:in | NM_001199459 | RDH10 | LOC420183 |
| DMC | 2 | 117447407 | 231 | intron1 | 0183:XM_015282780:in | XM_015282780 | RDH10 | LOC420183 |
| DMC | 2 | 117447407 | 231 | downstream | 3132:XR_005858031:do | XR_005858031 | LOC121113132 | LOC121113132 |
| DMC | 2 | 120841166 | -1047 | promoter | 0195:XM_040695800:pr | XM_040695800 | PAG1 | LOC420195 |
| DMC | 2 | 120841166 | -1047 | UTR5 | 95:XM_046937995:5'UT | XM_046937995 | PAG1 | LOC420195 |
| DMC | 2 | 120841166 | -1047 | exons | 0195:XM_046937995:e | XM_046937995 | PAG1 | LOC420195 |
| DMC | 2 | 120841166 | -1047 | introns | 0195:XM_046937996:in | XM_046937996 | PAG1 | LOC420195 |
| DMC | 2 | 120841166 | -1047 | introns | 0195:XM_015282858:in | XM_015282858 | PAG1 | LOC420195 |
| DMC | 2 | 120841166 | -1047 | introns | 0195:XM_025148017:in | XM_025148017 | PAG1 | LOC420195 |
| DMC | 2 | 121642303 | 1957 | introns | 013432:ENSGALT00010 | NSGALT0001003233 | NSGALG00010013 | NSGALG0001001343 |
| DMC | 3 | 4570946 | -213 | promoter | 1257:XR_006938646:pr | XR_006938646 | PTK7 | LOC421257 |
| DMC | 3 | 4570946 | -213 | promoter | 257:NM_001031035:pr | NM_001031035 | PTK7 | LOC421257 |
| DMC | 3 | 4570946 | -213 | promoter | 257:XM_015283375:pr | XM_015283375 | PTK7 | LOC421257 |
| DMC | 3 | 4570946 | -213 | promoter | 257:XM_046938693:pr | XM_046938693 | PTK7 | LOC421257 |
| DMC | 3 | 4570946 | -213 | promoter | 257:XM_046938694:pr | XM_046938694 | PTK7 | LOC421257 |
| DMC | 3 | 4570946 | -213 | promoter | 257:XM_046938695:pr | XM_046938695 | PTK7 | LOC421257 |
| DMC | 3 | 4570946 | -213 | exons | 052850:XM_015283418 | XM_015283418 | LOC107052850 | LOC107052850 |
| DMC | 3 | 4570946 | -213 | UTR3 | 850:XM_015283418:3' | XM_015283418 | LOC107052850 | LOC107052850 |
| DMC | 3 | 7887938 | 33 | introns | 32054:XR_006938684:i | XR_006938684 | LOC112532054 | LOC112532054 |
| DMC | 3 | 10145632 | -938 | exons | 421274:XM_419343:exo | XM_419343 | CEP68 | LOC421274 |
| DMC | 3 | 10145632 | -938 | exons | 1274:XM_015278090:e | XM_015278090 | CEP68 | LOC421274 |
| DMC | 3 | 10145632 | -938 | exons | 1274:XM_015278091:e | XM_015278091 | CEP68 | LOC421274 |
| DMC | 3 | 10145632 | -938 | exons | 1274:XM_015278092:e | XM_015278092 | CEP68 | LOC421274 |
| DMC | 3 | 10145632 | -938 | exons | 1274:XM_015278093:e | XM_015278093 | CEP68 | LOC421274 |
| DMC | 3 | 10145632 | -938 | exons | 1274:XM_040697716:e | XM_040697716 | CEP68 | LOC421274 |
| DMC | 3 | 10145632 | -938 | exons | 1274:XM_040697717:e | XM_040697717 | CEP68 | LOC421274 |
| DMC | 3 | 10145632 | -938 | cds | 421274:XM_419343:exo | XM_419343 | CEP68 | LOC421274 |
| DMC | 3 | 10145632 | -938 | cds | 1274:XM_015278090:e | XM_015278090 | CEP68 | LOC421274 |
| DMC | 3 | 10145632 | -938 | cds | 1274:XM_015278091:e | XM_015278091 | CEP68 | LOC421274 |
| DMC | 3 | 10145632 | -938 | cds | 1274:XM_015278092:e | XM_015278092 | CEP68 | LOC421274 |
| DMC | 3 | 10145632 | -938 | cds | 1274:XM_015278093:e | XM_015278093 | CEP68 | LOC421274 |
| DMC | 3 | 10145632 | -938 | cds | 1274:XM_040697716:e | XM_040697716 | CEP68 | LOC421274 |
| DMC | 3 | 10145632 | -938 | cds | 1274:XM_040697717:e | XM_040697717 | CEP68 | LOC421274 |
| DMC | 3 | 12455537 | -302 | promoter | 019102:ENSGALT00010 | NSGALT0001004622 | NSGALG00010019 | NSGALG0001001910 |
| DMC | 3 | 12455537 | -302 | introns | 52858:XR_005858755:i | XR_005858755 | LOC107052858 | LOC107052858 |
| DMC | 3 | 12455537 | -302 | introns | 52858:XR_005858754:i | XR_005858754 | LOC107052858 | LOC107052858 |
| DMC | 3 | 12455537 | -302 | introns | 6740:XM_015283525:in | XM_015283525 | ISM1 | LOC416740 |
| DMC | 3 | 12463247 | -957 | introns | 52858:XR_005858754:i | XR_005858754 | LOC107052858 | LOC107052858 |
| DMC | 3 | 12463247 | -957 | downstream | 0202:DAVISGALT00202 | AVISGALT00202020 | AVISGALG0000202 | AVISGALG00002020 |
| DMC | 3 | 12698597 | 134 | promoter | 017700:TAGAGALT0000 | GAGALT0000002634 | GAGALG000000017 | GAGALG0000000177 |
| DMC | 3 | 12698597 | 134 | exons | 017700:TAGAGALT0000 | GAGALT0000002634 | GAGALG000000017 | GAGALG0000000177 |
| DMC | 3 | 12698597 | 134 | exon1 | 017700:TAGAGALT0000 | GAGALT0000002634 | GAGALG000000017 | GAGALG0000000177 |
| DMC | 3 | 12698597 | 134 | introns | 538:XM_046939517:int | XM_046939517 | SPTLC3 | LOC768538 |
| DMC | 3 | 12698597 | 134 | introns | 538:XM_046939520:int | XM_046939520 | SPTLC3 | LOC768538 |
| DMC | 3 | 87305106 | 173 | promoter | 644:NM_001024584:pr | NM_001024584 | HCRTR2 | LOC428644 |
| DMC | 3 | 87305106 | 173 | exons | 8644:NM_001024584:e | NM_001024584 | HCRTR2 | LOC428644 |
| DMC | 3 | 87305106 | 173 | exon1 | 8644:NM_001024584:e | NM_001024584 | HCRTR2 | LOC428644 |
| DMC | 3 | 87305106 | 173 | cds | 8644:NM_001024584:e | NM_001024584 | HCRTR2 | LOC428644 |
| DMC | 3 | 94371830 | 529 | introns | 004062:ENSGALT00010 | NSGALT0001000942 | NSGALG00010040 | NSGALG0001000406 |
| DMC | 3 | 94371830 | 529 | downstream | 3891:ENSGALT0001000 | NSGALT0001000900 | NSGALG00010038 | NSGALG0001000389 |
| DMC | 3 | 107122051 | 3184 | other | NA | NA | NA | NA |
| DMC | 3 | 107128966 | 13 | promoter | 58393:XR_001466211:p | XR_001466211 | LOC100858393 | LOC100858393 |
| DMC | 3 | 107128966 | 13 | promoter | 58393:XR_001466212:p | XR_001466212 | LOC100858393 | LOC100858393 |
| DMC | 3 | 107128966 | 13 | promoter | 58393:XR_006938775:p | XR_006938775 | LOC100858393 | LOC100858393 |
| DMC | 3 | 107128966 | 13 | promoter | 58393:XR_006938776:p | XR_006938776 | LOC100858393 | LOC100858393 |
| DMC | 3 | 107128966 | 13 | promoter | 58393:XR_006938777:p | XR_006938777 | LOC100858393 | LOC100858393 |
| DMC | 3 | 107128966 | 13 | UTR5 | 92:NM_001397479:5'UT | NM_001397479 | GATA4 | LOC396392 |
| DMC | 3 | 107128966 | 13 | UTR5 | 92:XM_046938577:5'UT | XM_046938577 | GATA4 | LOC396392 |
| DMC | 3 | 107128966 | 13 | UTR5 | 92:XM_046938576:5'UT | XM_046938576 | GATA4 | LOC396392 |
| DMC | 3 | 107128966 | 13 | UTR5 | 92:XM_046938578:5'UT | XM_046938578 | GATA4 | LOC396392 |
| DMC | 3 | 107128966 | 13 | exons | 6392:NM_001397479:e | NM_001397479 | GATA4 | LOC396392 |

| | | | | | | | | |
|---|---|---|---|---|---|---|---|---|
| DMC | 3 | 107128966 | 13 | exons | 6392:XM_046938577:e | XM_046938577 | GATA4 | LOC396392 |
| DMC | 3 | 107128966 | 13 | exons | 6392:NM_001293106:e | NM_001293106 | GATA4 | LOC396392 |
| DMC | 3 | 107128966 | 13 | exons | 6392:NM_001397478:e | NM_001397478 | GATA4 | LOC396392 |
| DMC | 3 | 107128966 | 13 | exons | 6392:XM_046938576:e | XM_046938576 | GATA4 | LOC396392 |
| DMC | 3 | 107128966 | 13 | exons | 6392:XM_046938578:e | XM_046938578 | GATA4 | LOC396392 |
| DMC | 3 | 107128966 | 13 | exons | 358393:XR_006938774: | XR_006938774 | LOC100858393 | LOC100858393 |
| DMC | 3 | 107128966 | 13 | exon1 | 6392:NM_001293106:e | NM_001293106 | GATA4 | LOC396392 |
| DMC | 3 | 107128966 | 13 | exon1 | 6392:NM_001397478:e | NM_001397478 | GATA4 | LOC396392 |
| DMC | 3 | 107128966 | 13 | cds | 6392:NM_001293106:e | NM_001293106 | GATA4 | LOC396392 |
| DMC | 3 | 107128966 | 13 | cds | 6392:NM_001397478:e | NM_001397478 | GATA4 | LOC396392 |
| DMC | 3 | 107129478 | -499 | promoter | 58393:XR_001466211:p | XR_001466211 | LOC100858393 | LOC100858393 |
| DMC | 3 | 107129478 | -499 | promoter | 58393:XR_001466212:p | XR_001466212 | LOC100858393 | LOC100858393 |
| DMC | 3 | 107129478 | -499 | promoter | 58393:XR_006938775:p | XR_006938775 | LOC100858393 | LOC100858393 |
| DMC | 3 | 107129478 | -499 | promoter | 58393:XR_006938776:p | XR_006938776 | LOC100858393 | LOC100858393 |
| DMC | 3 | 107129478 | -499 | promoter | 58393:XR_006938777:p | XR_006938777 | LOC100858393 | LOC100858393 |
| DMC | 3 | 107129478 | -499 | exons | 358393:XR_006938774: | XR_006938774 | LOC100858393 | LOC100858393 |
| DMC | 3 | 107129478 | -499 | introns | 392:NM_001397479:in | NM_001397479 | GATA4 | LOC396392 |
| DMC | 3 | 107129478 | -499 | introns | 392:XM_046938577:in | XM_046938577 | GATA4 | LOC396392 |
| DMC | 3 | 107129478 | -499 | introns | 392:NM_001293106:in | NM_001293106 | GATA4 | LOC396392 |
| DMC | 3 | 107129478 | -499 | introns | 392:NM_001397478:in | NM_001397478 | GATA4 | LOC396392 |
| DMC | 3 | 107129478 | -499 | introns | 392:XM_046938576:in | XM_046938576 | GATA4 | LOC396392 |
| DMC | 3 | 107129478 | -499 | introns | 392:XM_046938578:in | XM_046938578 | GATA4 | LOC396392 |
| DMC | 3 | 107129478 | -499 | intron1 | 392:NM_001293106:in | NM_001293106 | GATA4 | LOC396392 |
| DMC | 3 | 107129478 | -499 | intron1 | 392:NM_001397478:in | NM_001397478 | GATA4 | LOC396392 |
| DMC | 3 | 107134489 | -612 | promoter | 58393:XR_006938774:p | XR_006938774 | LOC100858393 | LOC100858393 |
| DMC | 3 | 107134489 | -612 | introns | 392:NM_001397479:in | NM_001397479 | GATA4 | LOC396392 |
| DMC | 3 | 107134489 | -612 | introns | 392:XM_046938577:in | XM_046938577 | GATA4 | LOC396392 |
| DMC | 3 | 107134489 | -612 | introns | 392:NM_001293106:in | NM_001293106 | GATA4 | LOC396392 |
| DMC | 3 | 107134489 | -612 | introns | 392:NM_001397478:in | NM_001397478 | GATA4 | LOC396392 |
| DMC | 3 | 107134489 | -612 | introns | 392:XM_046938576:in | XM_046938576 | GATA4 | LOC396392 |
| DMC | 3 | 107134489 | -612 | introns | 392:XM_046938578:in | XM_046938578 | GATA4 | LOC396392 |
| DMC | 3 | 107134489 | -612 | intron1 | 392:NM_001293106:in | NM_001293106 | GATA4 | LOC396392 |
| DMC | 3 | 107134489 | -612 | intron1 | 392:NM_001397478:in | NM_001397478 | GATA4 | LOC396392 |
| DMC | 3 | 107285743 | -20 | promoter | 2040:XM_025149201:pr | XM_025149201 | TRAM2 | LOC422040 |
| DMC | 3 | 107285743 | -20 | promoter | 2040:XM_015285175:pr | XM_015285175 | TRAM2 | LOC422040 |
| DMC | 3 | 107285743 | -20 | exons | 2039:XM_004935904:e | XM_004935904 | XKR5 | LOC422039 |
| DMC | 3 | 107285743 | -20 | UTR3 | 39:XM_004935904:3'UT | XM_004935904 | XKR5 | LOC422039 |
| DMC | 3 | 107285743 | -20 | downstream | 39:XM_003641074:dov | XM_003641074 | XKR5 | LOC422039 |
| DMC | 3 | 107285743 | -20 | downstream | 039:XR_001466208:dow | XR_001466208 | XKR5 | LOC422039 |
| DMC | 4 | 522082 | -1437 | introns | .3224:XM_040699969:i | XM_040699969 | STARD8 | LOC121113224 |
| DMC | 4 | 522082 | -1437 | introns | .3224:XM_046940744:i | XM_046940744 | STARD8 | LOC121113224 |
| DMC | 4 | 522082 | -1437 | introns | .3224:XM_040699968:i | XM_040699968 | STARD8 | LOC121113224 |
| DMC | 4 | 522082 | -1437 | intron1 | .3224:XM_046940744:i | XM_046940744 | STARD8 | LOC121113224 |
| DMC | 4 | 522082 | -1437 | intron1 | .3224:XM_040699968:i | XM_040699968 | STARD8 | LOC121113224 |
| DMC | 4 | 747663 | -381 | introns | 009658:NONGGAT01516 | NONGGAT015169 | NONGGAG009658 | NONGGAG009658 |
| DMC | 4 | 976557 | 5913 | other | NA | NA | NA | NA |
| DMC | 4 | 995188 | -890 | exons | 025192:DAVISGALT002 | DAVISGALT00251920 | AVISGALG0000251 | AVISGALG00002519 |
| DMC | 4 | 995188 | -890 | exon1 | 025192:DAVISGALT002 | DAVISGALT00251920 | AVISGALG0000251 | AVISGALG00002519 |
| DMC | 4 | 995188 | -890 | introns | 5896:XM_015278194:in | XM_015278194 | EFNB1 | LOC395896 |
| DMC | 4 | 995188 | -890 | introns | 5896:XM_015278195:in | XM_015278195 | EFNB1 | LOC395896 |
| DMC | 4 | 995188 | -890 | introns | 95896:NM_205035:intr | NM_205035 | EFNB1 | LOC395896 |
| DMC | 4 | 1008340 | -317 | exons | 011640:TAGAGALT0000 | GAGALT0000003140 | GAGALG000000011 | GAGALG0000000116 |
| DMC | 4 | 1008340 | -317 | introns | 95896:NM_205035:intr | NM_205035 | EFNB1 | LOC395896 |
| DMC | 4 | 1008340 | -317 | intron1 | 95896:NM_205035:intr | NM_205035 | EFNB1 | LOC395896 |
| DMC | 4 | 1040863 | -6401 | other | NA | NA | NA | NA |
| DMC | 4 | 1048857 | 7750 | other | NA | NA | NA | NA |
| DMC | 4 | 1076947 | 58 | exons | 9051:XM_015278333:e | XM_015278333 | FAM155B | LOC769051 |
| DMC | 4 | 1076947 | 58 | cds | 9051:XM_015278333:e | XM_015278333 | FAM155B | LOC769051 |
| DMC | 4 | 1106637 | -258 | exons | 019382:ENSGALT00010 | NSGALT0001004685 | NSGALG000100193 | NSGALG0001001938 |
| DMC | 4 | 1106637 | -258 | introns | 9069:XM_015278334:in | XM_015278334 | EDA | LOC769069 |
| DMC | 4 | 1106637 | -258 | introns | 9069:XM_015278335:in | XM_015278335 | EDA | LOC769069 |
| DMC | 4 | 1106637 | -258 | introns | 9069:XM_015278336:in | XM_015278336 | EDA | LOC769069 |
| DMC | 4 | 1106637 | -258 | introns | 019609:ENSGALT00010 | NSGALT0001004730 | NSGALG000100196 | NSGALG0001001960 |
| DMC | 4 | 1106637 | -258 | introns | 019380:ENSGALT00010 | NSGALT0001004685 | NSGALG000100193 | NSGALG0001001938 |
| DMC | 4 | 1106637 | -258 | intron1 | 9069:XM_015278334:in | XM_015278334 | EDA | LOC769069 |
| DMC | 4 | 1106637 | -258 | intron1 | 9069:XM_015278335:in | XM_015278335 | EDA | LOC769069 |
| DMC | 4 | 1106637 | -258 | intron1 | 9069:XM_015278336:in | XM_015278336 | EDA | LOC769069 |

| | | | | | | | | |
|---|---|---|---|---|---|---|---|---|
| DMC | 4 | 1106637 | -258 | intron1 | 19609:ENSGALT00010 | NSGALT0001004730 | NSGALG000100196 | NSGALG0001001960 |
| DMC | 4 | 1106637 | -258 | intron1 | 19380:ENSGALT00010 | NSGALT0001004685 | NSGALG000100193 | NSGALG0001001938 |
| DMC | 4 | 1117489 | -5548 | introns | 069:XM_015278334:in | XM_015278334 | EDA | LOC769069 |
| DMC | 4 | 1117489 | -5548 | introns | 069:XM_015278335:in | XM_015278335 | EDA | LOC769069 |
| DMC | 4 | 1117489 | -5548 | introns | 069:XM_015278336:in | XM_015278336 | EDA | LOC769069 |
| DMC | 4 | 1117489 | -5548 | intron1 | 069:XM_015278334:in | XM_015278334 | EDA | LOC769069 |
| DMC | 4 | 1117489 | -5548 | intron1 | 069:XM_015278335:in | XM_015278335 | EDA | LOC769069 |
| DMC | 4 | 1117489 | -5548 | intron1 | 069:XM_015278336:in | XM_015278336 | EDA | LOC769069 |
| DMC | 4 | 1138677 | 399 | introns | 069:XM_015278334:in | XM_015278334 | EDA | LOC769069 |
| DMC | 4 | 1138677 | 399 | introns | 069:XM_015278335:in | XM_015278335 | EDA | LOC769069 |
| DMC | 4 | 1138677 | 399 | introns | 069:XM_015278336:in | XM_015278336 | EDA | LOC769069 |
| DMC | 4 | 1144432 | 700 | introns | 069:XM_015278334:in | XM_015278334 | EDA | LOC769069 |
| DMC | 4 | 1144432 | 700 | introns | 069:XM_015278335:in | XM_015278335 | EDA | LOC769069 |
| DMC | 4 | 1144432 | 700 | introns | 069:XM_015278336:in | XM_015278336 | EDA | LOC769069 |
| DMC | 4 | 1175536 | -60 | promoter | 693:NM_001398400:pr | NM_001398400 | AWAT1 | LOC428693 |
| DMC | 4 | 1175536 | -60 | promoter | 693:XM_046940301:pr | XM_046940301 | AWAT1 | LOC428693 |
| DMC | 4 | 1175536 | -60 | promoter | 693:NM_001318429:pr | NM_001318429 | AWAT1 | LOC428693 |
| DMC | 4 | 1175536 | -60 | introns | 693:XM_040699093:in | XM_040699093 | AWAT1 | LOC428693 |
| DMC | 4 | 1175536 | -60 | introns | 693:XM_046940300:in | XM_046940300 | AWAT1 | LOC428693 |
| DMC | 4 | 1175536 | -60 | introns | 693:NM_001398400:in | NM_001398400 | AWAT1 | LOC428693 |
| DMC | 4 | 1175536 | -60 | introns | 693:XM_046940301:in | XM_046940301 | AWAT1 | LOC428693 |
| DMC | 4 | 1175536 | -60 | introns | 693:NM_001318429:in | NM_001318429 | AWAT1 | LOC428693 |
| DMC | 4 | 1175536 | -60 | intron1 | 693:NM_001398400:in | NM_001398400 | AWAT1 | LOC428693 |
| DMC | 4 | 1175536 | -60 | intron1 | 693:XM_046940301:in | XM_046940301 | AWAT1 | LOC428693 |
| DMC | 4 | 1175536 | -60 | intron1 | 693:NM_001318429:in | NM_001318429 | AWAT1 | LOC428693 |
| DMC | 4 | 1219130 | -38 | promoter | 22154:XM_420155:pro | XM_420155 | LOC422154 | LOC422154 |
| DMC | 4 | 1219130 | -38 | promoter | 154:XM_040699978:pr | XM_040699978 | LOC422154 | LOC422154 |
| DMC | 4 | 1219130 | -38 | exons | 22154:XM_420155:exo | XM_420155 | LOC422154 | LOC422154 |
| DMC | 4 | 1219130 | -38 | exon1 | 22154:XM_420155:exo | XM_420155 | LOC422154 | LOC422154 |
| DMC | 4 | 1219130 | -38 | introns | 154:XM_040699977:in | XM_040699977 | LOC422154 | LOC422154 |
| DMC | 4 | 1219130 | -38 | intron1 | 154:XM_040699977:in | XM_040699977 | LOC422154 | LOC422154 |
| DMC | 4 | 1219130 | -38 | cds | 22154:XM_420155:exo | XM_420155 | LOC422154 | LOC422154 |
| DMC | 4 | 1219130 | -38 | downstream | 1235:XM_015278352:d | XM_015278352 | RAB41 | LOC101751235 |
| DMC | 4 | 1219130 | -38 | downstream | 1235:XM_015278354:d | XM_015278354 | RAB41 | LOC101751235 |
| DMC | 4 | 1219130 | -38 | downstream | 1235:XM_040699980:d | XM_040699980 | RAB41 | LOC101751235 |
| DMC | 4 | 1219130 | -38 | downstream | 1235:XM_040699981:d | XM_040699981 | RAB41 | LOC101751235 |
| DMC | 4 | 1219130 | -38 | downstream | 1235:XM_040699982:d | XM_040699982 | RAB41 | LOC101751235 |
| DMC | 4 | 1219130 | -38 | downstream | 1235:XM_040699983:d | XM_040699983 | RAB41 | LOC101751235 |
| DMC | 4 | 1219130 | -38 | downstream | 1235:XM_040699984:d | XM_040699984 | RAB41 | LOC101751235 |
| DMC | 4 | 1219130 | -38 | downstream | 1235:XM_040699985:d | XM_040699985 | RAB41 | LOC101751235 |
| DMC | 4 | 1219130 | -38 | downstream | 1235:XM_040699986:d | XM_040699986 | RAB41 | LOC101751235 |
| DMC | 4 | 1219130 | -38 | downstream | 1235:XR_005859738:do | XR_005859738 | RAB41 | LOC101751235 |
| DMC | 4 | 1219130 | -38 | downstream | 1235:XM_046940945:d | XM_046940945 | RAB41 | LOC101751235 |
| DMC | 4 | 1219130 | -38 | downstream | 1235:XM_046940946:d | XM_046940946 | RAB41 | LOC101751235 |
| DMC | 4 | 1604540 | 1687 | downstream | 36:XM_004940575:dov | XM_004940575 | OCRL | LOC422136 |
| DMC | 4 | 1604540 | 1687 | downstream | 36:XM_004940576:dov | XM_004940576 | OCRL | LOC422136 |
| DMC | 4 | 1604540 | 1687 | downstream | 36:XM_004940577:dov | XM_004940577 | OCRL | LOC422136 |
| DMC | 4 | 1604540 | 1687 | downstream | 36:XM_040699990:dov | XM_040699990 | OCRL | LOC422136 |
| DMC | 4 | 1604540 | 1687 | downstream | 2136:XM_420138:down | XM_420138 | OCRL | LOC422136 |
| DMC | 4 | 1787553 | 5 | introns | 179:XM_015278377:in | XM_015278377 | EVA1CL | LOC422179 |
| DMC | 4 | 1787553 | 5 | introns | 179:XM_015278378:in | XM_015278378 | EVA1CL | LOC422179 |
| DMC | 4 | 1787553 | 5 | introns | 179:XM_015278379:in | XM_015278379 | EVA1CL | LOC422179 |
| DMC | 4 | 1787553 | 5 | introns | 179:XM_025150328:in | XM_025150328 | EVA1CL | LOC422179 |
| DMC | 4 | 1787553 | 5 | introns | 179:XM_025150329:in | XM_025150329 | EVA1CL | LOC422179 |
| DMC | 4 | 1787553 | 5 | introns | 179:XM_040699996:in | XM_040699996 | EVA1CL | LOC422179 |
| DMC | 4 | 1787553 | 5 | introns | 2179:XR_003075082:int | XR_003075082 | EVA1CL | LOC422179 |
| DMC | 4 | 1787553 | 5 | introns | 179:XM_025150330:in | XM_025150330 | EVA1CL | LOC422179 |
| DMC | 4 | 1787553 | 5 | introns | 179:XM_025150331:in | XM_025150331 | EVA1CL | LOC422179 |
| DMC | 4 | 1787553 | 5 | introns | 179:XM_040699997:in | XM_040699997 | EVA1CL | LOC422179 |
| DMC | 4 | 1787553 | 5 | introns | 179:XM_040699998:in | XM_040699998 | EVA1CL | LOC422179 |
| DMC | 4 | 1787553 | 5 | introns | 179:XM_046940953:in | XM_046940953 | EVA1CL | LOC422179 |
| DMC | 4 | 1787553 | 5 | introns | 22179:XM_420175:intr | XM_420175 | EVA1CL | LOC422179 |
| DMC | 4 | 1787553 | 5 | introns | 2179:XR_006939241:int | XR_006939241 | EVA1CL | LOC422179 |
| DMC | 4 | 1787553 | 5 | intron1 | 179:XM_015278377:in | XM_015278377 | EVA1CL | LOC422179 |
| DMC | 4 | 1787553 | 5 | intron1 | 179:XM_015278378:in | XM_015278378 | EVA1CL | LOC422179 |
| DMC | 4 | 1787553 | 5 | intron1 | 179:XM_015278379:in | XM_015278379 | EVA1CL | LOC422179 |
| DMC | 4 | 1787553 | 5 | intron1 | 179:XM_025150328:in | XM_025150328 | EVA1CL | LOC422179 |

| | | | | | | | | |
|---|---|---|---|---|---|---|---|---|
| DMC | 4 | 1787553 | 5 | intron1 | !179:XM_025150329:in | XM_025150329 | EVA1CL | LOC422179 |
| DMC | 4 | 1787553 | 5 | intron1 | !179:XM_040699996:in | XM_040699996 | EVA1CL | LOC422179 |
| DMC | 4 | 1787553 | 5 | intron1 | !179:XR_003075082:int | XR_003075082 | EVA1CL | LOC422179 |
| DMC | 4 | 1787553 | 5 | intron1 | !179:XM_040699998:in | XM_040699998 | EVA1CL | LOC422179 |
| DMC | 4 | 1787620 | -62 | promoter | 96167:NM_205239:pro | NM_205239 | CCNB3 | LOC396167 |
| DMC | 4 | 1787620 | -62 | promoter | 5167:XM_015278200:pr | XM_015278200 | CCNB3 | LOC396167 |
| DMC | 4 | 1787620 | -62 | promoter | 5167:XM_015278199:pr | XM_015278199 | CCNB3 | LOC396167 |
| DMC | 4 | 1787620 | -62 | promoter | 5167:XM_040698748:pr | XM_040698748 | CCNB3 | LOC396167 |
| DMC | 4 | 1787620 | -62 | exons | 2179:XM_015278377:e | XM_015278377 | EVA1CL | LOC422179 |
| DMC | 4 | 1787620 | -62 | exons | 2179:XM_015278378:e | XM_015278378 | EVA1CL | LOC422179 |
| DMC | 4 | 1787620 | -62 | exons | 2179:XM_015278379:e | XM_015278379 | EVA1CL | LOC422179 |
| DMC | 4 | 1787620 | -62 | exons | 2179:XM_025150328:e | XM_025150328 | EVA1CL | LOC422179 |
| DMC | 4 | 1787620 | -62 | exons | 2179:XM_025150329:e | XM_025150329 | EVA1CL | LOC422179 |
| DMC | 4 | 1787620 | -62 | exons | 2179:XM_040699996:e | XM_040699996 | EVA1CL | LOC422179 |
| DMC | 4 | 1787620 | -62 | exons | 2179:XR_003075082:e | XR_003075082 | EVA1CL | LOC422179 |
| DMC | 4 | 1787620 | -62 | exons | 2179:XM_025150330:e | XM_025150330 | EVA1CL | LOC422179 |
| DMC | 4 | 1787620 | -62 | exons | 2179:XM_025150331:e | XM_025150331 | EVA1CL | LOC422179 |
| DMC | 4 | 1787620 | -62 | exons | 2179:XM_040699997:e | XM_040699997 | EVA1CL | LOC422179 |
| DMC | 4 | 1787620 | -62 | exons | 2179:XM_046940953:e | XM_046940953 | EVA1CL | LOC422179 |
| DMC | 4 | 1787620 | -62 | exons | 422179:XM_420175:exo | XM_420175 | EVA1CL | LOC422179 |
| DMC | 4 | 1787620 | -62 | exons | 2179:XR_006939241:e | XR_006939241 | EVA1CL | LOC422179 |
| DMC | 4 | 1787620 | -62 | introns | !179:XM_040699998:in | XM_040699998 | EVA1CL | LOC422179 |
| DMC | 4 | 1787620 | -62 | intron1 | !179:XM_040699998:in | XM_040699998 | EVA1CL | LOC422179 |
| DMC | 4 | 1787620 | -62 | cds | 2179:XM_015278377:e | XM_015278377 | EVA1CL | LOC422179 |
| DMC | 4 | 1787620 | -62 | cds | 2179:XM_015278378:e | XM_015278378 | EVA1CL | LOC422179 |
| DMC | 4 | 1787620 | -62 | cds | 2179:XM_015278379:e | XM_015278379 | EVA1CL | LOC422179 |
| DMC | 4 | 1787620 | -62 | cds | 2179:XM_025150328:e | XM_025150328 | EVA1CL | LOC422179 |
| DMC | 4 | 1787620 | -62 | cds | 2179:XM_025150329:e | XM_025150329 | EVA1CL | LOC422179 |
| DMC | 4 | 1787620 | -62 | cds | 2179:XM_040699996:e | XM_040699996 | EVA1CL | LOC422179 |
| DMC | 4 | 1787620 | -62 | cds | 2179:XM_025150330:e | XM_025150330 | EVA1CL | LOC422179 |
| DMC | 4 | 1787620 | -62 | cds | 2179:XM_025150331:e | XM_025150331 | EVA1CL | LOC422179 |
| DMC | 4 | 1787620 | -62 | cds | 2179:XM_040699997:e | XM_040699997 | EVA1CL | LOC422179 |
| DMC | 4 | 1787620 | -62 | cds | 2179:XM_046940953:e | XM_046940953 | EVA1CL | LOC422179 |
| DMC | 4 | 1787620 | -62 | cds | 422179:XM_420175:exo | XM_420175 | EVA1CL | LOC422179 |
| DMC | 4 | 1795869 | 77 | exons | 129:XM_040700001:ex | XM_040700001 | DGKK | LOC422129 |
| DMC | 4 | 1795869 | 77 | exons | 129:XM_015278364:ex | XM_015278364 | DGKK | LOC422129 |
| DMC | 4 | 1795869 | 77 | exons | 129:XM_015278365:ex | XM_015278365 | DGKK | LOC422129 |
| DMC | 4 | 1795869 | 77 | exons | 129:XM_040700000:ex | XM_040700000 | DGKK | LOC422129 |
| DMC | 4 | 1795869 | 77 | cds | 129:XM_040700001:ex | XM_040700001 | DGKK | LOC422129 |
| DMC | 4 | 1795869 | 77 | cds | 129:XM_015278364:ex | XM_015278364 | DGKK | LOC422129 |
| DMC | 4 | 1795869 | 77 | cds | 129:XM_015278365:ex | XM_015278365 | DGKK | LOC422129 |
| DMC | 4 | 1795869 | 77 | cds | 129:XM_040700000:ex | XM_040700000 | DGKK | LOC422129 |
| DMC | 4 | 1802727 | -64 | introns | 129:XM_040700001:int | XM_040700001 | DGKK | LOC422129 |
| DMC | 4 | 1802727 | -64 | introns | 129:XM_015278364:int | XM_015278364 | DGKK | LOC422129 |
| DMC | 4 | 1802727 | -64 | introns | 129:XM_015278365:int | XM_015278365 | DGKK | LOC422129 |
| DMC | 4 | 1802727 | -64 | introns | 129:XM_040700000:int | XM_040700000 | DGKK | LOC422129 |
| DMC | 4 | 1815781 | -483 | promoter | 3697:NM_001006589:pr | NM_001006589 | BMP15 | LOC428697 |
| DMC | 4 | 1815781 | -483 | promoter | 7072:XM_015278366:pr | XM_015278366 | SHROOM4 | LOC777072 |
| DMC | 4 | 1815781 | -483 | promoter | 7072:XM_015278367:pr | XM_015278367 | SHROOM4 | LOC777072 |
| DMC | 4 | 1818418 | 187 | promoter | !181:XM_040700008:pr | XM_040700008 | PHKA1 | LOC422181 |
| DMC | 4 | 1818418 | 187 | promoter | !181:XM_040700009:pr | XM_040700009 | PHKA1 | LOC422181 |
| DMC | 4 | 1818418 | 187 | promoter | !181:XM_004940598:pr | XM_004940598 | PHKA1 | LOC422181 |
| DMC | 4 | 1818418 | 187 | promoter | !181:XM_004940599:pr | XM_004940599 | PHKA1 | LOC422181 |
| DMC | 4 | 1818418 | 187 | promoter | !181:XM_004940600:pr | XM_004940600 | PHKA1 | LOC422181 |
| DMC | 4 | 1818418 | 187 | exons | 3697:NM_001006589:e | NM_001006589 | BMP15 | LOC428697 |
| DMC | 4 | 1818418 | 187 | cds | 3697:NM_001006589:e | NM_001006589 | BMP15 | LOC428697 |
| DMC | 4 | 2008788 | -50 | promoter | 068379:TAGAGALT0000 | GAGALT00000031577 | GAGALG000000068 | GAGALG0000000683 |
| DMC | 4 | 2008788 | -50 | exons | 9785:XM_001233084:e | XM_001233084 | AIPL1 | LOC769785 |
| DMC | 4 | 2008788 | -50 | exons | 9785:XM_040700022:e | XM_040700022 | AIPL1 | LOC769785 |
| DMC | 4 | 2008788 | -50 | cds | 9785:XM_001233084:e | XM_001233084 | AIPL1 | LOC769785 |
| DMC | 4 | 2008788 | -50 | cds | 9785:XM_040700022:e | XM_040700022 | AIPL1 | LOC769785 |
| DMC | 4 | 2017161 | -147 | introns | .93:XM_015278394:intr | XM_015278394 | DRP2 | LOC422193 |
| DMC | 4 | 2017161 | -147 | introns | .93:XM_015278392:intr | XM_015278392 | DRP2 | LOC422193 |
| DMC | 4 | 2017161 | -147 | introns | .93:XM_015278393:intr | XM_015278393 | DRP2 | LOC422193 |
| DMC | 4 | 2017161 | -147 | introns | .93:XM_025150290:intr | XM_025150290 | DRP2 | LOC422193 |
| DMC | 4 | 2017161 | -147 | introns | .93:XM_040700012:intr | XM_040700012 | DRP2 | LOC422193 |
| DMC | 4 | 2017161 | -147 | introns | .93:XM_040700020:intr | XM_040700020 | DRP2 | LOC422193 |

| | | | | | | | | |
|---|---|---|---|---|---|---|---|---|
| DMC | 4 | 2017161 | -147 | downstream | 2017:INRAGALT000000 | IRAGALT0000002522 | RAGALG000000120 | IRAGALG0000001201 |
| DMC | 4 | 2020846 | 4 | exons | 2193:XM_015278394:ex | XM_015278394 | DRP2 | LOC422193 |
| DMC | 4 | 2020846 | 4 | exons | 2193:XM_015278392:ex | XM_015278392 | DRP2 | LOC422193 |
| DMC | 4 | 2020846 | 4 | exons | 2193:XM_015278393:ex | XM_015278393 | DRP2 | LOC422193 |
| DMC | 4 | 2020846 | 4 | exons | 2193:XM_025150290:ex | XM_025150290 | DRP2 | LOC422193 |
| DMC | 4 | 2020846 | 4 | exons | 2193:XM_040700012:ex | XM_040700012 | DRP2 | LOC422193 |
| DMC | 4 | 2020846 | 4 | exons | 2193:XM_040700020:ex | XM_040700020 | DRP2 | LOC422193 |
| DMC | 4 | 2020846 | 4 | cds | 2193:XM_015278394:ex | XM_015278394 | DRP2 | LOC422193 |
| DMC | 4 | 2020846 | 4 | cds | 2193:XM_015278392:ex | XM_015278392 | DRP2 | LOC422193 |
| DMC | 4 | 2020846 | 4 | cds | 2193:XM_015278393:ex | XM_015278393 | DRP2 | LOC422193 |
| DMC | 4 | 2020846 | 4 | cds | 2193:XM_025150290:ex | XM_025150290 | DRP2 | LOC422193 |
| DMC | 4 | 2020846 | 4 | cds | 2193:XM_040700012:ex | XM_040700012 | DRP2 | LOC422193 |
| DMC | 4 | 2020846 | 4 | cds | 2193:XM_040700020:ex | XM_040700020 | DRP2 | LOC422193 |
| DMC | 4 | 2025252 | -413 | promoter | 2193:XM_015278394:pr | XM_015278394 | DRP2 | LOC422193 |
| DMC | 4 | 2025252 | -413 | introns | 193:XM_015278392:int | XM_015278392 | DRP2 | LOC422193 |
| DMC | 4 | 2025252 | -413 | introns | 193:XM_015278393:int | XM_015278393 | DRP2 | LOC422193 |
| DMC | 4 | 2025252 | -413 | introns | 193:XM_025150290:int | XM_025150290 | DRP2 | LOC422193 |
| DMC | 4 | 2025252 | -413 | introns | 193:XM_040700012:int | XM_040700012 | DRP2 | LOC422193 |
| DMC | 4 | 2025252 | -413 | introns | 193:XM_040700020:int | XM_040700020 | DRP2 | LOC422193 |
| DMC | 4 | 2025252 | -413 | intron1 | 193:XM_015278392:int | XM_015278392 | DRP2 | LOC422193 |
| DMC | 4 | 2025252 | -413 | intron1 | 193:XM_015278393:int | XM_015278393 | DRP2 | LOC422193 |
| DMC | 4 | 2025252 | -413 | intron1 | 193:XM_025150290:int | XM_025150290 | DRP2 | LOC422193 |
| DMC | 4 | 2025252 | -413 | intron1 | 193:XM_040700012:int | XM_040700012 | DRP2 | LOC422193 |
| DMC | 4 | 2025252 | -413 | intron1 | 193:XM_040700020:int | XM_040700020 | DRP2 | LOC422193 |
| DMC | 4 | 2435902 | -37 | introns | 706:XM_040700055:int | XM_040700055 | DLG3 | LOC428706 |
| DMC | 4 | 2435902 | -37 | introns | 706:XM_015278421:int | XM_015278421 | DLG3 | LOC428706 |
| DMC | 4 | 2435902 | -37 | introns | 706:XM_015278422:int | XM_015278422 | DLG3 | LOC428706 |
| DMC | 4 | 2435902 | -37 | introns | 706:XM_040700056:int | XM_040700056 | DLG3 | LOC428706 |
| DMC | 4 | 2435902 | -37 | introns | 706:XM_046940963:int | XM_046940963 | DLG3 | LOC428706 |
| DMC | 4 | 2435902 | -37 | introns | 706:XM_046940962:int | XM_046940962 | DLG3 | LOC428706 |
| DMC | 4 | 2435902 | -37 | introns | '06:XM_046940961:intr | XM_046940961 | DLG3 | LOC428706 |
| DMC | 4 | 2435902 | -37 | introns | '06:XM_015278412:intr | XM_015278412 | DLG3 | LOC428706 |
| DMC | 4 | 2435902 | -37 | introns | '06:XM_040700053:intr | XM_040700053 | DLG3 | LOC428706 |
| DMC | 4 | 2435902 | -37 | introns | '06:XM_046940960:intr | XM_046940960 | DLG3 | LOC428706 |
| DMC | 4 | 2435902 | -37 | introns | 8706:XM_426264:intro | XM_426264 | DLG3 | LOC428706 |
| DMC | 4 | 2449662 | -320 | promoter | 8706:XM_015278421:pr | XM_015278421 | DLG3 | LOC428706 |
| DMC | 4 | 2449662 | -320 | promoter | 8706:XM_015278422:pr | XM_015278422 | DLG3 | LOC428706 |
| DMC | 4 | 2449662 | -320 | promoter | 8706:XM_040700056:pr | XM_040700056 | DLG3 | LOC428706 |
| DMC | 4 | 2449662 | -320 | promoter | 8706:XM_046940963:pr | XM_046940963 | DLG3 | LOC428706 |
| DMC | 4 | 2449662 | -320 | promoter | 8706:XM_046940962:pr | XM_046940962 | DLG3 | LOC428706 |
| DMC | 4 | 2449662 | -320 | introns | 706:XM_046940962:int | XM_046940962 | DLG3 | LOC428706 |
| DMC | 4 | 2449662 | -320 | introns | 706:XM_046940961:int | XM_046940961 | DLG3 | LOC428706 |
| DMC | 4 | 2449662 | -320 | introns | '06:XM_015278412:intr | XM_015278412 | DLG3 | LOC428706 |
| DMC | 4 | 2449662 | -320 | introns | '06:XM_040700053:intr | XM_040700053 | DLG3 | LOC428706 |
| DMC | 4 | 2449662 | -320 | introns | '06:XM_046940960:intr | XM_046940960 | DLG3 | LOC428706 |
| DMC | 4 | 2449662 | -320 | introns | 8706:XM_426264:intro | XM_426264 | DLG3 | LOC428706 |
| DMC | 4 | 2449662 | -320 | intron1 | 706:XM_046940962:int | XM_046940962 | DLG3 | LOC428706 |
| DMC | 4 | 2858109 | -202 | exons | 110553:XR_005859750: | XR_005859750 | LOC121110553 | LOC121110553 |
| DMC | 4 | 2858109 | -202 | introns | 10554:XR_005859751:i | XR_005859751 | LOC121110554 | LOC121110554 |
| DMC | 4 | 2858109 | -202 | intron1 | 10554:XR_005859751:i | XR_005859751 | LOC121110554 | LOC121110554 |
| DMC | 4 | 2858139 | -232 | exons | 110553:XR_005859750: | XR_005859750 | LOC121110553 | LOC121110553 |
| DMC | 4 | 2858139 | -232 | introns | 10554:XR_005859751:i | XR_005859751 | LOC121110554 | LOC121110554 |
| DMC | 4 | 2858139 | -232 | intron1 | 10554:XR_005859751:i | XR_005859751 | LOC121110554 | LOC121110554 |
| DMC | 4 | 2858165 | -258 | exons | 110553:XR_005859750: | XR_005859750 | LOC121110553 | LOC121110553 |
| DMC | 4 | 2858165 | -258 | introns | 10554:XR_005859751:i | XR_005859751 | LOC121110554 | LOC121110554 |
| DMC | 4 | 2858165 | -258 | intron1 | 10554:XR_005859751:i | XR_005859751 | LOC121110554 | LOC121110554 |
| DMC | 4 | 2863959 | -100 | promoter | 012033:INRAGALT00000 | IRAGALT0000002525 | RAGALG000000120 | IRAGALG0000001203 |
| DMC | 4 | 2863959 | -100 | exons | 2221:XM_015278433:e | XM_015278433 | FYTTD1L | LOC422221 |
| DMC | 4 | 2863959 | -100 | exons | 422221:XM_420211:exo | XM_420211 | FYTTD1L | LOC422221 |
| DMC | 4 | 2863959 | -100 | introns | 10554:XR_005859751:i | XR_005859751 | LOC121110554 | LOC121110554 |
| DMC | 4 | 2863959 | -100 | intron1 | 10554:XR_005859751:i | XR_005859751 | LOC121110554 | LOC121110554 |
| DMC | 4 | 2863959 | -100 | cds | 2221:XM_015278433:e | XM_015278433 | FYTTD1L | LOC422221 |
| DMC | 4 | 2863959 | -100 | cds | 422221:XM_420211:exo | XM_420211 | FYTTD1L | LOC422221 |
| DMC | 4 | 28935161 | -1131 | exons | 009577:ENSGALT00010 | NSGALT0001002282 | NSGALG000100095 | NSGALG0001000957 |
| DMC | 4 | 28935161 | -1131 | introns | 009577:ENSGALT00010 | NSGALT0001002281 | NSGALG000100095 | NSGALG0001000957 |
| DMC | 4 | 28935161 | -1131 | introns | 009577:ENSGALT00010 | NSGALT0001002282 | NSGALG000100095 | NSGALG0001000957 |
| DMC | 4 | 28935161 | -1131 | intron1 | 009577:ENSGALT00010 | NSGALT0001002281 | NSGALG000100095 | NSGALG0001000957 |

| | | | | | | | | |
|---|---|---|---|---|---|---|---|---|
| DMC | 4 | 28935161 | -1131 | intron1 | 09577:ENSGALT00010( | NSGALT0001002282 | NSGALG000100095 | NSGALG0001000957 |
| DMC | 4 | 36617413 | 597 | promoter | 50213:XM_004941130: | XM_004941130 | ATOH1 | LOC101750213 |
| DMC | 4 | 36617413 | 597 | exons | 026940:DAVISGALT002( | AVISGALT00269400 | AVISGALG0000269 | AVISGALG00002694 |
| DMC | 4 | 36617413 | 597 | exon1 | 026940:DAVISGALT002( | AVISGALT00269400 | AVISGALG0000269 | AVISGALG00002694 |
| DMC | 4 | 36617413 | 597 | downstream | 1313:TAGAGALT00000( | GAGALT00000327 | GAGALG000000061 | GAGALG0000000613 |
| DMC | 4 | 37458757 | -3071 | exons | 027058:DAVISGALT002 | AVISGALT00270580 | AVISGALG0000270 | AVISGALG00002705 |
| DMC | 4 | 37458757 | -3071 | exon1 | 027058:DAVISGALT002 | AVISGALT00270580 | AVISGALG0000270 | AVISGALG00002705 |
| DMC | 4 | 37458757 | -3071 | introns | 865:NM_001398055:int | NM_001398055 | LEF1 | LOC395865 |
| DMC | 4 | 37458757 | -3071 | introns | 865:NM_001398056:int | NM_001398056 | LEF1 | LOC395865 |
| DMC | 4 | 37458757 | -3071 | introns | 865:NM_001398057:in | NM_001398057 | LEF1 | LOC395865 |
| DMC | 4 | 37458757 | -3071 | introns | 865:NM_001398058:int | NM_001398058 | LEF1 | LOC395865 |
| DMC | 4 | 37458757 | -3071 | introns | 5865:NM_205013:intro | NM_205013 | LEF1 | LOC395865 |
| DMC | 4 | 37460974 | 3395 | exons | 027058:DAVISGALT002 | AVISGALT00270580 | AVISGALG0000270 | AVISGALG00002705 |
| DMC | 4 | 37460974 | 3395 | exon1 | 027058:DAVISGALT002 | AVISGALT00270580 | AVISGALG0000270 | AVISGALG00002705 |
| DMC | 4 | 37460974 | 3395 | introns | 865:NM_001398055:int | NM_001398055 | LEF1 | LOC395865 |
| DMC | 4 | 37460974 | 3395 | introns | 865:NM_001398056:int | NM_001398056 | LEF1 | LOC395865 |
| DMC | 4 | 37460974 | 3395 | introns | 865:NM_001398057:in | NM_001398057 | LEF1 | LOC395865 |
| DMC | 4 | 37460974 | 3395 | introns | 865:NM_001398058:int | NM_001398058 | LEF1 | LOC395865 |
| DMC | 4 | 37460974 | 3395 | introns | 5865:NM_205013:intro | NM_205013 | LEF1 | LOC395865 |
| DMC | 4 | 90662682 | -47 | introns | 47944:XM_025150449: | XM_025150449 | LOC101747944 | LOC101747944 |
| DMC | 5 | 715156 | -635 | exons | 053358:XR_001466523: | XR_001466523 | LOC107053358 | LOC107053358 |
| DMC | 5 | 715156 | -635 | exon1 | 053358:XR_001466523: | XR_001466523 | LOC107053358 | LOC107053358 |
| DMC | 5 | 715156 | -635 | introns | 081:XM_015286284:int | XM_015286284 | EPS8L2 | LOC770081 |
| DMC | 5 | 715156 | -635 | intron1 | 081:XM_015286284:int | XM_015286284 | EPS8L2 | LOC770081 |
| DMC | 5 | 715156 | -635 | downstream | 9748:DAVISGALT00297 | AVISGALT00297480 | AVISGALG0000297 | AVISGALG00002974 |
| DMC | 5 | 9096345 | -582 | promoter | 18386:XR_006939440: | XR_006939440 | LOC124418386 | LOC124418386 |
| DMC | 5 | 9096345 | -582 | introns | 048:XM_040701185:int | XM_040701185 | DENND5A | LOC423048 |
| DMC | 5 | 9096345 | -582 | introns | 048:XM_015286468:int | XM_015286468 | DENND5A | LOC423048 |
| DMC | 5 | 9096345 | -582 | introns | 048:XM_015286469:int | XM_015286469 | DENND5A | LOC423048 |
| DMC | 5 | 9096345 | -582 | introns | 048:XM_015286470:int | XM_015286470 | DENND5A | LOC423048 |
| DMC | 5 | 9096345 | -582 | introns | 048:XM_015286472:int | XM_015286472 | DENND5A | LOC423048 |
| DMC | 5 | 9096345 | -582 | intron1 | 048:XM_040701185:int | XM_040701185 | DENND5A | LOC423048 |
| DMC | 5 | 9096345 | -582 | intron1 | 048:XM_015286468:int | XM_015286468 | DENND5A | LOC423048 |
| DMC | 5 | 9096345 | -582 | intron1 | 048:XM_015286469:int | XM_015286469 | DENND5A | LOC423048 |
| DMC | 5 | 9096345 | -582 | intron1 | 048:XM_015286470:int | XM_015286470 | DENND5A | LOC423048 |
| DMC | 5 | 9096345 | -582 | intron1 | 048:XM_015286472:int | XM_015286472 | DENND5A | LOC423048 |
| DMC | 5 | 9160003 | 54 | introns | 049:XM_015286476:int | XM_015286476 | SCUBE2 | LOC423049 |
| DMC | 5 | 9160003 | 54 | introns | 049:XM_015286474:int | XM_015286474 | SCUBE2 | LOC423049 |
| DMC | 5 | 9160003 | 54 | introns | 049:XM_015286475:int | XM_015286475 | SCUBE2 | LOC423049 |
| DMC | 5 | 9160003 | 54 | introns | 049:XM_015286477:int | XM_015286477 | SCUBE2 | LOC423049 |
| DMC | 5 | 9160003 | 54 | introns | 049:XM_040701526:int | XM_040701526 | SCUBE2 | LOC423049 |
| DMC | 5 | 9160003 | 54 | introns | 049:XR_001466551:intr | XR_001466551 | SCUBE2 | LOC423049 |
| DMC | 5 | 9160003 | 54 | introns | 049:XR_005860192:intr | XR_005860192 | SCUBE2 | LOC423049 |
| DMC | 5 | 9160003 | 54 | introns | 23049:XM_420982:intro | XM_420982 | SCUBE2 | LOC423049 |
| DMC | 5 | 9160003 | 54 | intron1 | 049:XM_015286476:int | XM_015286476 | SCUBE2 | LOC423049 |
| DMC | 5 | 9160003 | 54 | intron1 | 049:XM_015286474:int | XM_015286474 | SCUBE2 | LOC423049 |
| DMC | 5 | 9160003 | 54 | intron1 | 049:XM_015286475:int | XM_015286475 | SCUBE2 | LOC423049 |
| DMC | 5 | 9160003 | 54 | intron1 | 049:XM_015286477:int | XM_015286477 | SCUBE2 | LOC423049 |
| DMC | 5 | 9160003 | 54 | intron1 | 049:XM_040701526:int | XM_040701526 | SCUBE2 | LOC423049 |
| DMC | 5 | 9160003 | 54 | intron1 | 049:XR_001466551:intr | XR_001466551 | SCUBE2 | LOC423049 |
| DMC | 5 | 9160003 | 54 | intron1 | 049:XR_005860192:intr | XR_005860192 | SCUBE2 | LOC423049 |
| DMC | 5 | 9160003 | 54 | intron1 | 23049:XM_420982:intro | XM_420982 | SCUBE2 | LOC423049 |
| DMC | 5 | 9207733 | -1781 | introns | 23050:XM_420983:intr | XM_420983 | NRIP2 | LOC423050 |
| DMC | 5 | 9207733 | -1781 | intron1 | 23050:XM_420983:intr | XM_420983 | NRIP2 | LOC423050 |
| DMC | 5 | 9380651 | -1416 | introns | 055:XM_004941391:intr | XM_004941391 | DENND2B | LOC423055 |
| DMC | 5 | 9380651 | -1416 | introns | 055:XM_015286487:intr | XM_015286487 | DENND2B | LOC423055 |
| DMC | 5 | 9380651 | -1416 | introns | 055:XM_046941985:intr | XM_046941985 | DENND2B | LOC423055 |
| DMC | 5 | 9380651 | -1416 | introns | 055:XM_046941986:intr | XM_046941986 | DENND2B | LOC423055 |
| DMC | 5 | 9380651 | -1416 | introns | 055:XM_015286486:intr | XM_015286486 | DENND2B | LOC423055 |
| DMC | 5 | 9380651 | -1416 | introns | 055:XM_015286490:intr | XM_015286490 | DENND2B | LOC423055 |
| DMC | 5 | 9380651 | -1416 | introns | 055:XM_040701334:intr | XM_040701334 | DENND2B | LOC423055 |
| DMC | 5 | 9380651 | -1416 | introns | 055:XM_040701337:intr | XM_040701337 | DENND2B | LOC423055 |
| DMC | 5 | 9380651 | -1416 | introns | 055:XM_046941980:intr | XM_046941980 | DENND2B | LOC423055 |
| DMC | 5 | 9380651 | -1416 | introns | 055:XM_046941981:intr | XM_046941981 | DENND2B | LOC423055 |
| DMC | 5 | 9380651 | -1416 | introns | 055:XM_046941983:intr | XM_046941983 | DENND2B | LOC423055 |
| DMC | 5 | 9380651 | -1416 | introns | 055:XM_046941988:intr | XM_046941988 | DENND2B | LOC423055 |
| DMC | 5 | 9380651 | -1416 | introns | 055:XM_046941989:intr | XM_046941989 | DENND2B | LOC423055 |

| | | | | | | | | |
|---|---|---|---|---|---|---|---|---|
| DMC | 5 | 9380651 | -1416 | introns | 3055:XM_420988:intro | XM_420988 | DENND2B | LOC423055 |
| DMC | 5 | 9380651 | -1416 | introns | 055:XM_046941982:intr | XM_046941982 | DENND2B | LOC423055 |
| DMC | 5 | 9380651 | -1416 | introns | 055:XM_004941390:intr | XM_004941390 | DENND2B | LOC423055 |
| DMC | 5 | 9380651 | -1416 | introns | 055:XM_025150880:intr | XM_025150880 | DENND2B | LOC423055 |
| DMC | 5 | 9380651 | -1416 | introns | 055:XM_040701336:intr | XM_040701336 | DENND2B | LOC423055 |
| DMC | 5 | 9380651 | -1416 | introns | 055:XM_046941979:intr | XM_046941979 | DENND2B | LOC423055 |
| DMC | 5 | 9380651 | -1416 | introns | 055:XM_046941984:intr | XM_046941984 | DENND2B | LOC423055 |
| DMC | 5 | 9380651 | -1416 | introns | 055:XM_015286489:intr | XM_015286489 | DENND2B | LOC423055 |
| DMC | 5 | 9380651 | -1416 | introns | 055:XM_046941990:intr | XM_046941990 | DENND2B | LOC423055 |
| DMC | 5 | 9380651 | -1416 | introns | 055:XM_046941987:intr | XM_046941987 | DENND2B | LOC423055 |
| DMC | 5 | 9380651 | -1416 | introns | 055:XM_046941992:intr | XM_046941992 | DENND2B | LOC423055 |
| DMC | 5 | 9380651 | -1416 | introns | 055:XM_040701335:intr | XM_040701335 | DENND2B | LOC423055 |
| DMC | 5 | 9380651 | -1416 | introns | 055:XM_040701339:intr | XM_040701339 | DENND2B | LOC423055 |
| DMC | 5 | 15080911 | -2286 | introns | 3104:XM_046942326:in | XM_046942326 | TSPAN4 | LOC423104 |
| DMC | 5 | 15080911 | -2286 | introns | 3104:XM_046942330:in | XM_046942330 | TSPAN4 | LOC423104 |
| DMC | 5 | 15080911 | -2286 | introns | 104:XM_015286698:int | XM_015286698 | TSPAN4 | LOC423104 |
| DMC | 5 | 15080911 | -2286 | introns | 104:XM_040701678:int | XM_040701678 | TSPAN4 | LOC423104 |
| DMC | 5 | 15080911 | -2286 | introns | 104:XM_040701677:int | XM_040701677 | TSPAN4 | LOC423104 |
| DMC | 5 | 15080911 | -2286 | introns | 104:XM_046942329:int | XM_046942329 | TSPAN4 | LOC423104 |
| DMC | 5 | 15080911 | -2286 | introns | 104:XM_015286700:int | XM_015286700 | TSPAN4 | LOC423104 |
| DMC | 5 | 15080911 | -2286 | introns | 3104:XM_040701680:in | XM_040701680 | TSPAN4 | LOC423104 |
| DMC | 5 | 15080911 | -2286 | introns | 104:XM_015286697:int | XM_015286697 | TSPAN4 | LOC423104 |
| DMC | 5 | 15080911 | -2286 | introns | 3104:XM_040701679:in | XM_040701679 | TSPAN4 | LOC423104 |
| DMC | 5 | 15080911 | -2286 | introns | 104:XM_046942327:int | XM_046942327 | TSPAN4 | LOC423104 |
| DMC | 5 | 15080911 | -2286 | introns | 104:XM_015286696:int | XM_015286696 | TSPAN4 | LOC423104 |
| DMC | 5 | 15080911 | -2286 | introns | 3104:XM_015286701:in | XM_015286701 | TSPAN4 | LOC423104 |
| DMC | 5 | 15080911 | -2286 | introns | 104:XM_046942328:int | XM_046942328 | TSPAN4 | LOC423104 |
| DMC | 5 | 15080911 | -2286 | introns | 104:XM_046942331:int | XM_046942331 | TSPAN4 | LOC423104 |
| DMC | 5 | 15080911 | -2286 | intron1 | 3104:XM_046942326:in | XM_046942326 | TSPAN4 | LOC423104 |
| DMC | 5 | 15080911 | -2286 | downstream | 0622:DAVISGALT00306 | DAVISGALT00306220 | DAVISGALG0000306 | DAVISGALG00003062 |
| DMC | 5 | 55342231 | -1919 | introns | 1752106:XR_210740:int | XR_210740 | LOC101752106 | LOC101752106 |
| DMC | 5 | 55342231 | -1919 | intron1 | 1752106:XR_210740:int | XR_210740 | LOC101752106 | LOC101752106 |
| DMC | 6 | 17819124 | -28339 | introns | 49795:XR_005861100:in | XR_005861100 | LOC101749795 | LOC101749795 |
| DMC | 6 | 17819124 | -28339 | introns | 49795:XR_005861103:in | XR_005861103 | LOC101749795 | LOC101749795 |
| DMC | 6 | 17819124 | -28339 | introns | 49795:XR_005861104:in | XR_005861104 | LOC101749795 | LOC101749795 |
| DMC | 6 | 17819124 | -28339 | introns | 49795:XR_005861105:in | XR_005861105 | LOC101749795 | LOC101749795 |
| DMC | 6 | 17819124 | -28339 | introns | 49795:XR_005861107:in | XR_005861107 | LOC101749795 | LOC101749795 |
| DMC | 6 | 33988936 | -37 | exons | 58023:XM_046943089: | XM_046943089 | NPS | LOC100858023 |
| DMC | 6 | 33988936 | -37 | UTR3 | 8023:XM_046943089:3' | XM_046943089 | NPS | LOC100858023 |
| DMC | 7 | 3140595 | 50 | exons | 671:XM_040703313:exo | XM_040703313 | ERBB4 | LOC395671 |
| DMC | 7 | 3140595 | 50 | exons | 671:XM_040703314:exo | XM_040703314 | ERBB4 | LOC395671 |
| DMC | 7 | 3140595 | 50 | exons | 671:XM_046943423:exo | XM_046943423 | ERBB4 | LOC395671 |
| DMC | 7 | 3140595 | 50 | exons | 671:XM_015289115:exo | XM_015289115 | ERBB4 | LOC395671 |
| DMC | 7 | 3140595 | 50 | exons | 671:XM_015289117:exo | XM_015289117 | ERBB4 | LOC395671 |
| DMC | 7 | 3140595 | 50 | exons | 671:XM_015289118:exo | XM_015289118 | ERBB4 | LOC395671 |
| DMC | 7 | 3140595 | 50 | exons | 671:XM_040703312:exo | XM_040703312 | ERBB4 | LOC395671 |
| DMC | 7 | 3140595 | 50 | exons | 571:NM_001030365:exo | NM_001030365 | ERBB4 | LOC395671 |
| DMC | 7 | 3140595 | 50 | cds | 671:XM_040703313:exo | XM_040703313 | ERBB4 | LOC395671 |
| DMC | 7 | 3140595 | 50 | cds | 671:XM_040703314:exo | XM_040703314 | ERBB4 | LOC395671 |
| DMC | 7 | 3140595 | 50 | cds | 671:XM_046943423:exo | XM_046943423 | ERBB4 | LOC395671 |
| DMC | 7 | 3140595 | 50 | cds | 671:XM_015289115:exo | XM_015289115 | ERBB4 | LOC395671 |
| DMC | 7 | 3140595 | 50 | cds | 671:XM_015289117:exo | XM_015289117 | ERBB4 | LOC395671 |
| DMC | 7 | 3140595 | 50 | cds | 671:XM_015289118:exo | XM_015289118 | ERBB4 | LOC395671 |
| DMC | 7 | 3140595 | 50 | cds | 671:XM_040703312:exo | XM_040703312 | ERBB4 | LOC395671 |
| DMC | 7 | 3140595 | 50 | cds | 571:NM_001030365:exo | NM_001030365 | ERBB4 | LOC395671 |
| DMC | 7 | 27210934 | -2187 | promoter | 036642:DAVISGALT0036 | DAVISGALT00366420 | DAVISGALG0000366 | DAVISGALG00003664 |
| DMC | 7 | 27210934 | -2187 | introns | 253:XM_015289974:int | XM_015289974 | KALRN | LOC424253 |
| DMC | 7 | 27210934 | -2187 | introns | 253:XM_015289984:int | XM_015289984 | KALRN | LOC424253 |
| DMC | 7 | 27210934 | -2187 | introns | 253:XM_046944184:int | XM_046944184 | KALRN | LOC424253 |
| DMC | 7 | 27210934 | -2187 | introns | 253:XM_046944185:int | XM_046944185 | KALRN | LOC424253 |
| DMC | 7 | 27210934 | -2187 | introns | 253:XM_040703549:int | XM_040703549 | KALRN | LOC424253 |
| DMC | 7 | 27210934 | -2187 | introns | 253:XM_046944183:int | XM_046944183 | KALRN | LOC424253 |
| DMC | 7 | 27210934 | -2187 | introns | 253:XM_040703548:int | XM_040703548 | KALRN | LOC424253 |
| DMC | 7 | 27210934 | -2187 | introns | 253:XM_040703550:int | XM_040703550 | KALRN | LOC424253 |
| DMC | 7 | 27210934 | -2187 | introns | 253:XM_015289976:int | XM_015289976 | KALRN | LOC424253 |
| DMC | 7 | 27210934 | -2187 | introns | 253:XM_015289977:int | XM_015289977 | KALRN | LOC424253 |
| DMC | 7 | 27210934 | -2187 | introns | 253:XM_015289978:int | XM_015289978 | KALRN | LOC424253 |

| | | | | | | | | |
|---|---|---|---|---|---|---|---|---|
| DMC | 7 | 27210934 | -2187 | introns | 253:XM_015289980:int | XM_015289980 | KALRN | LOC424253 |
| DMC | 7 | 27210934 | -2187 | introns | 253:XM_015289981:int | XM_015289981 | KALRN | LOC424253 |
| DMC | 7 | 27210934 | -2187 | introns | 253:XM_015289982:int | XM_015289982 | KALRN | LOC424253 |
| DMC | 7 | 27210934 | -2187 | introns | 253:XM_040703547:int | XM_040703547 | KALRN | LOC424253 |
| DMC | 8 | 2258778 | 10934 | other | NA | NA | NA | NA |
| DMC | 8 | 3707756 | 32 | promoter | 4363:XM_015290397:pr | XM_015290397 | HTATIP2 | LOC424363 |
| DMC | 8 | 3707756 | 32 | promoter | 24363:XM_422206:pror | XM_422206 | HTATIP2 | LOC424363 |
| DMC | 8 | 3707756 | 32 | exons | '51271:XM_004943133 | XM_004943133 | LOC101751271 | LOC101751271 |
| DMC | 8 | 3707756 | 32 | exons | '51271:XM_015290395 | XM_015290395 | LOC101751271 | LOC101751271 |
| DMC | 8 | 3707756 | 32 | exons | 751271:XR_001467726: | XR_001467726 | LOC101751271 | LOC101751271 |
| DMC | 8 | 3707756 | 32 | exons | 01751271:XR_212726:ex | XR_212726 | LOC101751271 | LOC101751271 |
| DMC | 8 | 3707756 | 32 | exon1 | '51271:XM_004943133 | XM_004943133 | LOC101751271 | LOC101751271 |
| DMC | 8 | 3707756 | 32 | exon1 | '51271:XM_015290395 | XM_015290395 | LOC101751271 | LOC101751271 |
| DMC | 8 | 3707756 | 32 | exon1 | 751271:XR_001467726: | XR_001467726 | LOC101751271 | LOC101751271 |
| DMC | 8 | 3707756 | 32 | exon1 | 01751271:XR_212726:ex | XR_212726 | LOC101751271 | LOC101751271 |
| DMC | 8 | 3707756 | 32 | cds | '51271:XM_004943133 | XM_004943133 | LOC101751271 | LOC101751271 |
| DMC | 8 | 3707756 | 32 | cds | '51271:XM_015290395 | XM_015290395 | LOC101751271 | LOC101751271 |
| DMC | 8 | 3794822 | -565 | promoter | 024649:TAGAGALT0000( | GAGALT00000040200 | GAGALG000000024 | GAGALG0000000246 |
| DMC | 8 | 3794822 | -565 | introns | 24366:XM_422209:intr | XM_422209 | OLFML2B | LOC424366 |
| DMC | 8 | 5164023 | -3340 | introns | '133:XM_015290334:in | XM_015290334 | LMX1A | LOC777133 |
| DMC | 8 | 5164023 | -3340 | introns | '133:NM_001257388:in | NM_001257388 | LMX1A | LOC777133 |
| DMC | 8 | 5183625 | -830 | exons | 7133:XM_015290334:e | XM_015290334 | LMX1A | LOC777133 |
| DMC | 8 | 5183625 | -830 | exons | 7133:NM_001257388:e | NM_001257388 | LMX1A | LOC777133 |
| DMC | 8 | 5183625 | -830 | UTR3 | 33:XM_015290334:3'UT | XM_015290334 | LMX1A | LOC777133 |
| DMC | 8 | 5183625 | -830 | UTR3 | 33:NM_001257388:3'U | NM_001257388 | LMX1A | LOC777133 |
| DMC | 8 | 14412257 | -296 | exons | 418408:XR_006939995: | XR_006939995 | LOC124418408 | LOC124418408 |
| DMC | 8 | 15983726 | 10820 | introns | 898:NM_001396492:in | NM_001396492 | DDAH1 | LOC378898 |
| DMC | 8 | 15983726 | 10820 | intron1 | 898:NM_001396492:in | NM_001396492 | DDAH1 | LOC378898 |
| DMC | 8 | 24968182 | -55 | exons | 4664:XM_025153036:ex | XM_025153036 | PCSK9 | LOC424664 |
| DMC | 8 | 24968182 | -55 | cds | 4664:XM_025153036:ex | XM_025153036 | PCSK9 | LOC424664 |
| DMC | 8 | 24968187 | -60 | exons | 4664:XM_025153036:ex | XM_025153036 | PCSK9 | LOC424664 |
| DMC | 8 | 24968187 | -60 | cds | 4664:XM_025153036:ex | XM_025153036 | PCSK9 | LOC424664 |
| DMC | 8 | 25383827 | 172 | promoter | 024321:TAGAGALT0000( | GAGALT00000041250 | GAGALG000000024 | GAGALG0000000243 |
| DMC | 8 | 25383827 | 172 | promoter | 024321:TAGAGALT0000( | GAGALT00000041250 | GAGALG000000024 | GAGALG0000000243 |
| DMC | 8 | 25383827 | 172 | promoter | 057766:TAGAGALT0000( | GAGALT00000041250 | GAGALG000000057 | GAGALG0000000577 |
| DMC | 8 | 25383827 | 172 | exons | 4666:NM_001130488:e | NM_001130488 | PLPP3 | LOC424666 |
| DMC | 8 | 25383827 | 172 | UTR3 | 56:NM_001130488:3'U | NM_001130488 | PLPP3 | LOC424666 |
| DMC | 8 | 25383827 | 172 | downstream | 4033:XR_005862074:dc | XR_005862074 | LOC107054033 | LOC107054033 |
| DMC | 8 | 28105010 | -440 | introns | 700:XM_015291029:int | XM_015291029 | PDE4B | LOC424700 |
| DMC | 8 | 28105010 | -440 | introns | 700:XM_046944521:int | XM_046944521 | PDE4B | LOC424700 |
| DMC | 8 | 28105010 | -440 | introns | 700:XM_015291030:int | XM_015291030 | PDE4B | LOC424700 |
| DMC | 8 | 28105010 | -440 | intron1 | 700:XM_015291030:int | XM_015291030 | PDE4B | LOC424700 |
| DMC | 9 | 782408 | -89 | exons | 745:XM_025153529:exo | XM_025153529 | INPP5D | LOC424745 |
| DMC | 9 | 782408 | -89 | cds | 745:XM_025153529:exo | XM_025153529 | INPP5D | LOC424745 |
| DMC | 9 | 847676 | -2321 | promoter | 33001:XR_005862332:p | XR_005862332 | LOC112533001 | LOC112533001 |
| DMC | 9 | 847676 | -2321 | promoter | 33001:XR_005862333:p | XR_005862333 | LOC112533001 | LOC112533001 |
| DMC | 9 | 847676 | -2321 | introns | 747:XM_015277197:int | XM_015277197 | DGKD | LOC424747 |
| DMC | 9 | 847676 | -2321 | introns | 747:XM_040706007:int | XM_040706007 | DGKD | LOC424747 |
| DMC | 9 | 847676 | -2321 | intron1 | 747:XM_015277197:int | XM_015277197 | DGKD | LOC424747 |
| DMC | 9 | 847676 | -2321 | intron1 | 747:XM_040706007:int | XM_040706007 | DGKD | LOC424747 |
| DMC | 9 | 5375386 | 4632 | introns | 826:XM_046898396:int | XM_046898396 | PIK3CB | LOC424826 |
| DMC | 9 | 5375386 | 4632 | intron1 | 826:XM_046898396:int | XM_046898396 | PIK3CB | LOC424826 |
| DMC | 9 | 15547353 | -66 | promoter | 47338:XM_015291556: | XM_015291556 | LOC101747338 | LOC101747338 |
| DMC | 9 | 15547353 | -66 | promoter | 47338:XM_015291557: | XM_015291557 | LOC101747338 | LOC101747338 |
| DMC | 9 | 15547353 | -66 | exons | 950:XM_040705689:exo | XM_040705689 | ABCF3 | LOC424950 |
| DMC | 9 | 15547353 | -66 | exons | 24950:XM_422757:exor | XM_422757 | ABCF3 | LOC424950 |
| DMC | 9 | 15547353 | -66 | UTR3 | 50:XM_040705689:3'UT | XM_040705689 | ABCF3 | LOC424950 |
| DMC | 9 | 15547353 | -66 | UTR3 | 950:XM_422757:3'UTR( | XM_422757 | ABCF3 | LOC424950 |
| DMC | 9 | 15558108 | -102 | promoter | 039727:DAVISGALT0039 | AVISGALT00397270 | AVISGALG00003972 | AVISGALG00003972 |
| DMC | 9 | 15558108 | -102 | exons | 4951:XM_040706135:ex | XM_040706135 | VWA5B2 | LOC424951 |
| DMC | 9 | 15558108 | -102 | cds | 4951:XM_040706135:ex | XM_040706135 | VWA5B2 | LOC424951 |
| DMC | 9 | 15558965 | -83 | promoter | 039727:DAVISGALT0039 | AVISGALT00397270 | AVISGALG00003972 | AVISGALG00003972 |
| DMC | 9 | 15558965 | -83 | exons | 4951:XM_040706135:ex | XM_040706135 | VWA5B2 | LOC424951 |
| DMC | 9 | 15558965 | -83 | cds | 4951:XM_040706135:ex | XM_040706135 | VWA5B2 | LOC424951 |
| DMC | 9 | 15559065 | 64 | promoter | 039727:DAVISGALT0039 | AVISGALT00397270 | AVISGALG00003972 | AVISGALG00003972 |
| DMC | 9 | 15559065 | 64 | introns | 951:XM_040706135:int | XM_040706135 | VWA5B2 | LOC424951 |
| DMC | 9 | 15559300 | 46 | promoter | 039727:DAVISGALT0039 | AVISGALT00397270 | AVISGALG00003972 | AVISGALG00003972 |

| | | | | | | | | |
|---|---|---|---|---|---|---|---|---|
| DMC | 9 | 15559300 | 46 | introns | 951:XM_040706135:int | XM_040706135 | VWA5B2 | LOC424951 |
| DMC | 9 | 15559797 | 14 | promoter | 039727:DAVISGALT0039 | DAVISGALT00397270 | AVISGALG00003972 | DAVISGALG00003972 |
| DMC | 9 | 15559797 | 14 | introns | 951:XM_040706135:intr | XM_040706135 | VWA5B2 | LOC424951 |
| DMC | 9 | 15564193 | 146 | promoter | 1092:XM_040705700:pr | XM_040705700 | LOC431092 | LOC431092 |
| DMC | 9 | 15564193 | 146 | promoter | 1092:XM_046898849:pr | XM_046898849 | LOC431092 | LOC431092 |
| DMC | 9 | 15564193 | 146 | exons | 951:XM_040706135:ex | XM_040706135 | VWA5B2 | LOC424951 |
| DMC | 9 | 15564193 | 146 | cds | 951:XM_040706135:ex | XM_040706135 | VWA5B2 | LOC424951 |
| DMC | 9 | 15564193 | 146 | downstream | 52:XM_004943402:dov | XM_004943402 | ALG3 | LOC424952 |
| DMC | 9 | 15564193 | 146 | downstream | 4952:XM_422759:down | XM_422759 | ALG3 | LOC424952 |
| DMC | 9 | 15588919 | -137 | promoter | 57430:XM_004943409: | XM_004943409 | ECE2 | LOC100857430 |
| DMC | 9 | 15588919 | -137 | promoter | 57430:XM_025153673: | XM_025153673 | ECE2 | LOC100857430 |
| DMC | 9 | 15588919 | -137 | promoter | 57430:XM_025153674: | XM_025153674 | ECE2 | LOC100857430 |
| DMC | 9 | 15588919 | -137 | promoter | 57430:XR_001468044:p | XR_001468044 | ECE2 | LOC100857430 |
| DMC | 9 | 15588919 | -137 | exons | 294:NM_001012934:ex | NM_001012934 | PSMD2 | LOC425294 |
| DMC | 9 | 15588919 | -137 | cds | 294:NM_001012934:ex | NM_001012934 | PSMD2 | LOC425294 |
| DMC | 10 | 1621489 | -140 | introns | 456:XM_046898944:int | XM_046898944 | PKLR | LOC396456 |
| DMC | 10 | 1621489 | -140 | introns | 96456:NM_205469:intro | NM_205469 | PKLR | LOC396456 |
| DMC | 10 | 1621489 | -140 | introns | 456:XM_015278795:int | XM_015278795 | PKLR | LOC396456 |
| DMC | 10 | 1621489 | -140 | introns | 456:XM_015278796:int | XM_015278796 | PKLR | LOC396456 |
| DMC | 10 | 1621489 | -140 | introns | 456:XM_025153808:int | XM_025153808 | PKLR | LOC396456 |
| DMC | 10 | 1621489 | -140 | introns | 456:XM_046898945:int | XM_046898945 | PKLR | LOC396456 |
| DMC | 10 | 1621489 | -140 | intron1 | 456:XM_046898944:int | XM_046898944 | PKLR | LOC396456 |
| DMC | 10 | 1621489 | -140 | intron1 | 96456:NM_205469:intro | NM_205469 | PKLR | LOC396456 |
| DMC | 10 | 1621489 | -140 | intron1 | 456:XM_015278795:int | XM_015278795 | PKLR | LOC396456 |
| DMC | 10 | 1621489 | -140 | intron1 | 456:XM_015278796:int | XM_015278796 | PKLR | LOC396456 |
| DMC | 10 | 1621489 | -140 | intron1 | 456:XM_025153808:int | XM_025153808 | PKLR | LOC396456 |
| DMC | 10 | 1621489 | -140 | intron1 | 456:XM_046898945:int | XM_046898945 | PKLR | LOC396456 |
| DMC | 10 | 1633735 | 35 | introns | 472:XR_005862709:intr | XR_005862709 | PARP6 | LOC769472 |
| DMC | 10 | 1633735 | 35 | introns | 472:XR_005862710:intr | XR_005862710 | PARP6 | LOC769472 |
| DMC | 10 | 1633735 | 35 | introns | 472:XM_040706575:intr | XM_040706575 | PARP6 | LOC769472 |
| DMC | 10 | 1633735 | 35 | introns | 472:XM_015279056:intr | XM_015279056 | PARP6 | LOC769472 |
| DMC | 10 | 1633735 | 35 | introns | 472:XM_015279058:intr | XM_015279058 | PARP6 | LOC769472 |
| DMC | 10 | 1633735 | 35 | introns | 472:XM_015279059:intr | XM_015279059 | PARP6 | LOC769472 |
| DMC | 10 | 1633735 | 35 | introns | 472:XM_015279060:intr | XM_015279060 | PARP6 | LOC769472 |
| DMC | 10 | 1633735 | 35 | introns | 472:XM_040706567:intr | XM_040706567 | PARP6 | LOC769472 |
| DMC | 10 | 1633735 | 35 | introns | 472:XM_040706568:intr | XM_040706568 | PARP6 | LOC769472 |
| DMC | 10 | 1633735 | 35 | introns | 472:XM_040706569:intr | XM_040706569 | PARP6 | LOC769472 |
| DMC | 10 | 1633735 | 35 | introns | 472:XM_040706571:intr | XM_040706571 | PARP6 | LOC769472 |
| DMC | 10 | 1633735 | 35 | introns | 472:XM_040706574:intr | XM_040706574 | PARP6 | LOC769472 |
| DMC | 10 | 2627156 | -420 | exons | 9957:XM_015279034:e | XM_015279034 | ISLR | LOC769957 |
| DMC | 10 | 2627156 | -420 | exons | 9957:XM_015279036:e | XM_015279036 | ISLR | LOC769957 |
| DMC | 10 | 2627156 | -420 | exons | 9957:XM_015279035:e | XM_015279035 | ISLR | LOC769957 |
| DMC | 10 | 2627156 | -420 | exons | 9957:XM_001233272:e | XM_001233272 | ISLR | LOC769957 |
| DMC | 10 | 2627156 | -420 | cds | 9957:XM_015279034:e | XM_015279034 | ISLR | LOC769957 |
| DMC | 10 | 2627156 | -420 | cds | 9957:XM_015279036:e | XM_015279036 | ISLR | LOC769957 |
| DMC | 10 | 2627156 | -420 | cds | 9957:XM_015279035:e | XM_015279035 | ISLR | LOC769957 |
| DMC | 10 | 2627156 | -420 | cds | 9957:XM_001233272:e | XM_001233272 | ISLR | LOC769957 |
| DMC | 10 | 2648462 | -346 | promoter | 52152:XM_015279033: | XM_015279033 | LOC107052152 | LOC107052152 |
| DMC | 10 | 2648462 | -346 | exons | 857563:XM_004943699 | XM_004943699 | PMLL | LOC100857563 |
| DMC | 10 | 2648462 | -346 | exons | 857563:XM_040706616 | XM_040706616 | PMLL | LOC100857563 |
| DMC | 10 | 2648462 | -346 | cds | 857563:XM_004943699 | XM_004943699 | PMLL | LOC100857563 |
| DMC | 10 | 2648462 | -346 | cds | 857563:XM_040706616 | XM_040706616 | PMLL | LOC100857563 |
| DMC | 10 | 2648462 | -346 | downstream | 2183:TAGAGALT00000( | GAGALT0000000617 | GAGALG000000082 | GAGALG0000000821 |
| DMC | 10 | 17981868 | 134 | exons | 5537:XM_040706662:ex | XM_040706662 | IGDCC4 | LOC415537 |
| DMC | 10 | 17981868 | 134 | exons | 5537:XM_003641826:ex | XM_003641826 | IGDCC4 | LOC415537 |
| DMC | 10 | 17981868 | 134 | exons | 5537:XM_004943893:ex | XM_004943893 | IGDCC4 | LOC415537 |
| DMC | 10 | 17981868 | 134 | exons | 5537:XM_025154131:ex | XM_025154131 | IGDCC4 | LOC415537 |
| DMC | 10 | 17981868 | 134 | cds | 5537:XM_040706662:ex | XM_040706662 | IGDCC4 | LOC415537 |
| DMC | 10 | 17981868 | 134 | cds | 5537:XM_003641826:ex | XM_003641826 | IGDCC4 | LOC415537 |
| DMC | 10 | 17981868 | 134 | cds | 5537:XM_004943893:ex | XM_004943893 | IGDCC4 | LOC415537 |
| DMC | 10 | 17981868 | 134 | cds | 5537:XM_025154131:ex | XM_025154131 | IGDCC4 | LOC415537 |
| DMC | 11 | 5594013 | -146 | exons | 5446:XM_025154146:e | XM_025154146 | SALL1 | LOC395446 |
| DMC | 11 | 5594013 | -146 | exons | 5446:XM_040707004:e | XM_040707004 | SALL1 | LOC395446 |
| DMC | 11 | 5594013 | -146 | exons | 5446:XM_046899380:e | XM_046899380 | SALL1 | LOC395446 |
| DMC | 11 | 5594013 | -146 | exons | 395446:NM_204707:exo | NM_204707 | SALL1 | LOC395446 |
| DMC | 11 | 5594013 | -146 | exons | 5446:XM_046899381:e | XM_046899381 | SALL1 | LOC395446 |
| DMC | 11 | 5594013 | -146 | cds | 5446:XM_025154146:e | XM_025154146 | SALL1 | LOC395446 |

| | | | | | | | | |
|---|---|---|---|---|---|---|---|---|
| DMC | 11 | 5594013 | -146 | cds | 5446:XM_040707004:e | XM_040707004 | SALL1 | LOC395446 |
| DMC | 11 | 5594013 | -146 | cds | 5446:XM_046899380:e | XM_046899380 | SALL1 | LOC395446 |
| DMC | 11 | 5594013 | -146 | cds | 95446:NM_204707:exo | NM_204707 | SALL1 | LOC395446 |
| DMC | 11 | 5594013 | -146 | cds | 5446:XM_046899381:e | XM_046899381 | SALL1 | LOC395446 |
| DMC | 11 | 8542050 | 8467 | introns | 5761:XM_004944212:in | XM_004944212 | ZNF536 | LOC415761 |
| DMC | 11 | 8542050 | 8467 | introns | 5761:XM_040645957:in | XM_040645957 | ZNF536 | LOC415761 |
| DMC | 11 | 8542050 | 8467 | introns | 5761:XM_004944211:in | XM_004944211 | ZNF536 | LOC415761 |
| DMC | 11 | 8542050 | 8467 | introns | 5761:XM_015292488:in | XM_015292488 | ZNF536 | LOC415761 |
| DMC | 11 | 8542050 | 8467 | introns | 5761:XM_004944214:in | XM_004944214 | ZNF536 | LOC415761 |
| DMC | 11 | 8542050 | 8467 | introns | 5761:XM_040645958:in | XM_040645958 | ZNF536 | LOC415761 |
| DMC | 11 | 8542050 | 8467 | introns | 5761:XM_040645959:in | XM_040645959 | ZNF536 | LOC415761 |
| DMC | 11 | 8542050 | 8467 | introns | 5761:XM_040645956:in | XM_040645956 | ZNF536 | LOC415761 |
| DMC | 11 | 18167101 | 42 | exons | 9028:XM_001232210:e | XM_001232210 | PABPN1L | LOC769028 |
| DMC | 11 | 18167101 | 42 | cds | 9028:XM_001232210:e | XM_001232210 | PABPN1L | LOC769028 |
| DMC | 11 | 18394188 | 48 | exons | 415852:XM_414212:exo | XM_414212 | LOC415852 | LOC415852 |
| DMC | 11 | 18394188 | 48 | exons | 5852:XM_040707208:e | XM_040707208 | LOC415852 | LOC415852 |
| DMC | 11 | 18394188 | 48 | UTR3 | 5852:XM_414212:3'UTR | XM_414212 | LOC415852 | LOC415852 |
| DMC | 11 | 18394188 | 48 | UTR3 | 52:XM_040707208:3'U | XM_040707208 | LOC415852 | LOC415852 |
| DMC | 11 | 19128275 | 8229 | introns | 682:XM_040707006:int | XM_040707006 | ZFHX3 | LOC395682 |
| DMC | 11 | 19128275 | 8229 | introns | 682:XM_046899382:int | XM_046899382 | ZFHX3 | LOC395682 |
| DMC | 11 | 19128275 | 8229 | introns | 682:XM_046899383:int | XM_046899383 | ZFHX3 | LOC395682 |
| DMC | 11 | 19128275 | 8229 | introns | 682:XM_046899384:int | XM_046899384 | ZFHX3 | LOC395682 |
| DMC | 11 | 19128275 | 8229 | introns | 5682:XM_040707009:in | XM_040707009 | ZFHX3 | LOC395682 |
| DMC | 11 | 19128275 | 8229 | intron1 | 5682:XM_040707009:in | XM_040707009 | ZFHX3 | LOC395682 |
| DMC | 11 | 19242844 | 2703 | promoter | 004095:INRAGALT00000 | IRAGALT000000086 | RAGALG00000004 | IRAGALG00000004 |
| DMC | 12 | 396767 | 551 | introns | 54342:XR_006931366:i | XR_006931366 | LOC107054342 | LOC107054342 |
| DMC | 12 | 396767 | 551 | introns | 54342:XR_006931367:i | XR_006931367 | LOC107054342 | LOC107054342 |
| DMC | 12 | 396767 | 551 | introns | 54342:XR_006931365:i | XR_006931365 | LOC107054342 | LOC107054342 |
| DMC | 12 | 396767 | 551 | introns | 028315:ENSGALT00010 | NSGALT0001006854 | NSGALG00010028 | NSGALG0001002831 |
| DMC | 12 | 396767 | 551 | introns | 028315:ENSGALT00010 | NSGALT0001006854 | NSGALG00010028 | NSGALG0001002831 |
| DMC | 12 | 396767 | 551 | introns | 028315:ENSGALT00010 | NSGALT0001006854 | NSGALG00010028 | NSGALG0001002831 |
| DMC | 12 | 396767 | 551 | downstream | 8317:ENSGALT0001006 | NSGALT0001006854 | NSGALG00010028 | NSGALG0001002831 |
| DMC | 12 | 1554669 | 467 | introns | .50:XM_015292785:intr | XM_015292785 | CACNA2D2 | LOC430150 |
| DMC | 12 | 1554669 | 467 | introns | .50:XM_015292786:intr | XM_015292786 | CACNA2D2 | LOC430150 |
| DMC | 12 | 1554669 | 467 | introns | .50:XM_015292787:intr | XM_015292787 | CACNA2D2 | LOC430150 |
| DMC | 12 | 1554669 | 467 | introns | .50:XM_040646449:intr | XM_040646449 | CACNA2D2 | LOC430150 |
| DMC | 12 | 1562135 | -547 | promoter | 001174:NONGGAT0020 | NONGGAT002058 | NONGGAG001174 | NONGGAG001174 |
| DMC | 12 | 1562135 | -547 | introns | 150:XM_015292785:int | XM_015292785 | CACNA2D2 | LOC430150 |
| DMC | 12 | 1562135 | -547 | introns | 150:XM_015292786:int | XM_015292786 | CACNA2D2 | LOC430150 |
| DMC | 12 | 1562135 | -547 | introns | 150:XM_015292787:int | XM_015292787 | CACNA2D2 | LOC430150 |
| DMC | 12 | 1562135 | -547 | introns | 150:XM_040646449:int | XM_040646449 | CACNA2D2 | LOC430150 |
| DMC | 12 | 1774382 | 831 | introns | 028237:ENSGALT00010 | NSGALT0001006842 | NSGALG00010028 | NSGALG0001002823 |
| DMC | 12 | 3134049 | 39 | exons | 5940:XM_040646196:e | XM_040646196 | TWF2 | LOC415940 |
| DMC | 12 | 3134049 | 39 | introns | 47452:XM_040646531: | XM_040646531 | LOC101747452 | LOC101747452 |
| DMC | 12 | 3134049 | 39 | introns | 940:XM_040646192:int | XM_040646192 | TWF2 | LOC415940 |
| DMC | 12 | 3134049 | 39 | introns | 940:NM_001030589:in | NM_001030589 | TWF2 | LOC415940 |
| DMC | 12 | 3134049 | 39 | introns | 940:XM_015292830:int | XM_015292830 | TWF2 | LOC415940 |
| DMC | 12 | 3134049 | 39 | introns | 940:XM_015292834:int | XM_015292834 | TWF2 | LOC415940 |
| DMC | 12 | 3134049 | 39 | introns | 940:XM_015292835:int | XM_015292835 | TWF2 | LOC415940 |
| DMC | 12 | 3134049 | 39 | introns | 940:XM_015292836:int | XM_015292836 | TWF2 | LOC415940 |
| DMC | 12 | 3134049 | 39 | introns | 940:XM_040646193:int | XM_040646193 | TWF2 | LOC415940 |
| DMC | 12 | 3134049 | 39 | introns | 5940:XM_040646195:in | XM_040646195 | TWF2 | LOC415940 |
| DMC | 12 | 3134049 | 39 | intron1 | 47452:XM_040646531: | XM_040646531 | LOC101747452 | LOC101747452 |
| DMC | 12 | 3134049 | 39 | UTR3 | 40:XM_040646196:3'U | XM_040646196 | TWF2 | LOC415940 |
| DMC | 12 | 3142614 | 49 | promoter | 5940:XM_040646192:pr | XM_040646192 | TWF2 | LOC415940 |
| DMC | 12 | 3142614 | 49 | UTR5 | 40:XM_040646192:5'U | XM_040646192 | TWF2 | LOC415940 |
| DMC | 12 | 3142614 | 49 | exons | 5940:XM_040646192:ex | XM_040646192 | TWF2 | LOC415940 |
| DMC | 12 | 3142614 | 49 | exon1 | 5940:XM_040646192:ex | XM_040646192 | TWF2 | LOC415940 |
| DMC | 12 | 3142614 | 49 | introns | 47452:XM_040646531: | XM_040646531 | LOC101747452 | LOC101747452 |
| DMC | 12 | 3142614 | 49 | introns | 940:NM_001030589:in | NM_001030589 | TWF2 | LOC415940 |
| DMC | 12 | 3142614 | 49 | introns | 940:XM_015292830:int | XM_015292830 | TWF2 | LOC415940 |
| DMC | 12 | 3142614 | 49 | introns | 940:XM_015292834:int | XM_015292834 | TWF2 | LOC415940 |
| DMC | 12 | 3142614 | 49 | introns | 940:XM_015292835:int | XM_015292835 | TWF2 | LOC415940 |
| DMC | 12 | 3142614 | 49 | introns | 940:XM_015292836:int | XM_015292836 | TWF2 | LOC415940 |
| DMC | 12 | 3142614 | 49 | introns | 940:XM_040646193:int | XM_040646193 | TWF2 | LOC415940 |
| DMC | 12 | 3142614 | 49 | introns | 5940:XM_040646195:in | XM_040646195 | TWF2 | LOC415940 |
| DMC | 12 | 3142614 | 49 | introns | 5940:XM_040646196:in | XM_040646196 | TWF2 | LOC415940 |

| | | | | | | | | |
|---|---|---|---|---|---|---|---|---|
| DMC | 12 | 3142614 | 49 | intron1 | 47452:XM_040646531: | XM_040646531 | LOC101747452 | LOC101747452 |
| DMC | 12 | 3142614 | 49 | intron1 | 940:NM_001030589:in | NM_001030589 | TWF2 | LOC415940 |
| DMC | 12 | 3142614 | 49 | intron1 | 940:XM_040646193:int | XM_040646193 | TWF2 | LOC415940 |
| DMC | 12 | 3316417 | 30 | promoter | 56877:XM_015301851: | XM_015301851 | LOC107056877 | LOC107056877 |
| DMC | 12 | 3316417 | 30 | promoter | 58836:XM_015292854: | XM_015292854 | ACY1 | LOC100858836 |
| DMC | 12 | 3316417 | 30 | promoter | 58836:XM_046900131: | XM_046900131 | ACY1 | LOC100858836 |
| DMC | 12 | 3316417 | 30 | exons | 358554:XM_015292858 | XM_015292858 | ABHD14A | LOC100858554 |
| DMC | 12 | 3316417 | 30 | introns | 47452:XM_040646531: | XM_040646531 | LOC101747452 | LOC101747452 |
| DMC | 12 | 3316417 | 30 | intron1 | 47452:XM_040646531: | XM_040646531 | LOC101747452 | LOC101747452 |
| DMC | 12 | 3316417 | 30 | cds | 358554:XM_015292858 | XM_015292858 | ABHD14A | LOC100858554 |
| DMC | 12 | 3316517 | -70 | promoter | 56877:XM_015301851: | XM_015301851 | LOC107056877 | LOC107056877 |
| DMC | 12 | 3316517 | -70 | promoter | 58836:XM_015292854: | XM_015292854 | ACY1 | LOC100858836 |
| DMC | 12 | 3316517 | -70 | promoter | 58836:XM_046900131: | XM_046900131 | ACY1 | LOC100858836 |
| DMC | 12 | 3316517 | -70 | introns | 47452:XM_040646531: | XM_040646531 | LOC101747452 | LOC101747452 |
| DMC | 12 | 3316517 | -70 | introns | 58554:XM_015292858: | XM_015292858 | ABHD14A | LOC100858554 |
| DMC | 12 | 3316517 | -70 | intron1 | 47452:XM_040646531: | XM_040646531 | LOC101747452 | LOC101747452 |
| DMC | 12 | 3419356 | -308 | promoter | 47452:XM_025154514: | XM_025154514 | LOC101747452 | LOC101747452 |
| DMC | 12 | 3419356 | -308 | promoter | 58703:XM_015293494: | XM_015293494 | LSMEM2 | LOC100858703 |
| DMC | 12 | 3419356 | -308 | promoter | 58703:XM_040646575: | XM_040646575 | LSMEM2 | LOC100858703 |
| DMC | 12 | 3419356 | -308 | promoter | 58703:XM_046900158: | XM_046900158 | LSMEM2 | LOC100858703 |
| DMC | 12 | 3419356 | -308 | promoter | 59903:XM_003641994: | XM_003641994 | HYAL3 | LOC100859903 |
| DMC | 12 | 3419356 | -308 | introns | 47452:XM_040646531: | XM_040646531 | LOC101747452 | LOC101747452 |
| DMC | 12 | 3419356 | -308 | introns | 59903:XM_015293492: | XM_015293492 | HYAL3 | LOC100859903 |
| DMC | 12 | 3419356 | -308 | intron1 | 47452:XM_040646531: | XM_040646531 | LOC101747452 | LOC101747452 |
| DMC | 12 | 3419356 | -308 | intron1 | 59903:XM_015293492: | XM_015293492 | HYAL3 | LOC100859903 |
| DMC | 13 | 9906370 | -1409 | introns | 145:XM_040647273:int | XM_040647273 | UIMC1 | LOC416145 |
| DMC | 13 | 9906370 | -1409 | introns | 145:XM_040647274:int | XM_040647274 | UIMC1 | LOC416145 |
| DMC | 13 | 9906370 | -1409 | introns | 6145:XM_040647277:int | XM_040647277 | UIMC1 | LOC416145 |
| DMC | 13 | 9906370 | -1409 | introns | 145:XR_006931396:intr | XR_006931396 | UIMC1 | LOC416145 |
| DMC | 13 | 9906370 | -1409 | introns | 145:XM_040647275:int | XM_040647275 | UIMC1 | LOC416145 |
| DMC | 13 | 9906370 | -1409 | introns | 145:XM_040647269:int | XM_040647269 | UIMC1 | LOC416145 |
| DMC | 13 | 9906370 | -1409 | introns | 145:XM_040647272:int | XM_040647272 | UIMC1 | LOC416145 |
| DMC | 13 | 9906370 | -1409 | introns | 145:XM_040647276:int | XM_040647276 | UIMC1 | LOC416145 |
| DMC | 13 | 9906370 | -1409 | introns | 145:XR_005839710:intr | XR_005839710 | UIMC1 | LOC416145 |
| DMC | 13 | 9906370 | -1409 | introns | 145:XR_005839715:intr | XR_005839715 | UIMC1 | LOC416145 |
| DMC | 13 | 9906370 | -1409 | introns | 145:XR_005839717:intr | XR_005839717 | UIMC1 | LOC416145 |
| DMC | 13 | 9906370 | -1409 | introns | 145:XR_005839706:intr | XR_005839706 | UIMC1 | LOC416145 |
| DMC | 13 | 9906370 | -1409 | introns | 145:XR_005839707:intr | XR_005839707 | UIMC1 | LOC416145 |
| DMC | 13 | 9906370 | -1409 | introns | 145:XR_005839709:intr | XR_005839709 | UIMC1 | LOC416145 |
| DMC | 13 | 9906370 | -1409 | introns | 145:XR_005839711:intr | XR_005839711 | UIMC1 | LOC416145 |
| DMC | 13 | 9906370 | -1409 | introns | 145:XR_005839712:intr | XR_005839712 | UIMC1 | LOC416145 |
| DMC | 13 | 9906370 | -1409 | introns | 145:XR_005839713:intr | XR_005839713 | UIMC1 | LOC416145 |
| DMC | 13 | 9906370 | -1409 | introns | 145:XR_005839714:intr | XR_005839714 | UIMC1 | LOC416145 |
| DMC | 13 | 9906370 | -1409 | introns | 145:XR_005839716:intr | XR_005839716 | UIMC1 | LOC416145 |
| DMC | 13 | 9906370 | -1409 | introns | 145:XR_005839708:intr | XR_005839708 | UIMC1 | LOC416145 |
| DMC | 13 | 9906370 | -1409 | introns | 145:XM_040647270:int | XM_040647270 | UIMC1 | LOC416145 |
| DMC | 13 | 9906370 | -1409 | introns | 145:XM_040647266:int | XM_040647266 | UIMC1 | LOC416145 |
| DMC | 13 | 9906370 | -1409 | introns | 145:XM_040647263:int | XM_040647263 | UIMC1 | LOC416145 |
| DMC | 13 | 9906370 | -1409 | introns | 145:XM_040647264:int | XM_040647264 | UIMC1 | LOC416145 |
| DMC | 13 | 9906370 | -1409 | introns | 145:XM_040647265:int | XM_040647265 | UIMC1 | LOC416145 |
| DMC | 13 | 9906370 | -1409 | introns | 145:XM_040647271:int | XM_040647271 | UIMC1 | LOC416145 |
| DMC | 13 | 9906370 | -1409 | introns | 145:XM_040647268:int | XM_040647268 | UIMC1 | LOC416145 |
| DMC | 13 | 9906370 | -1409 | introns | 001381:NONGGAT00239 | NONGGAT002396 | NONGGAG001381 | NONGGAG001381 |
| DMC | 13 | 9906370 | -1409 | intron1 | 001381:NONGGAT00239 | NONGGAT002396 | NONGGAG001381 | NONGGAG001381 |
| DMC | 13 | 9959153 | -334 | exons | 5391:XM_015293731:e | XM_015293731 | SNCB | LOC395391 |
| DMC | 13 | 9959153 | -334 | exons | 95391:NM_204671:exo | NM_204671 | SNCB | LOC395391 |
| DMC | 13 | 9959153 | -334 | UTR3 | 91:XM_015293731:3'UT | XM_015293731 | SNCB | LOC395391 |
| DMC | 13 | 9959153 | -334 | UTR3 | 391:NM_204671:3'UTR | NM_204671 | SNCB | LOC395391 |
| DMC | 13 | 10057932 | 684 | promoter | 017192:ENSGALT00010( | NSGALT0001004144 | NSGALG000100171 | NSGALG0001001719 |
| DMC | 13 | 10057932 | 684 | introns | 57281:XM_015293891: | XM_015293891 | CPLX2 | LOC100857281 |
| DMC | 13 | 10057932 | 684 | intron1 | 57281:XM_015293891: | XM_015293891 | CPLX2 | LOC100857281 |
| DMC | 14 | 2527831 | 4755 | other | NA | NA | NA | NA |
| DMC | 14 | 2669066 | -6197 | introns | 5465:XR_005839952:int | XR_005839952 | ELFN1 | LOC416465 |
| DMC | 14 | 2669066 | -6197 | introns | 5465:XM_015294341:in | XM_015294341 | ELFN1 | LOC416465 |
| DMC | 14 | 2669066 | -6197 | introns | 5465:XM_040647847:in | XM_040647847 | ELFN1 | LOC416465 |
| DMC | 14 | 2669066 | -6197 | introns | 5465:XM_046900993:in | XM_046900993 | ELFN1 | LOC416465 |
| DMC | 14 | 2669066 | -6197 | introns | 5465:XM_040647848:in | XM_040647848 | ELFN1 | LOC416465 |

| | | | | | | | | |
|---|---|---|---|---|---|---|---|---|
| DMC | 14 | 2669066 | -6197 | introns | 5465:XM_040647849:in | XM_040647849 | ELFN1 | LOC416465 |
| DMC | 14 | 2669066 | -6197 | intron1 | 5465:XR_005839952:int | XR_005839952 | ELFN1 | LOC416465 |
| DMC | 14 | 2669066 | -6197 | intron1 | 5465:XM_015294341:in | XM_015294341 | ELFN1 | LOC416465 |
| DMC | 14 | 2669066 | -6197 | intron1 | 5465:XM_040647847:in | XM_040647847 | ELFN1 | LOC416465 |
| DMC | 14 | 2669066 | -6197 | intron1 | 5465:XM_046900993:in | XM_046900993 | ELFN1 | LOC416465 |
| DMC | 14 | 2669066 | -6197 | intron1 | 5465:XM_040647848:in | XM_040647848 | ELFN1 | LOC416465 |
| DMC | 14 | 2669066 | -6197 | intron1 | 5465:XM_040647849:in | XM_040647849 | ELFN1 | LOC416465 |
| DMC | 14 | 3067919 | 2344 | introns | 658:XM_004945136:int | XM_004945136 | MAD1L1 | LOC427658 |
| DMC | 14 | 3067919 | 2344 | introns | 658:XM_040647852:int | XM_040647852 | MAD1L1 | LOC427658 |
| DMC | 14 | 3067919 | 2344 | introns | 658:XM_040647851:int | XM_040647851 | MAD1L1 | LOC427658 |
| DMC | 14 | 3067919 | 2344 | introns | 658:XM_040647853:int | XM_040647853 | MAD1L1 | LOC427658 |
| DMC | 14 | 3146650 | 410 | promoter | 04964:INRAGALT00000 | IRAGALT000000105 | RAGALG00000049 | RAGALG0000000496 |
| DMC | 14 | 3960061 | 714 | promoter | 5463:XM_015294358:pr | XM_015294358 | FOXK1 | LOC395463 |
| DMC | 14 | 3960061 | 714 | promoter | 5463:XM_015294360:pr | XM_015294360 | FOXK1 | LOC395463 |
| DMC | 14 | 3960061 | 714 | exons | 008923:TAGAGALT0000 | GAGALT0000001036 | GAGALG000000008 | GAGALG0000000089 |
| DMC | 14 | 3960061 | 714 | exon1 | 008923:TAGAGALT0000 | GAGALT0000001036 | GAGALG000000008 | GAGALG0000000089 |
| DMC | 14 | 12686663 | -150 | promoter | 5269:NM_001396212:pr | NM_001396212 | SRL | LOC396269 |
| DMC | 14 | 12686663 | -150 | promoter | 5269:NM_001396213:pr | NM_001396213 | SRL | LOC396269 |
| DMC | 14 | 12686663 | -150 | promoter | 96269:NM_205329:pro | NM_205329 | SRL | LOC396269 |
| DMC | 14 | 12686663 | -150 | promoter | 5269:XM_046900561:pr | XM_046900561 | SRL | LOC396269 |
| DMC | 14 | 12686663 | -150 | downstream | 33:XM_015294648:dov | XM_015294648 | TFAP4 | LOC430233 |
| DMC | 14 | 12686663 | -150 | downstream | 33:XM_015294647:dov | XM_015294647 | TFAP4 | LOC430233 |
| DMC | 14 | 13299599 | 2351 | promoter | 57632:XM_046900748: | XM_046900748 | LOC100857632 | LOC100857632 |
| DMC | 15 | 7007155 | -615 | downstream | 5088:NM_205175:dow | NM_205175 | CRYBB2 | LOC396088 |
| DMC | 15 | 7007155 | -615 | downstream | 88:XM_015275225:dov | XM_015275225 | CRYBB2 | LOC396088 |
| DMC | 15 | 7112579 | 3857 | other | NA | NA | NA | NA |
| DMC | 15 | 8104445 | -190 | promoter | 9847:XM_040648116:pr | XM_040648116 | GSTT1L | LOC769847 |
| DMC | 15 | 8104445 | -190 | introns | 936:XM_004934333:intr | XM_004934333 | DDX51 | LOC416936 |
| DMC | 15 | 10798413 | -423 | promoter | 17006:XM_415296:pro | XM_415296 | GAL3ST1 | LOC417006 |
| DMC | 15 | 10798413 | -423 | promoter | 7006:XM_025155687:pr | XM_025155687 | GAL3ST1 | LOC417006 |
| DMC | 15 | 10798413 | -423 | promoter | 7006:XM_004934444:pr | XM_004934444 | GAL3ST1 | LOC417006 |
| DMC | 15 | 10798413 | -423 | promoter | 7006:XM_004934445:pr | XM_004934445 | GAL3ST1 | LOC417006 |
| DMC | 15 | 10798413 | -423 | introns | 17006:XM_415296:intr | XM_415296 | GAL3ST1 | LOC417006 |
| DMC | 15 | 10798413 | -423 | intron1 | 17006:XM_415296:intr | XM_415296 | GAL3ST1 | LOC417006 |
| DMC | 16 | 498166 | -97 | promoter | 019875:INRAGALT00000 | IRAGALT000000438 | RAGALG00000019 | RAGALG0000001987 |
| DMC | 16 | 498166 | -97 | promoter | 020519:INRAGALT00000 | IRAGALT000000459 | RAGALG00000020 | RAGALG0000002051 |
| DMC | 16 | 498166 | -97 | promoter | 020519:INRAGALT00000 | IRAGALT000000459 | RAGALG00000020 | RAGALG0000002051 |
| DMC | 16 | 498166 | -97 | downstream | 7023:XR_005840286:do | XR_005840286 | LOC121107023 | LOC121107023 |
| DMC | 16 | 498174 | -105 | promoter | 019875:INRAGALT00000 | IRAGALT000000438 | RAGALG00000019 | RAGALG0000001987 |
| DMC | 16 | 498174 | -105 | promoter | 020519:INRAGALT00000 | IRAGALT000000459 | RAGALG00000020 | RAGALG0000002051 |
| DMC | 16 | 498174 | -105 | promoter | 020519:INRAGALT00000 | IRAGALT000000459 | RAGALG00000020 | RAGALG0000002051 |
| DMC | 16 | 498174 | -105 | downstream | 7023:XR_005840286:do | XR_005840286 | LOC121107023 | LOC121107023 |
| DMC | 16 | 2070908 | -285 | promoter | 54697:XM_040648873: | XM_040648873 | LOC107054697 | LOC107054697 |
| DMC | 16 | 2070908 | -285 | promoter | 54697:XM_040648874: | XM_040648874 | LOC107054697 | LOC107054697 |
| DMC | 16 | 2070908 | -285 | promoter | 54697:XM_046901605: | XM_046901605 | LOC107054697 | LOC107054697 |
| DMC | 16 | 2070908 | -285 | exons | 7095:XM_046901500:e | XM_046901500 | ZNF692 | LOC427095 |
| DMC | 16 | 2070908 | -285 | exons | 7095:XR_006931602:ex | XR_006931602 | ZNF692 | LOC427095 |
| DMC | 16 | 2070908 | -285 | exons | 7095:XR_006931603:ex | XR_006931603 | ZNF692 | LOC427095 |
| DMC | 16 | 2070908 | -285 | exons | 7095:XR_006931601:ex | XR_006931601 | ZNF692 | LOC427095 |
| DMC | 16 | 2070908 | -285 | exons | 7095:XR_006931604:ex | XR_006931604 | ZNF692 | LOC427095 |
| DMC | 16 | 2070908 | -285 | exons | 7095:XR_006931605:ex | XR_006931605 | ZNF692 | LOC427095 |
| DMC | 16 | 2070908 | -285 | exons | 7095:XR_006931606:ex | XR_006931606 | ZNF692 | LOC427095 |
| DMC | 16 | 2070908 | -285 | exons | 7095:XR_006931607:ex | XR_006931607 | ZNF692 | LOC427095 |
| DMC | 16 | 2070908 | -285 | exons | 095:NM_001099356:ex | NM_001099356 | ZNF692 | LOC427095 |
| DMC | 16 | 2070908 | -285 | exons | 095:NM_001396499:ex | NM_001396499 | ZNF692 | LOC427095 |
| DMC | 16 | 2070908 | -285 | exons | 7095:XM_046901498:ex | XM_046901498 | ZNF692 | LOC427095 |
| DMC | 16 | 2070908 | -285 | exons | 7095:XM_046901499:ex | XM_046901499 | ZNF692 | LOC427095 |
| DMC | 16 | 2070908 | -285 | cds | 7095:XM_046901500:e | XM_046901500 | ZNF692 | LOC427095 |
| DMC | 16 | 2070908 | -285 | cds | 095:NM_001099356:ex | NM_001099356 | ZNF692 | LOC427095 |
| DMC | 16 | 2070908 | -285 | cds | 095:NM_001396499:ex | NM_001396499 | ZNF692 | LOC427095 |
| DMC | 16 | 2070908 | -285 | cds | 7095:XM_046901498:ex | XM_046901498 | ZNF692 | LOC427095 |
| DMC | 16 | 2070908 | -285 | cds | 7095:XM_046901499:ex | XM_046901499 | ZNF692 | LOC427095 |
| DMC | 16 | 2178551 | 30 | promoter | 01750:INRAGALT00000 | IRAGALT000001021 | RAGALG00000101 | RAGALG0000010175 |
| DMC | 16 | 2178551 | 30 | promoter | 4776:NM_001397836:pr | NM_001397836 | BLEC1 | LOC404776 |
| DMC | 16 | 2178551 | 30 | promoter | 4776:NM_001397837:pr | NM_001397837 | BLEC1 | LOC404776 |
| DMC | 16 | 2178551 | 30 | promoter | 4776:XM_040648586:pr | XM_040648586 | BLEC1 | LOC404776 |
| DMC | 16 | 2178551 | 30 | promoter | 4776:XM_046901430:pr | XM_046901430 | BLEC1 | LOC404776 |

| | | | | | | | | |
|---|---|---|---|---|---|---|---|---|
| DMC | 16 | 2178551 | 30 | promoter | 4776:XM_046901431:pr | XM_046901431 | BLEC1 | LOC404776 |
| DMC | 16 | 2178551 | 30 | promoter | 4776:XM_046901432:pr | XM_046901432 | BLEC1 | LOC404776 |
| DMC | 16 | 2178551 | 30 | promoter | 4776:XM_046901434:pr | XM_046901434 | BLEC1 | LOC404776 |
| DMC | 16 | 2178551 | 30 | exons | 3259:NM_001044682:e | NM_001044682 | BLEC2 | LOC693259 |
| DMC | 16 | 2178551 | 30 | introns | 4776:XM_046901433:in | XM_046901433 | BLEC1 | LOC404776 |
| DMC | 16 | 2178551 | 30 | intron1 | 4776:XM_046901433:in | XM_046901433 | BLEC1 | LOC404776 |
| DMC | 16 | 2178551 | 30 | UTR3 | 59:NM_001044682:3'U | NM_001044682 | BLEC2 | LOC693259 |
| DMC | 16 | 2189109 | 117 | promoter | 256:NM_001044679:pr | NM_001044679 | BLB1 | LOC693256 |
| DMC | 16 | 2189109 | 117 | promoter | 3256:XM_015294994:pr | XM_015294994 | BLB1 | LOC693256 |
| DMC | 16 | 2189109 | 117 | exons | 7048:NM_001034816:e | NM_001034816 | TAPBP | LOC417048 |
| DMC | 16 | 2189109 | 117 | exons | 7048:NM_001206611:e | NM_001206611 | TAPBP | LOC417048 |
| DMC | 16 | 2189109 | 117 | cds | 7048:NM_001034816:e | NM_001034816 | TAPBP | LOC417048 |
| DMC | 16 | 2189109 | 117 | cds | 7048:NM_001206611:e | NM_001206611 | TAPBP | LOC417048 |
| DMC | 16 | 2196382 | 95 | exons | '049:NM_001398126:ex | NM_001398126 | BRD2 | LOC417049 |
| DMC | 16 | 2196382 | 95 | exons | '049:NM_001398127:ex | NM_001398127 | BRD2 | LOC417049 |
| DMC | 16 | 2196382 | 95 | exons | '049:NM_001398128:ex | NM_001398128 | BRD2 | LOC417049 |
| DMC | 16 | 2196382 | 95 | exons | )49:NM_001398129:ex | NM_001398129 | BRD2 | LOC417049 |
| DMC | 16 | 2196382 | 95 | cds | '049:NM_001398126:ex | NM_001398126 | BRD2 | LOC417049 |
| DMC | 16 | 2196382 | 95 | cds | '049:NM_001398127:ex | NM_001398127 | BRD2 | LOC417049 |
| DMC | 16 | 2196382 | 95 | cds | '049:NM_001398128:ex | NM_001398128 | BRD2 | LOC417049 |
| DMC | 16 | 2196382 | 95 | cds | )49:NM_001398129:ex | NM_001398129 | BRD2 | LOC417049 |
| DMC | 16 | 2222961 | 96 | promoter | '727:NM_001135968:pr | NM_001135968 | TAP1 | LOC427727 |
| DMC | 16 | 2222961 | 96 | promoter | '727:XM_015294989:pr | XM_015294989 | TAP1 | LOC427727 |
| DMC | 16 | 2222961 | 96 | exons | '728:NM_001099357:ex | NM_001099357 | TAP2 | LOC427728 |
| DMC | 16 | 2222961 | 96 | exons | '728:XM_046901501:ex | XM_046901501 | TAP2 | LOC427728 |
| DMC | 16 | 2222961 | 96 | cds | 7728:NM_001099357:e | NM_001099357 | TAP2 | LOC427728 |
| DMC | 16 | 2222961 | 96 | cds | 7728:XM_046901501:e | XM_046901501 | TAP2 | LOC427728 |
| DMC | 17 | 5614668 | -95 | promoter | )08246:NONGGAT0135 | NONGGAT013567 | NONGGAG008246 | NONGGAG008246 |
| DMC | 17 | 5614668 | -95 | promoter | '768:XM_003642278:pr | XM_003642278 | CERCAM | LOC427768 |
| DMC | 17 | 5614668 | -95 | introns | 768:XM_015279709:int | XM_015279709 | CERCAM | LOC427768 |
| DMC | 17 | 5614668 | -95 | introns | 768:XM_015279708:int | XM_015279708 | CERCAM | LOC427768 |
| DMC | 17 | 5614668 | -95 | introns | 768:XM_046901886:int | XM_046901886 | CERCAM | LOC427768 |
| DMC | 17 | 7073889 | 1670 | introns | 171:XM_015279756:int | XM_015279756 | NTNG2 | LOC417171 |
| DMC | 17 | 7073889 | 1670 | introns | '171:XM_015279758:in | XM_015279758 | NTNG2 | LOC417171 |
| DMC | 17 | 10083304 | -856 | promoter | !325:XM_001235477:pr | XM_001235477 | NR6A1 | LOC772325 |
| DMC | 17 | 10083304 | -856 | promoter | !325:XM_040649148:pr | XM_040649148 | NR6A1 | LOC772325 |
| DMC | 17 | 10083304 | -856 | introns | !325:XM_015279582:in | XM_015279582 | NR6A1 | LOC772325 |
| DMC | 17 | 10083304 | -856 | introns | !325:XM_015279583:in | XM_015279583 | NR6A1 | LOC772325 |
| DMC | 17 | 10083304 | -856 | introns | !325:XM_015279584:in | XM_015279584 | NR6A1 | LOC772325 |
| DMC | 17 | 10083304 | -856 | intron1 | !325:XM_015279584:in | XM_015279584 | NR6A1 | LOC772325 |
| DMC | 17 | 10083318 | -870 | promoter | !325:XM_001235477:pr | XM_001235477 | NR6A1 | LOC772325 |
| DMC | 17 | 10083318 | -870 | promoter | !325:XM_040649148:pr | XM_040649148 | NR6A1 | LOC772325 |
| DMC | 17 | 10083318 | -870 | introns | !325:XM_015279582:in | XM_015279582 | NR6A1 | LOC772325 |
| DMC | 17 | 10083318 | -870 | introns | !325:XM_015279583:in | XM_015279583 | NR6A1 | LOC772325 |
| DMC | 17 | 10083318 | -870 | introns | !325:XM_015279584:in | XM_015279584 | NR6A1 | LOC772325 |
| DMC | 17 | 10083318 | -870 | intron1 | !325:XM_015279584:in | XM_015279584 | NR6A1 | LOC772325 |
| DMC | 17 | 10083453 | -1005 | promoter | !325:XM_001235477:pr | XM_001235477 | NR6A1 | LOC772325 |
| DMC | 17 | 10083453 | -1005 | promoter | !325:XM_040649148:pr | XM_040649148 | NR6A1 | LOC772325 |
| DMC | 17 | 10083453 | -1005 | introns | !325:XM_015279582:in | XM_015279582 | NR6A1 | LOC772325 |
| DMC | 17 | 10083453 | -1005 | introns | !325:XM_015279583:in | XM_015279583 | NR6A1 | LOC772325 |
| DMC | 17 | 10083453 | -1005 | introns | !325:XM_015279584:in | XM_015279584 | NR6A1 | LOC772325 |
| DMC | 17 | 10083453 | -1005 | intron1 | !325:XM_015279584:in | XM_015279584 | NR6A1 | LOC772325 |
| DMC | 17 | 10247319 | 1794 | downstream | 5682:INRAGALT000000 | RAGALT000000121 | RAGALG00000056 | RAGALG00000056 |
| DMC | 17 | 10402661 | 10230 | introns | '093:NM_001397668:in | NM_001397668 | PBX3 | LOC417093 |
| DMC | 17 | 10402661 | 10230 | introns | '093:NM_001397670:in | NM_001397670 | PBX3 | LOC417093 |
| DMC | 17 | 10402661 | 10230 | introns | '093:NM_001397673:in | NM_001397673 | PBX3 | LOC417093 |
| DMC | 17 | 10402661 | 10230 | introns | '093:XM_040649039:in | XM_040649039 | PBX3 | LOC417093 |
| DMC | 17 | 10402661 | 10230 | introns | '093:NM_001397671:in | NM_001397671 | PBX3 | LOC417093 |
| DMC | 17 | 10402661 | 10230 | introns | '093:NM_001397674:in | NM_001397674 | PBX3 | LOC417093 |
| DMC | 17 | 10402661 | 10230 | introns | '093:NM_001030678:in | NM_001030678 | PBX3 | LOC417093 |
| DMC | 17 | 10402661 | 10230 | introns | '093:NM_001397672:in | NM_001397672 | PBX3 | LOC417093 |
| DMC | 17 | 10524429 | 4289 | other | NA | NA | NA | NA |
| DMC | 17 | 11003840 | -110 | promoter | )05696:INRAGALT00000 | RAGALT00000012 | RAGALG00000056 | RAGALG00000056 |
| DMC | 17 | 11003840 | -110 | promoter | )05697:INRAGALT00000 | RAGALT00000012 | RAGALG00000056 | RAGALG00000056 |
| DMC | 17 | 11003840 | -110 | promoter | )05694:INRAGALT00000 | RAGALT00000012 | RAGALG00000056 | RAGALG00000056 |
| DMC | 17 | 11003840 | -110 | exons | )05696:INRAGALT0000 | RAGALT00000012 | RAGALG00000056 | RAGALG00000056 |
| DMC | 17 | 11003840 | -110 | exon1 | )05696:INRAGALT0000 | RAGALT00000012 | RAGALG00000056 | RAGALG00000056 |

| | | | | | | | | |
|---|---|---|---|---|---|---|---|---|
| DMC | 17 | 11003840 | -110 | downstream | 5695:INRAGALT000000 | RAGALT000000122( | RAGALG000000056 | RAGALG0000000569 |
| DMC | 17 | 11003847 | -117 | promoter | 05696:INRAGALT00000 | RAGALT000000122( | RAGALG000000056 | RAGALG0000000569 |
| DMC | 17 | 11003847 | -117 | promoter | 05697:INRAGALT00000 | RAGALT000000122( | RAGALG000000056 | RAGALG0000000569 |
| DMC | 17 | 11003847 | -117 | promoter | 05694:INRAGALT00000 | RAGALT000000121 | RAGALG000000056 | RAGALG0000000569 |
| DMC | 17 | 11003847 | -117 | exons | 005696:INRAGALT0000 | RAGALT000000122( | RAGALG000000056 | RAGALG0000000569 |
| DMC | 17 | 11003847 | -117 | exon1 | 005696:INRAGALT0000 | RAGALT000000122( | RAGALG000000056 | RAGALG0000000569 |
| DMC | 17 | 11003847 | -117 | downstream | 5695:INRAGALT000000 | RAGALT000000122( | RAGALG000000056 | RAGALG0000000569 |
| DMC | 17 | 11003858 | -128 | promoter | 05696:INRAGALT00000 | RAGALT000000122( | RAGALG000000056 | RAGALG0000000569 |
| DMC | 17 | 11003858 | -128 | promoter | 05697:INRAGALT00000 | RAGALT000000122( | RAGALG000000056 | RAGALG0000000569 |
| DMC | 17 | 11003858 | -128 | promoter | 05694:INRAGALT00000 | RAGALT000000121 | RAGALG000000056 | RAGALG0000000569 |
| DMC | 17 | 11003858 | -128 | exons | 005696:INRAGALT0000 | RAGALT000000122( | RAGALG000000056 | RAGALG0000000569 |
| DMC | 17 | 11003858 | -128 | exon1 | 005696:INRAGALT0000 | RAGALT000000122( | RAGALG000000056 | RAGALG0000000569 |
| DMC | 17 | 11003858 | -128 | downstream | 5695:INRAGALT000000 | RAGALT000000122( | RAGALG000000056 | RAGALG0000000569 |
| DMC | 17 | 11006346 | 467 | promoter | '087:XM_040648984:pr | XM_040648984 | GARNL3 | LOC417087 |
| DMC | 17 | 11006346 | 467 | promoter | '087:XM_040648985:pr | XM_040648985 | GARNL3 | LOC417087 |
| DMC | 17 | 11006346 | 467 | promoter | '087:XM_040648989:pr | XM_040648989 | GARNL3 | LOC417087 |
| DMC | 17 | 11006346 | 467 | promoter | '087:XM_040648990:pr | XM_040648990 | GARNL3 | LOC417087 |
| DMC | 17 | 11006346 | 467 | promoter | '087:XM_040648991:pr | XM_040648991 | GARNL3 | LOC417087 |
| DMC | 17 | 11006346 | 467 | promoter | '087:XM_040648992:pr | XM_040648992 | GARNL3 | LOC417087 |
| DMC | 17 | 11006346 | 467 | promoter | '087:XM_040648995:pr | XM_040648995 | GARNL3 | LOC417087 |
| DMC | 17 | 11006346 | 467 | promoter | '087:XM_040648996:pr | XM_040648996 | GARNL3 | LOC417087 |
| DMC | 17 | 11006346 | 467 | promoter | '087:XM_040648997:pr | XM_040648997 | GARNL3 | LOC417087 |
| DMC | 17 | 11006346 | 467 | promoter | '087:XM_040648998:pr | XM_040648998 | GARNL3 | LOC417087 |
| DMC | 17 | 11006346 | 467 | promoter | '087:XM_040648999:pr | XM_040648999 | GARNL3 | LOC417087 |
| DMC | 17 | 11006346 | 467 | promoter | '087:XM_040649004:pr | XM_040649004 | GARNL3 | LOC417087 |
| DMC | 17 | 11006346 | 467 | promoter | '087:XM_040649014:pr | XM_040649014 | GARNL3 | LOC417087 |
| DMC | 17 | 11006346 | 467 | promoter | '087:XM_040649016:pr | XM_040649016 | GARNL3 | LOC417087 |
| DMC | 17 | 11006346 | 467 | promoter | '087:XM_040649021:pr | XM_040649021 | GARNL3 | LOC417087 |
| DMC | 17 | 11006346 | 467 | promoter | '087:XM_040649022:pr | XM_040649022 | GARNL3 | LOC417087 |
| DMC | 17 | 11006346 | 467 | promoter | '087:XM_040649023:pr | XM_040649023 | GARNL3 | LOC417087 |
| DMC | 17 | 11006346 | 467 | promoter | '087:XM_040649024:pr | XM_040649024 | GARNL3 | LOC417087 |
| DMC | 17 | 11006346 | 467 | promoter | '087:XM_046901680:pr | XM_046901680 | GARNL3 | LOC417087 |
| DMC | 17 | 11006346 | 467 | exons | 005697:INRAGALT0000 | RAGALT000000122( | RAGALG000000056 | RAGALG0000000569 |
| DMC | 17 | 11006346 | 467 | downstream | 9809:TAGAGALT00000( | GAGALT0000001351 | GAGALG000000059 | GAGALG0000000598 |
| DMC | 17 | 11006355 | 458 | promoter | '087:XM_040648984:pr | XM_040648984 | GARNL3 | LOC417087 |
| DMC | 17 | 11006355 | 458 | promoter | '087:XM_040648985:pr | XM_040648985 | GARNL3 | LOC417087 |
| DMC | 17 | 11006355 | 458 | promoter | '087:XM_040648989:pr | XM_040648989 | GARNL3 | LOC417087 |
| DMC | 17 | 11006355 | 458 | promoter | '087:XM_040648990:pr | XM_040648990 | GARNL3 | LOC417087 |
| DMC | 17 | 11006355 | 458 | promoter | '087:XM_040648991:pr | XM_040648991 | GARNL3 | LOC417087 |
| DMC | 17 | 11006355 | 458 | promoter | '087:XM_040648992:pr | XM_040648992 | GARNL3 | LOC417087 |
| DMC | 17 | 11006355 | 458 | promoter | '087:XM_040648995:pr | XM_040648995 | GARNL3 | LOC417087 |
| DMC | 17 | 11006355 | 458 | promoter | '087:XM_040648996:pr | XM_040648996 | GARNL3 | LOC417087 |
| DMC | 17 | 11006355 | 458 | promoter | '087:XM_040648997:pr | XM_040648997 | GARNL3 | LOC417087 |
| DMC | 17 | 11006355 | 458 | promoter | '087:XM_040648998:pr | XM_040648998 | GARNL3 | LOC417087 |
| DMC | 17 | 11006355 | 458 | promoter | '087:XM_040648999:pr | XM_040648999 | GARNL3 | LOC417087 |
| DMC | 17 | 11006355 | 458 | promoter | '087:XM_040649004:pr | XM_040649004 | GARNL3 | LOC417087 |
| DMC | 17 | 11006355 | 458 | promoter | '087:XM_040649014:pr | XM_040649014 | GARNL3 | LOC417087 |
| DMC | 17 | 11006355 | 458 | promoter | '087:XM_040649016:pr | XM_040649016 | GARNL3 | LOC417087 |
| DMC | 17 | 11006355 | 458 | promoter | '087:XM_040649021:pr | XM_040649021 | GARNL3 | LOC417087 |
| DMC | 17 | 11006355 | 458 | promoter | '087:XM_040649022:pr | XM_040649022 | GARNL3 | LOC417087 |
| DMC | 17 | 11006355 | 458 | promoter | '087:XM_040649023:pr | XM_040649023 | GARNL3 | LOC417087 |
| DMC | 17 | 11006355 | 458 | promoter | '087:XM_040649024:pr | XM_040649024 | GARNL3 | LOC417087 |
| DMC | 17 | 11006355 | 458 | promoter | '087:XM_046901680:pr | XM_046901680 | GARNL3 | LOC417087 |
| DMC | 17 | 11006355 | 458 | exons | 005697:INRAGALT0000 | RAGALT000000122( | RAGALG000000056 | RAGALG0000000569 |
| DMC | 17 | 11006355 | 458 | downstream | 9809:TAGAGALT00000( | GAGALT0000001351 | GAGALG000000059 | GAGALG0000000598 |
| DMC | 17 | 11006363 | 450 | promoter | '087:XM_040648984:pr | XM_040648984 | GARNL3 | LOC417087 |
| DMC | 17 | 11006363 | 450 | promoter | '087:XM_040648985:pr | XM_040648985 | GARNL3 | LOC417087 |
| DMC | 17 | 11006363 | 450 | promoter | '087:XM_040648989:pr | XM_040648989 | GARNL3 | LOC417087 |
| DMC | 17 | 11006363 | 450 | promoter | '087:XM_040648990:pr | XM_040648990 | GARNL3 | LOC417087 |
| DMC | 17 | 11006363 | 450 | promoter | '087:XM_040648991:pr | XM_040648991 | GARNL3 | LOC417087 |
| DMC | 17 | 11006363 | 450 | promoter | '087:XM_040648992:pr | XM_040648992 | GARNL3 | LOC417087 |
| DMC | 17 | 11006363 | 450 | promoter | '087:XM_040648995:pr | XM_040648995 | GARNL3 | LOC417087 |
| DMC | 17 | 11006363 | 450 | promoter | '087:XM_040648996:pr | XM_040648996 | GARNL3 | LOC417087 |
| DMC | 17 | 11006363 | 450 | promoter | '087:XM_040648997:pr | XM_040648997 | GARNL3 | LOC417087 |
| DMC | 17 | 11006363 | 450 | promoter | '087:XM_040648998:pr | XM_040648998 | GARNL3 | LOC417087 |
| DMC | 17 | 11006363 | 450 | promoter | '087:XM_040648999:pr | XM_040648999 | GARNL3 | LOC417087 |
| DMC | 17 | 11006363 | 450 | promoter | '087:XM_040649004:pr | XM_040649004 | GARNL3 | LOC417087 |

| | | | | | | | | |
|---|---|---|---|---|---|---|---|---|
| DMC | 17 | 11006363 | 450 | promoter | '087:XM_040649014:pr | XM_040649014 | GARNL3 | LOC417087 |
| DMC | 17 | 11006363 | 450 | promoter | '087:XM_040649016:pr | XM_040649016 | GARNL3 | LOC417087 |
| DMC | 17 | 11006363 | 450 | promoter | '087:XM_040649021:pr | XM_040649021 | GARNL3 | LOC417087 |
| DMC | 17 | 11006363 | 450 | promoter | '087:XM_040649022:pr | XM_040649022 | GARNL3 | LOC417087 |
| DMC | 17 | 11006363 | 450 | promoter | '087:XM_040649023:pr | XM_040649023 | GARNL3 | LOC417087 |
| DMC | 17 | 11006363 | 450 | promoter | '087:XM_040649024:pr | XM_040649024 | GARNL3 | LOC417087 |
| DMC | 17 | 11006363 | 450 | promoter | '087:XM_046901680:pr | XM_046901680 | GARNL3 | LOC417087 |
| DMC | 17 | 11006363 | 450 | exons | 005697:INRAGALT0000 | RAGALT000000122 | RAGALG000000056 | RAGALG0000000569 |
| DMC | 17 | 11006363 | 450 | downstream | 9809:TAGAGALT00000 | GAGALT0000001351 | GAGALG000000059 | GAGALG0000000598 |
| DMC | 17 | 11006375 | 438 | promoter | '087:XM_040648984:pr | XM_040648984 | GARNL3 | LOC417087 |
| DMC | 17 | 11006375 | 438 | promoter | '087:XM_040648985:pr | XM_040648985 | GARNL3 | LOC417087 |
| DMC | 17 | 11006375 | 438 | promoter | '087:XM_040648989:pr | XM_040648989 | GARNL3 | LOC417087 |
| DMC | 17 | 11006375 | 438 | promoter | '087:XM_040648990:pr | XM_040648990 | GARNL3 | LOC417087 |
| DMC | 17 | 11006375 | 438 | promoter | '087:XM_040648991:pr | XM_040648991 | GARNL3 | LOC417087 |
| DMC | 17 | 11006375 | 438 | promoter | '087:XM_040648992:pr | XM_040648992 | GARNL3 | LOC417087 |
| DMC | 17 | 11006375 | 438 | promoter | '087:XM_040648995:pr | XM_040648995 | GARNL3 | LOC417087 |
| DMC | 17 | 11006375 | 438 | promoter | '087:XM_040648996:pr | XM_040648996 | GARNL3 | LOC417087 |
| DMC | 17 | 11006375 | 438 | promoter | '087:XM_040648997:pr | XM_040648997 | GARNL3 | LOC417087 |
| DMC | 17 | 11006375 | 438 | promoter | '087:XM_040648998:pr | XM_040648998 | GARNL3 | LOC417087 |
| DMC | 17 | 11006375 | 438 | promoter | '087:XM_040648999:pr | XM_040648999 | GARNL3 | LOC417087 |
| DMC | 17 | 11006375 | 438 | promoter | '087:XM_040649004:pr | XM_040649004 | GARNL3 | LOC417087 |
| DMC | 17 | 11006375 | 438 | promoter | '087:XM_040649014:pr | XM_040649014 | GARNL3 | LOC417087 |
| DMC | 17 | 11006375 | 438 | promoter | '087:XM_040649016:pr | XM_040649016 | GARNL3 | LOC417087 |
| DMC | 17 | 11006375 | 438 | promoter | '087:XM_040649021:pr | XM_040649021 | GARNL3 | LOC417087 |
| DMC | 17 | 11006375 | 438 | promoter | '087:XM_040649022:pr | XM_040649022 | GARNL3 | LOC417087 |
| DMC | 17 | 11006375 | 438 | promoter | '087:XM_040649023:pr | XM_040649023 | GARNL3 | LOC417087 |
| DMC | 17 | 11006375 | 438 | promoter | '087:XM_040649024:pr | XM_040649024 | GARNL3 | LOC417087 |
| DMC | 17 | 11006375 | 438 | promoter | '087:XM_046901680:pr | XM_046901680 | GARNL3 | LOC417087 |
| DMC | 17 | 11006375 | 438 | exons | 005697:INRAGALT0000 | RAGALT000000122 | RAGALG000000056 | RAGALG0000000569 |
| DMC | 17 | 11006375 | 438 | downstream | 9809:TAGAGALT00000 | GAGALT0000001351 | GAGALG000000059 | GAGALG0000000598 |
| DMC | 17 | 11006377 | 436 | promoter | '087:XM_040648984:pr | XM_040648984 | GARNL3 | LOC417087 |
| DMC | 17 | 11006377 | 436 | promoter | '087:XM_040648985:pr | XM_040648985 | GARNL3 | LOC417087 |
| DMC | 17 | 11006377 | 436 | promoter | '087:XM_040648989:pr | XM_040648989 | GARNL3 | LOC417087 |
| DMC | 17 | 11006377 | 436 | promoter | '087:XM_040648990:pr | XM_040648990 | GARNL3 | LOC417087 |
| DMC | 17 | 11006377 | 436 | promoter | '087:XM_040648991:pr | XM_040648991 | GARNL3 | LOC417087 |
| DMC | 17 | 11006377 | 436 | promoter | '087:XM_040648992:pr | XM_040648992 | GARNL3 | LOC417087 |
| DMC | 17 | 11006377 | 436 | promoter | '087:XM_040648995:pr | XM_040648995 | GARNL3 | LOC417087 |
| DMC | 17 | 11006377 | 436 | promoter | '087:XM_040648996:pr | XM_040648996 | GARNL3 | LOC417087 |
| DMC | 17 | 11006377 | 436 | promoter | '087:XM_040648997:pr | XM_040648997 | GARNL3 | LOC417087 |
| DMC | 17 | 11006377 | 436 | promoter | '087:XM_040648998:pr | XM_040648998 | GARNL3 | LOC417087 |
| DMC | 17 | 11006377 | 436 | promoter | '087:XM_040648999:pr | XM_040648999 | GARNL3 | LOC417087 |
| DMC | 17 | 11006377 | 436 | promoter | '087:XM_040649004:pr | XM_040649004 | GARNL3 | LOC417087 |
| DMC | 17 | 11006377 | 436 | promoter | '087:XM_040649014:pr | XM_040649014 | GARNL3 | LOC417087 |
| DMC | 17 | 11006377 | 436 | promoter | '087:XM_040649016:pr | XM_040649016 | GARNL3 | LOC417087 |
| DMC | 17 | 11006377 | 436 | promoter | '087:XM_040649021:pr | XM_040649021 | GARNL3 | LOC417087 |
| DMC | 17 | 11006377 | 436 | promoter | '087:XM_040649022:pr | XM_040649022 | GARNL3 | LOC417087 |
| DMC | 17 | 11006377 | 436 | promoter | '087:XM_040649023:pr | XM_040649023 | GARNL3 | LOC417087 |
| DMC | 17 | 11006377 | 436 | promoter | '087:XM_040649024:pr | XM_040649024 | GARNL3 | LOC417087 |
| DMC | 17 | 11006377 | 436 | promoter | '087:XM_046901680:pr | XM_046901680 | GARNL3 | LOC417087 |
| DMC | 17 | 11006377 | 436 | exons | 005697:INRAGALT0000 | RAGALT000000122 | RAGALG000000056 | RAGALG0000000569 |
| DMC | 17 | 11006377 | 436 | downstream | 9809:TAGAGALT00000 | GAGALT0000001351 | GAGALG000000059 | GAGALG0000000598 |
| DMC | 17 | 11006379 | 434 | promoter | '087:XM_040648984:pr | XM_040648984 | GARNL3 | LOC417087 |
| DMC | 17 | 11006379 | 434 | promoter | '087:XM_040648985:pr | XM_040648985 | GARNL3 | LOC417087 |
| DMC | 17 | 11006379 | 434 | promoter | '087:XM_040648989:pr | XM_040648989 | GARNL3 | LOC417087 |
| DMC | 17 | 11006379 | 434 | promoter | '087:XM_040648990:pr | XM_040648990 | GARNL3 | LOC417087 |
| DMC | 17 | 11006379 | 434 | promoter | '087:XM_040648991:pr | XM_040648991 | GARNL3 | LOC417087 |
| DMC | 17 | 11006379 | 434 | promoter | '087:XM_040648992:pr | XM_040648992 | GARNL3 | LOC417087 |
| DMC | 17 | 11006379 | 434 | promoter | '087:XM_040648995:pr | XM_040648995 | GARNL3 | LOC417087 |
| DMC | 17 | 11006379 | 434 | promoter | '087:XM_040648996:pr | XM_040648996 | GARNL3 | LOC417087 |
| DMC | 17 | 11006379 | 434 | promoter | '087:XM_040648997:pr | XM_040648997 | GARNL3 | LOC417087 |
| DMC | 17 | 11006379 | 434 | promoter | '087:XM_040648998:pr | XM_040648998 | GARNL3 | LOC417087 |
| DMC | 17 | 11006379 | 434 | promoter | '087:XM_040648999:pr | XM_040648999 | GARNL3 | LOC417087 |
| DMC | 17 | 11006379 | 434 | promoter | '087:XM_040649004:pr | XM_040649004 | GARNL3 | LOC417087 |
| DMC | 17 | 11006379 | 434 | promoter | '087:XM_040649014:pr | XM_040649014 | GARNL3 | LOC417087 |
| DMC | 17 | 11006379 | 434 | promoter | '087:XM_040649016:pr | XM_040649016 | GARNL3 | LOC417087 |
| DMC | 17 | 11006379 | 434 | promoter | '087:XM_040649021:pr | XM_040649021 | GARNL3 | LOC417087 |
| DMC | 17 | 11006379 | 434 | promoter | '087:XM_040649022:pr | XM_040649022 | GARNL3 | LOC417087 |

| | | | | | | | | |
|---|---|---|---|---|---|---|---|---|
| DMC | 17 | 11006379 | 434 | promoter | '087:XM_040649023:pr | XM_040649023 | GARNL3 | LOC417087 |
| DMC | 17 | 11006379 | 434 | promoter | '087:XM_040649024:pr | XM_040649024 | GARNL3 | LOC417087 |
| DMC | 17 | 11006379 | 434 | promoter | '087:XM_046901680:pr | XM_046901680 | GARNL3 | LOC417087 |
| DMC | 17 | 11006379 | 434 | exons | )05697:INRAGALT0000 | RAGALT000000122 | RAGALG000000056 | RAGALG0000000569 |
| DMC | 17 | 11006379 | 434 | downstream | 9809:TAGAGALT00000 | GAGALT0000001351 | GAGALG000000059 | GAGALG0000000598 |
| DMC | 17 | 11006579 | 234 | promoter | '087:XM_040648984:pr | XM_040648984 | GARNL3 | LOC417087 |
| DMC | 17 | 11006579 | 234 | promoter | '087:XM_040648985:pr | XM_040648985 | GARNL3 | LOC417087 |
| DMC | 17 | 11006579 | 234 | promoter | '087:XM_040648989:pr | XM_040648989 | GARNL3 | LOC417087 |
| DMC | 17 | 11006579 | 234 | promoter | '087:XM_040648990:pr | XM_040648990 | GARNL3 | LOC417087 |
| DMC | 17 | 11006579 | 234 | promoter | '087:XM_040648991:pr | XM_040648991 | GARNL3 | LOC417087 |
| DMC | 17 | 11006579 | 234 | promoter | '087:XM_040648992:pr | XM_040648992 | GARNL3 | LOC417087 |
| DMC | 17 | 11006579 | 234 | promoter | '087:XM_040648995:pr | XM_040648995 | GARNL3 | LOC417087 |
| DMC | 17 | 11006579 | 234 | promoter | '087:XM_040648996:pr | XM_040648996 | GARNL3 | LOC417087 |
| DMC | 17 | 11006579 | 234 | promoter | '087:XM_040648997:pr | XM_040648997 | GARNL3 | LOC417087 |
| DMC | 17 | 11006579 | 234 | promoter | '087:XM_040648998:pr | XM_040648998 | GARNL3 | LOC417087 |
| DMC | 17 | 11006579 | 234 | promoter | '087:XM_040648999:pr | XM_040648999 | GARNL3 | LOC417087 |
| DMC | 17 | 11006579 | 234 | promoter | '087:XM_040649004:pr | XM_040649004 | GARNL3 | LOC417087 |
| DMC | 17 | 11006579 | 234 | promoter | '087:XM_040649014:pr | XM_040649014 | GARNL3 | LOC417087 |
| DMC | 17 | 11006579 | 234 | promoter | '087:XM_040649016:pr | XM_040649016 | GARNL3 | LOC417087 |
| DMC | 17 | 11006579 | 234 | promoter | '087:XM_040649021:pr | XM_040649021 | GARNL3 | LOC417087 |
| DMC | 17 | 11006579 | 234 | promoter | '087:XM_040649022:pr | XM_040649022 | GARNL3 | LOC417087 |
| DMC | 17 | 11006579 | 234 | promoter | '087:XM_040649023:pr | XM_040649023 | GARNL3 | LOC417087 |
| DMC | 17 | 11006579 | 234 | promoter | '087:XM_040649024:pr | XM_040649024 | GARNL3 | LOC417087 |
| DMC | 17 | 11006579 | 234 | promoter | '087:XM_046901680:pr | XM_046901680 | GARNL3 | LOC417087 |
| DMC | 17 | 11006579 | 234 | downstream | 5697:INRAGALT000000 | RAGALT000000122 | RAGALG000000056 | RAGALG0000000569 |
| DMC | 17 | 11006579 | 234 | downstream | 9809:TAGAGALT00000 | GAGALT0000001351 | GAGALG000000059 | GAGALG0000000598 |
| DMC | 17 | 11006587 | 226 | promoter | '087:XM_040648984:pr | XM_040648984 | GARNL3 | LOC417087 |
| DMC | 17 | 11006587 | 226 | promoter | '087:XM_040648985:pr | XM_040648985 | GARNL3 | LOC417087 |
| DMC | 17 | 11006587 | 226 | promoter | '087:XM_040648989:pr | XM_040648989 | GARNL3 | LOC417087 |
| DMC | 17 | 11006587 | 226 | promoter | '087:XM_040648990:pr | XM_040648990 | GARNL3 | LOC417087 |
| DMC | 17 | 11006587 | 226 | promoter | '087:XM_040648991:pr | XM_040648991 | GARNL3 | LOC417087 |
| DMC | 17 | 11006587 | 226 | promoter | '087:XM_040648992:pr | XM_040648992 | GARNL3 | LOC417087 |
| DMC | 17 | 11006587 | 226 | promoter | '087:XM_040648995:pr | XM_040648995 | GARNL3 | LOC417087 |
| DMC | 17 | 11006587 | 226 | promoter | '087:XM_040648996:pr | XM_040648996 | GARNL3 | LOC417087 |
| DMC | 17 | 11006587 | 226 | promoter | '087:XM_040648997:pr | XM_040648997 | GARNL3 | LOC417087 |
| DMC | 17 | 11006587 | 226 | promoter | '087:XM_040648998:pr | XM_040648998 | GARNL3 | LOC417087 |
| DMC | 17 | 11006587 | 226 | promoter | '087:XM_040648999:pr | XM_040648999 | GARNL3 | LOC417087 |
| DMC | 17 | 11006587 | 226 | promoter | '087:XM_040649004:pr | XM_040649004 | GARNL3 | LOC417087 |
| DMC | 17 | 11006587 | 226 | promoter | '087:XM_040649014:pr | XM_040649014 | GARNL3 | LOC417087 |
| DMC | 17 | 11006587 | 226 | promoter | '087:XM_040649016:pr | XM_040649016 | GARNL3 | LOC417087 |
| DMC | 17 | 11006587 | 226 | promoter | '087:XM_040649021:pr | XM_040649021 | GARNL3 | LOC417087 |
| DMC | 17 | 11006587 | 226 | promoter | '087:XM_040649022:pr | XM_040649022 | GARNL3 | LOC417087 |
| DMC | 17 | 11006587 | 226 | promoter | '087:XM_040649023:pr | XM_040649023 | GARNL3 | LOC417087 |
| DMC | 17 | 11006587 | 226 | promoter | '087:XM_040649024:pr | XM_040649024 | GARNL3 | LOC417087 |
| DMC | 17 | 11006587 | 226 | promoter | '087:XM_046901680:pr | XM_046901680 | GARNL3 | LOC417087 |
| DMC | 17 | 11006587 | 226 | downstream | 5697:INRAGALT000000 | RAGALT000000122 | RAGALG000000056 | RAGALG0000000569 |
| DMC | 17 | 11006587 | 226 | downstream | 9809:TAGAGALT00000 | GAGALT0000001351 | GAGALG000000059 | GAGALG0000000598 |
| DMC | 17 | 11006663 | 150 | promoter | '087:XM_040648984:pr | XM_040648984 | GARNL3 | LOC417087 |
| DMC | 17 | 11006663 | 150 | promoter | '087:XM_040648985:pr | XM_040648985 | GARNL3 | LOC417087 |
| DMC | 17 | 11006663 | 150 | promoter | '087:XM_040648989:pr | XM_040648989 | GARNL3 | LOC417087 |
| DMC | 17 | 11006663 | 150 | promoter | '087:XM_040648990:pr | XM_040648990 | GARNL3 | LOC417087 |
| DMC | 17 | 11006663 | 150 | promoter | '087:XM_040648991:pr | XM_040648991 | GARNL3 | LOC417087 |
| DMC | 17 | 11006663 | 150 | promoter | '087:XM_040648992:pr | XM_040648992 | GARNL3 | LOC417087 |
| DMC | 17 | 11006663 | 150 | promoter | '087:XM_040648995:pr | XM_040648995 | GARNL3 | LOC417087 |
| DMC | 17 | 11006663 | 150 | promoter | '087:XM_040648996:pr | XM_040648996 | GARNL3 | LOC417087 |
| DMC | 17 | 11006663 | 150 | promoter | '087:XM_040648997:pr | XM_040648997 | GARNL3 | LOC417087 |
| DMC | 17 | 11006663 | 150 | promoter | '087:XM_040648998:pr | XM_040648998 | GARNL3 | LOC417087 |
| DMC | 17 | 11006663 | 150 | promoter | '087:XM_040648999:pr | XM_040648999 | GARNL3 | LOC417087 |
| DMC | 17 | 11006663 | 150 | promoter | '087:XM_040649004:pr | XM_040649004 | GARNL3 | LOC417087 |
| DMC | 17 | 11006663 | 150 | promoter | '087:XM_040649014:pr | XM_040649014 | GARNL3 | LOC417087 |
| DMC | 17 | 11006663 | 150 | promoter | '087:XM_040649016:pr | XM_040649016 | GARNL3 | LOC417087 |
| DMC | 17 | 11006663 | 150 | promoter | '087:XM_040649021:pr | XM_040649021 | GARNL3 | LOC417087 |
| DMC | 17 | 11006663 | 150 | promoter | '087:XM_040649022:pr | XM_040649022 | GARNL3 | LOC417087 |
| DMC | 17 | 11006663 | 150 | promoter | '087:XM_040649023:pr | XM_040649023 | GARNL3 | LOC417087 |
| DMC | 17 | 11006663 | 150 | promoter | '087:XM_040649024:pr | XM_040649024 | GARNL3 | LOC417087 |
| DMC | 17 | 11006663 | 150 | promoter | '087:XM_046901680:pr | XM_046901680 | GARNL3 | LOC417087 |
| DMC | 17 | 11006663 | 150 | downstream | 5697:INRAGALT000000 | RAGALT000000122 | RAGALG000000056 | RAGALG0000000569 |

| | | | | | | | | |
|---|---|---|---|---|---|---|---|---|
| DMC | 17 | 11006663 | 150 | downstream | 9809:TAGAGALT00000( | GAGALT0000001351 | GAGALG000000059 | GAGALG0000000598 |
| DMC | 17 | 11007025 | 191 | promoter | '087:XM_040648984:pr | XM_040648984 | GARNL3 | LOC417087 |
| DMC | 17 | 11007025 | 191 | promoter | '087:XM_040648985:pr | XM_040648985 | GARNL3 | LOC417087 |
| DMC | 17 | 11007025 | 191 | promoter | '087:XM_040648989:pr | XM_040648989 | GARNL3 | LOC417087 |
| DMC | 17 | 11007025 | 191 | promoter | '087:XM_040648990:pr | XM_040648990 | GARNL3 | LOC417087 |
| DMC | 17 | 11007025 | 191 | promoter | '087:XM_040648991:pr | XM_040648991 | GARNL3 | LOC417087 |
| DMC | 17 | 11007025 | 191 | promoter | '087:XM_040648992:pr | XM_040648992 | GARNL3 | LOC417087 |
| DMC | 17 | 11007025 | 191 | promoter | '087:XM_040648995:pr | XM_040648995 | GARNL3 | LOC417087 |
| DMC | 17 | 11007025 | 191 | promoter | '087:XM_040648996:pr | XM_040648996 | GARNL3 | LOC417087 |
| DMC | 17 | 11007025 | 191 | promoter | '087:XM_040648997:pr | XM_040648997 | GARNL3 | LOC417087 |
| DMC | 17 | 11007025 | 191 | promoter | '087:XM_040648998:pr | XM_040648998 | GARNL3 | LOC417087 |
| DMC | 17 | 11007025 | 191 | promoter | '087:XM_040648999:pr | XM_040648999 | GARNL3 | LOC417087 |
| DMC | 17 | 11007025 | 191 | promoter | '087:XM_040649004:pr | XM_040649004 | GARNL3 | LOC417087 |
| DMC | 17 | 11007025 | 191 | promoter | '087:XM_040649014:pr | XM_040649014 | GARNL3 | LOC417087 |
| DMC | 17 | 11007025 | 191 | promoter | '087:XM_040649016:pr | XM_040649016 | GARNL3 | LOC417087 |
| DMC | 17 | 11007025 | 191 | promoter | '087:XM_040649021:pr | XM_040649021 | GARNL3 | LOC417087 |
| DMC | 17 | 11007025 | 191 | promoter | '087:XM_040649022:pr | XM_040649022 | GARNL3 | LOC417087 |
| DMC | 17 | 11007025 | 191 | promoter | '087:XM_040649023:pr | XM_040649023 | GARNL3 | LOC417087 |
| DMC | 17 | 11007025 | 191 | promoter | '087:XM_040649024:pr | XM_040649024 | GARNL3 | LOC417087 |
| DMC | 17 | 11007025 | 191 | promoter | '087:XM_046901680:pr | XM_046901680 | GARNL3 | LOC417087 |
| DMC | 17 | 11007025 | 191 | promoter | 059809:TAGAGALT0000( | GAGALT0000001351 | GAGALG000000059 | GAGALG0000000598 |
| DMC | 17 | 11007025 | 191 | introns | 059809:TAGAGALT0000( | GAGALT0000001351 | GAGALG000000059 | GAGALG0000000598 |
| DMC | 17 | 11007025 | 191 | intron1 | 059809:TAGAGALT0000( | GAGALT0000001351 | GAGALG000000059 | GAGALG0000000598 |
| DMC | 17 | 11007025 | 191 | downstream | 5697:INRAGALT000000 | IRAGALT000000122( | RAGALG000000056 | IRAGALG000000056( |
| DMC | 18 | 3228957 | 1850 | promoter | 047772:DAVISGALT0047 | DAVISGALT00477720 | AVISGALG0000477 | AVISGALG00004777 |
| DMC | 18 | 3228957 | 1850 | introns | '329:NM_001030700:in | NM_001030700 | COX10 | LOC417329 |
| DMC | 19 | 7740157 | 4707 | other | NA | NA | NA | NA |
| DMC | 19 | 8216649 | 8112 | introns | i43:XM_015296146:intr | XM_015296146 | BCAS3 | LOC417643 |
| DMC | 19 | 8216649 | 8112 | introns | i43:XM_004946728:intr | XM_004946728 | BCAS3 | LOC417643 |
| DMC | 19 | 8216649 | 8112 | introns | i43:XM_004946729:intr | XM_004946729 | BCAS3 | LOC417643 |
| DMC | 19 | 8216649 | 8112 | introns | i43:XM_004946730:intr | XM_004946730 | BCAS3 | LOC417643 |
| DMC | 19 | 8216649 | 8112 | introns | i43:XM_004946731:intr | XM_004946731 | BCAS3 | LOC417643 |
| DMC | 19 | 8216649 | 8112 | introns | i43:XM_004946733:intr | XM_004946733 | BCAS3 | LOC417643 |
| DMC | 19 | 8216649 | 8112 | introns | i43:XM_015296145:intr | XM_015296145 | BCAS3 | LOC417643 |
| DMC | 19 | 8216649 | 8112 | introns | i43:XM_015296147:intr | XM_015296147 | BCAS3 | LOC417643 |
| DMC | 19 | 8216649 | 8112 | introns | i43:XM_040650516:intr | XM_040650516 | BCAS3 | LOC417643 |
| DMC | 19 | 8216649 | 8112 | introns | i43:XM_040650517:intr | XM_040650517 | BCAS3 | LOC417643 |
| DMC | 19 | 8216649 | 8112 | introns | i43:XM_040650518:intr | XM_040650518 | BCAS3 | LOC417643 |
| DMC | 19 | 8216649 | 8112 | introns | i43:XM_040650519:intr | XM_040650519 | BCAS3 | LOC417643 |
| DMC | 19 | 8216649 | 8112 | introns | i43:XM_040650520:intr | XM_040650520 | BCAS3 | LOC417643 |
| DMC | 19 | 8216649 | 8112 | introns | i43:XM_040650521:intr | XM_040650521 | BCAS3 | LOC417643 |
| DMC | 19 | 8216649 | 8112 | introns | i43:XM_046902724:intr | XM_046902724 | BCAS3 | LOC417643 |
| DMC | 19 | 8216649 | 8112 | introns | 7643:XM_415889:intro | XM_415889 | BCAS3 | LOC417643 |
| DMC | 19 | 8216649 | 8112 | introns | i43:XM_004946732:intr | XM_004946732 | BCAS3 | LOC417643 |
| DMC | 19 | 8216649 | 8112 | introns | i43:XM_040650522:intr | XM_040650522 | BCAS3 | LOC417643 |
| DMC | 19 | 8250062 | 15463 | introns | i43:XM_015296146:intr | XM_015296146 | BCAS3 | LOC417643 |
| DMC | 19 | 8250062 | 15463 | introns | i43:XM_004946728:intr | XM_004946728 | BCAS3 | LOC417643 |
| DMC | 19 | 8250062 | 15463 | introns | i43:XM_004946729:intr | XM_004946729 | BCAS3 | LOC417643 |
| DMC | 19 | 8250062 | 15463 | introns | i43:XM_004946730:intr | XM_004946730 | BCAS3 | LOC417643 |
| DMC | 19 | 8250062 | 15463 | introns | i43:XM_004946731:intr | XM_004946731 | BCAS3 | LOC417643 |
| DMC | 19 | 8250062 | 15463 | introns | i43:XM_004946733:intr | XM_004946733 | BCAS3 | LOC417643 |
| DMC | 19 | 8250062 | 15463 | introns | i43:XM_015296145:intr | XM_015296145 | BCAS3 | LOC417643 |
| DMC | 19 | 8250062 | 15463 | introns | i43:XM_015296147:intr | XM_015296147 | BCAS3 | LOC417643 |
| DMC | 19 | 8250062 | 15463 | introns | i43:XM_040650516:intr | XM_040650516 | BCAS3 | LOC417643 |
| DMC | 19 | 8250062 | 15463 | introns | i43:XM_040650517:intr | XM_040650517 | BCAS3 | LOC417643 |
| DMC | 19 | 8250062 | 15463 | introns | i43:XM_040650518:intr | XM_040650518 | BCAS3 | LOC417643 |
| DMC | 19 | 8250062 | 15463 | introns | i43:XM_040650519:intr | XM_040650519 | BCAS3 | LOC417643 |
| DMC | 19 | 8250062 | 15463 | introns | i43:XM_040650520:intr | XM_040650520 | BCAS3 | LOC417643 |
| DMC | 19 | 8250062 | 15463 | introns | i43:XM_040650521:intr | XM_040650521 | BCAS3 | LOC417643 |
| DMC | 19 | 8250062 | 15463 | introns | i43:XM_046902724:intr | XM_046902724 | BCAS3 | LOC417643 |
| DMC | 19 | 8250062 | 15463 | introns | 7643:XM_415889:intro | XM_415889 | BCAS3 | LOC417643 |
| DMC | 19 | 8250062 | 15463 | introns | i43:XM_004946732:intr | XM_004946732 | BCAS3 | LOC417643 |
| DMC | 19 | 8250062 | 15463 | introns | i43:XM_040650522:intr | XM_040650522 | BCAS3 | LOC417643 |
| DMC | 19 | 8250062 | 15463 | introns | i43:XM_004946735:intr | XM_004946735 | BCAS3 | LOC417643 |
| DMC | 19 | 8250062 | 15463 | introns | i43:XM_040650523:intr | XM_040650523 | BCAS3 | LOC417643 |
| DMC | 19 | 8250062 | 15463 | introns | i43:XM_040650524:intr | XM_040650524 | BCAS3 | LOC417643 |
| DMC | 20 | 2792170 | -5551 | other | NA | NA | NA | NA |

| | | | | | | | | |
|---|---|---|---|---|---|---|---|---|
| DMC | 20 | 9107528 | 5609 | introns | )240:XM_015296646:in | XM_015296646 | NKAIN4 | LOC419240 |
| DMC | 20 | 9107528 | 5609 | introns | )240:XM_025142311:in | XM_025142311 | NKAIN4 | LOC419240 |
| DMC | 20 | 9107528 | 5609 | introns | )240:XM_046903023:in | XM_046903023 | NKAIN4 | LOC419240 |
| DMC | 20 | 9107528 | 5609 | introns | )240:XM_046903022:in | XM_046903022 | NKAIN4 | LOC419240 |
| DMC | 20 | 9107528 | 5609 | introns | )240:XM_015296645:in | XM_015296645 | NKAIN4 | LOC419240 |
| DMC | 20 | 9107528 | 5609 | intron1 | )240:XM_015296646:in | XM_015296646 | NKAIN4 | LOC419240 |
| DMC | 20 | 9107528 | 5609 | intron1 | )240:XM_025142311:in | XM_025142311 | NKAIN4 | LOC419240 |
| DMC | 20 | 9107528 | 5609 | intron1 | )240:XM_046903023:in | XM_046903023 | NKAIN4 | LOC419240 |
| DMC | 20 | 9107528 | 5609 | intron1 | )240:XM_046903022:in | XM_046903022 | NKAIN4 | LOC419240 |
| DMC | 20 | 9107528 | 5609 | intron1 | )240:XM_015296645:in | XM_015296645 | NKAIN4 | LOC419240 |
| DMC | 20 | 11841447 | -234 | promoter | 3166:XM_046902892:pr | XM_046902892 | PMEPA1 | LOC428166 |
| DMC | 20 | 11841447 | -234 | promoter | 166:NM_001031492:pr | NM_001031492 | PMEPA1 | LOC428166 |
| DMC | 20 | 11841447 | -234 | promoter | 3166:XM_040650746:pr | XM_040650746 | PMEPA1 | LOC428166 |
| DMC | 20 | 11841447 | -234 | promoter | 3166:XM_046902890:pr | XM_046902890 | PMEPA1 | LOC428166 |
| DMC | 20 | 11841447 | -234 | promoter | 3166:XM_046902891:pr | XM_046902891 | PMEPA1 | LOC428166 |
| DMC | 20 | 11841447 | -234 | promoter | )90026:TAGAGALT0000 | GAGALT0000002019 | GAGALG000000090 | GAGALG0000000900 |
| DMC | 20 | 11841447 | -234 | introns | 3166:XM_046902892:in | XM_046902892 | PMEPA1 | LOC428166 |
| DMC | 20 | 11841447 | -234 | intron1 | 3166:XM_046902892:in | XM_046902892 | PMEPA1 | LOC428166 |
| DMC | 20 | 13098919 | -61 | promoter | )08982:NONGGAT0144 | NONGGAT014424 | NONGGAG008982 | NONGGAG008982 |
| DMC | 20 | 13098919 | -61 | exons | 751795:XR_006931900: | XR_006931900 | LOC101751795 | LOC101751795 |
| DMC | 20 | 13098919 | -61 | exons | )08982:NONGGAT0144 | NONGGAT014424 | NONGGAG008982 | NONGGAG008982 |
| DMC | 20 | 13098919 | -61 | exon1 | )08982:NONGGAT0144 | NONGGAT014424 | NONGGAG008982 | NONGGAG008982 |
| DMC | 20 | 13098919 | -61 | introns | )341:XM_025142326:in | XM_025142326 | TSHZ2 | LOC419341 |
| DMC | 20 | 13098919 | -61 | introns | )341:XR_005841352:int | XR_005841352 | TSHZ2 | LOC419341 |
| DMC | 20 | 13098919 | -61 | introns | )341:XR_005841353:int | XR_005841353 | TSHZ2 | LOC419341 |
| DMC | 20 | 13098919 | -61 | introns | )341:XR_005841354:int | XR_005841354 | TSHZ2 | LOC419341 |
| DMC | 20 | 13098919 | -61 | introns | )341:XM_025142325:in | XM_025142325 | TSHZ2 | LOC419341 |
| DMC | 20 | 13098919 | -61 | introns | )341:XM_004947254:in | XM_004947254 | TSHZ2 | LOC419341 |
| DMC | 20 | 13098919 | -61 | introns | )341:XM_040651078:in | XM_040651078 | TSHZ2 | LOC419341 |
| DMC | 20 | 13098919 | -61 | intron1 | )341:XM_025142326:in | XM_025142326 | TSHZ2 | LOC419341 |
| DMC | 20 | 13098919 | -61 | intron1 | )341:XR_005841352:int | XR_005841352 | TSHZ2 | LOC419341 |
| DMC | 20 | 13098919 | -61 | intron1 | )341:XR_005841353:int | XR_005841353 | TSHZ2 | LOC419341 |
| DMC | 20 | 13098919 | -61 | intron1 | )341:XR_005841354:int | XR_005841354 | TSHZ2 | LOC419341 |
| DMC | 20 | 13098919 | -61 | intron1 | )341:XM_025142325:in | XM_025142325 | TSHZ2 | LOC419341 |
| DMC | 20 | 13098919 | -61 | intron1 | )341:XM_004947254:in | XM_004947254 | TSHZ2 | LOC419341 |
| DMC | 20 | 13098919 | -61 | intron1 | )341:XM_040651078:in | XM_040651078 | TSHZ2 | LOC419341 |
| DMC | 20 | 13295709 | 6110 | introns | 51795:XR_006931900:i | XR_006931900 | LOC101751795 | LOC101751795 |
| DMC | 20 | 13295709 | 6110 | introns | 51795:XR_001469592:i | XR_001469592 | LOC101751795 | LOC101751795 |
| DMC | 20 | 13295709 | 6110 | introns | 51795:XR_005841355:i | XR_005841355 | LOC101751795 | LOC101751795 |
| DMC | 20 | 13434035 | -249 | promoter | )286:XM_015296420:pr | XM_015296420 | SALL4 | LOC769286 |
| DMC | 20 | 13434035 | -249 | UTR5 | 36:XM_015296420:5'UT | XM_015296420 | SALL4 | LOC769286 |
| DMC | 20 | 13434035 | -249 | exons | 9286:XM_015296420:e | XM_015296420 | SALL4 | LOC769286 |
| DMC | 20 | 13434035 | -249 | exon1 | 9286:XM_015296420:e | XM_015296420 | SALL4 | LOC769286 |
| DMC | 20 | 13434035 | -249 | introns | )286:NM_001080872:in | NM_001080872 | SALL4 | LOC769286 |
| DMC | 20 | 13434035 | -249 | introns | )286:XM_015296421:in | XM_015296421 | SALL4 | LOC769286 |
| DMC | 20 | 13434035 | -249 | introns | )286:XM_046902893:in | XM_046902893 | SALL4 | LOC769286 |
| DMC | 20 | 13434035 | -249 | introns | )286:XM_046902894:in | XM_046902894 | SALL4 | LOC769286 |
| DMC | 20 | 13434035 | -249 | intron1 | )286:NM_001080872:in | NM_001080872 | SALL4 | LOC769286 |
| DMC | 20 | 13434035 | -249 | intron1 | )286:XM_015296421:in | XM_015296421 | SALL4 | LOC769286 |
| DMC | 20 | 13434035 | -249 | intron1 | )286:XM_046902893:in | XM_046902893 | SALL4 | LOC769286 |
| DMC | 20 | 13434035 | -249 | intron1 | )286:XM_046902894:in | XM_046902894 | SALL4 | LOC769286 |
| DMC | 20 | 13466986 | 349 | promoter | )50053:DAVISGALT0050 | AVISGALT00500530 | AVISGALG00005005 | AVISGALG00005005 |
| DMC | 20 | 13466986 | 349 | introns | 345:XR_003071384:intr | XR_003071384 | ATP9A | LOC419345 |
| DMC | 20 | 13466986 | 349 | introns | 345:XM_015296518:int | XM_015296518 | ATP9A | LOC419345 |
| DMC | 20 | 13466986 | 349 | introns | 345:XM_046902905:int | XM_046902905 | ATP9A | LOC419345 |
| DMC | 20 | 13467023 | 312 | promoter | )50053:DAVISGALT0050 | AVISGALT00500530 | AVISGALG00005005 | AVISGALG00005005 |
| DMC | 20 | 13467023 | 312 | introns | 345:XR_003071384:intr | XR_003071384 | ATP9A | LOC419345 |
| DMC | 20 | 13467023 | 312 | introns | 345:XM_015296518:int | XM_015296518 | ATP9A | LOC419345 |
| DMC | 20 | 13467023 | 312 | introns | 345:XM_046902905:int | XM_046902905 | ATP9A | LOC419345 |
| DMC | 20 | 13476693 | -110 | exons | 345:XR_003071384:exo | XR_003071384 | ATP9A | LOC419345 |
| DMC | 20 | 13476693 | -110 | exons | 345:XM_015296518:exo | XM_015296518 | ATP9A | LOC419345 |
| DMC | 20 | 13476693 | -110 | exons | 345:XM_046902905:exo | XM_046902905 | ATP9A | LOC419345 |
| DMC | 20 | 13476693 | -110 | cds | 345:XM_015296518:exo | XM_015296518 | ATP9A | LOC419345 |
| DMC | 20 | 13476693 | -110 | cds | 345:XM_046902905:exo | XM_046902905 | ATP9A | LOC419345 |
| DMC | 20 | 13490448 | 87 | promoter | )50056:DAVISGALT0050 | AVISGALT00500560 | AVISGALG00005005 | AVISGALG00005005 |
| DMC | 20 | 13490448 | 87 | introns | 345:XR_003071384:intr | XR_003071384 | ATP9A | LOC419345 |
| DMC | 20 | 13490448 | 87 | introns | 345:XM_015296518:intr | XM_015296518 | ATP9A | LOC419345 |

| | | | | | | | | |
|---|---|---|---|---|---|---|---|---|
| DMC | 20 | 13490448 | 87 | introns | 345:XM_046902905:intr | XM_046902905 | ATP9A | LOC419345 |
| DMC | 20 | 13511181 | 2967 | promoter | 9346:XM_025142327:pr | XM_025142327 | NFATC2 | LOC419346 |
| DMC | 20 | 13609479 | -3583 | other | NA | NA | NA | NA |
| DMC | 21 | 6060491 | 89 | promoter | 3197:XM_004947521:pr | XM_004947521 | C1QB | LOC428197 |
| DMC | 21 | 6060491 | 89 | exons | 419503:XM_417653:exo | XM_417653 | C1QC | LOC419503 |
| DMC | 21 | 6060491 | 89 | introns | 002918:NONGGAT00498 | NONGGAT004983 | NONGGAG002918 | NONGGAG002918 |
| DMC | 21 | 6060491 | 89 | intron1 | 002918:NONGGAT00498 | NONGGAT004983 | NONGGAG002918 | NONGGAG002918 |
| DMC | 21 | 6060491 | 89 | cds | 419503:XM_417653:exo | XM_417653 | C1QC | LOC419503 |
| DMC | 21 | 6113831 | 61 | exons | 3198:XM_040651421:ex | XM_040651421 | EPHA8 | LOC428198 |
| DMC | 21 | 6113831 | 61 | exons | 3198:XM_040651420:ex | XM_040651420 | EPHA8 | LOC428198 |
| DMC | 21 | 6113831 | 61 | exons | 3198:XM_015297218:ex | XM_015297218 | EPHA8 | LOC428198 |
| DMC | 21 | 6113831 | 61 | exons | 3198:XM_040651417:ex | XM_040651417 | EPHA8 | LOC428198 |
| DMC | 21 | 6113831 | 61 | exons | 3198:XM_040651418:ex | XM_040651418 | EPHA8 | LOC428198 |
| DMC | 21 | 6113831 | 61 | exons | 3198:XM_040651419:ex | XM_040651419 | EPHA8 | LOC428198 |
| DMC | 21 | 6113831 | 61 | cds | 3198:XM_040651421:ex | XM_040651421 | EPHA8 | LOC428198 |
| DMC | 21 | 6113831 | 61 | cds | 3198:XM_040651420:ex | XM_040651420 | EPHA8 | LOC428198 |
| DMC | 21 | 6113831 | 61 | cds | 3198:XM_015297218:ex | XM_015297218 | EPHA8 | LOC428198 |
| DMC | 21 | 6113831 | 61 | cds | 3198:XM_040651417:ex | XM_040651417 | EPHA8 | LOC428198 |
| DMC | 21 | 6113831 | 61 | cds | 3198:XM_040651418:ex | XM_040651418 | EPHA8 | LOC428198 |
| DMC | 21 | 6113831 | 61 | cds | 3198:XM_040651419:ex | XM_040651419 | EPHA8 | LOC428198 |
| DMC | 22 | 2386215 | 932 | promoter | 17353:XR_006931999:p | XR_006931999 | LOC124417353 | LOC124417353 |
| DMC | 22 | 2386215 | 932 | exons | 771:XM_015297425:exo | XM_015297425 | ADGRA2 | LOC426771 |
| DMC | 22 | 2386215 | 932 | exons | 771:XM_015297426:exo | XM_015297426 | ADGRA2 | LOC426771 |
| DMC | 22 | 2386215 | 932 | exons | 771:XM_015297424:exo | XM_015297424 | ADGRA2 | LOC426771 |
| DMC | 22 | 2386215 | 932 | exons | 771:XM_004947584:exo | XM_004947584 | ADGRA2 | LOC426771 |
| DMC | 22 | 2386215 | 932 | UTR3 | '1:XM_015297425:3'UT | XM_015297425 | ADGRA2 | LOC426771 |
| DMC | 22 | 2386215 | 932 | UTR3 | '1:XM_015297426:3'UT | XM_015297426 | ADGRA2 | LOC426771 |
| DMC | 22 | 2386215 | 932 | UTR3 | '1:XM_015297424:3'UT | XM_015297424 | ADGRA2 | LOC426771 |
| DMC | 22 | 2386215 | 932 | UTR3 | '1:XM_004947584:3'UT | XM_004947584 | ADGRA2 | LOC426771 |
| DMC | 22 | 2386215 | 932 | downstream | 6772:XM_424383:down | XM_424383 | BRF2 | LOC426772 |
| DMC | 22 | 2412845 | 101 | promoter | 009017:INRAGALT0000C | NRAGALT00000019O | RAGALG00000009O | RAGALG0000000901 |
| DMC | 22 | 2412845 | 101 | promoter | 54556:XM_040651462: | XM_040651462 | LOC117654556 | LOC117654556 |
| DMC | 22 | 2412845 | 101 | promoter | 0991:XM_040651525:pr | XM_040651525 | ADRB3 | LOC430991 |
| DMC | 22 | 2412845 | 101 | promoter | 30991:XM_428541:pro | XM_428541 | ADRB3 | LOC430991 |
| DMC | 22 | 2412845 | 101 | exons | 0991:XM_040651525:e | XM_040651525 | ADRB3 | LOC430991 |
| DMC | 22 | 2412845 | 101 | exons | 430991:XM_428541:exo | XM_428541 | ADRB3 | LOC430991 |
| DMC | 22 | 2412845 | 101 | exon1 | 0991:XM_040651525:e | XM_040651525 | ADRB3 | LOC430991 |
| DMC | 22 | 2412845 | 101 | exon1 | 430991:XM_428541:exo | XM_428541 | ADRB3 | LOC430991 |
| DMC | 22 | 2412845 | 101 | cds | 0991:XM_040651525:e | XM_040651525 | ADRB3 | LOC430991 |
| DMC | 22 | 2412845 | 101 | cds | 430991:XM_428541:exo | XM_428541 | ADRB3 | LOC430991 |
| DMC | 22 | 2586957 | 37 | exons | 435:XM_025142766:exo | XM_025142766 | PLEKHA2 | LOC395435 |
| DMC | 22 | 2586957 | 37 | exons | 435:XM_046903351:exo | XM_046903351 | PLEKHA2 | LOC395435 |
| DMC | 22 | 2586957 | 37 | exons | 5435:NM_204698:exor | NM_204698 | PLEKHA2 | LOC395435 |
| DMC | 22 | 2586957 | 37 | exons | 435:XM_025142765:exo | XM_025142765 | PLEKHA2 | LOC395435 |
| DMC | 22 | 2586957 | 37 | exons | 435:XM_025142767:exo | XM_025142767 | PLEKHA2 | LOC395435 |
| DMC | 22 | 2586957 | 37 | cds | 435:XM_025142766:exo | XM_025142766 | PLEKHA2 | LOC395435 |
| DMC | 22 | 2586957 | 37 | cds | 435:XM_046903351:exo | XM_046903351 | PLEKHA2 | LOC395435 |
| DMC | 22 | 2586957 | 37 | cds | 5435:NM_204698:exor | NM_204698 | PLEKHA2 | LOC395435 |
| DMC | 22 | 2586957 | 37 | cds | 435:XM_025142765:exo | XM_025142765 | PLEKHA2 | LOC395435 |
| DMC | 22 | 2586957 | 37 | cds | 435:XM_025142767:exo | XM_025142767 | PLEKHA2 | LOC395435 |
| DMC | 22 | 2587154 | -160 | exons | 435:XM_025142766:exo | XM_025142766 | PLEKHA2 | LOC395435 |
| DMC | 22 | 2587154 | -160 | exons | 435:XM_046903351:exo | XM_046903351 | PLEKHA2 | LOC395435 |
| DMC | 22 | 2587154 | -160 | exons | 5435:NM_204698:exor | NM_204698 | PLEKHA2 | LOC395435 |
| DMC | 22 | 2587154 | -160 | exons | 435:XM_025142765:exo | XM_025142765 | PLEKHA2 | LOC395435 |
| DMC | 22 | 2587154 | -160 | exons | 435:XM_025142767:exo | XM_025142767 | PLEKHA2 | LOC395435 |
| DMC | 22 | 2587154 | -160 | UTR3 | 5:XM_025142766:3'UT | XM_025142766 | PLEKHA2 | LOC395435 |
| DMC | 22 | 2587154 | -160 | UTR3 | 5:XM_046903351:3'UT | XM_046903351 | PLEKHA2 | LOC395435 |
| DMC | 22 | 2587154 | -160 | UTR3 | 435:NM_204698:3'UTR( | NM_204698 | PLEKHA2 | LOC395435 |
| DMC | 22 | 2587154 | -160 | UTR3 | 5:XM_025142765:3'UT | XM_025142765 | PLEKHA2 | LOC395435 |
| DMC | 22 | 2587154 | -160 | UTR3 | 5:XM_025142767:3'UT | XM_025142767 | PLEKHA2 | LOC395435 |
| DMC | 23 | 4352825 | 596 | exons | 009158:INRAGALT0000I | NRAGALT00000019O | RAGALG000000091 | RAGALG0000000915 |
| DMC | 23 | 4352825 | 596 | exons | 009158:INRAGALT0000I | NRAGALT00000019O | RAGALG000000091 | RAGALG0000000915 |
| DMC | 23 | 4352825 | 596 | exons | 009158:INRAGALT0000I | NRAGALT0000001941 | RAGALG000000091 | RAGALG0000000915 |
| DMC | 23 | 4352825 | 596 | exons | 009158:INRAGALT0000I | NRAGALT0000001941 | RAGALG000000091 | RAGALG0000000915 |
| DMC | 23 | 4352825 | 596 | introns | 009158:INRAGALT0000C | NRAGALT0000001941 | RAGALG000000091 | RAGALG0000000915 |
| DMC | 23 | 4352825 | 596 | intron1 | 009158:INRAGALT0000C | NRAGALT0000001941 | RAGALG000000091 | RAGALG0000000915 |
| DMC | 23 | 5219377 | -3564 | other | NA | NA | NA | NA |

| | | | | | | | | |
|---|---|---|---|---|---|---|---|---|
| DMC | 23 | 5236064 | 47 | exons | 524:NM_001271988:ex | NM_001271988 | COL9A2 | LOC396524 |
| DMC | 23 | 5236064 | 47 | exons | 524:XM_015297640:ex | XM_015297640 | COL9A2 | LOC396524 |
| DMC | 23 | 5236064 | 47 | cds | 524:NM_001271988:ex | NM_001271988 | COL9A2 | LOC396524 |
| DMC | 23 | 5236064 | 47 | cds | 524:XM_015297640:ex | XM_015297640 | COL9A2 | LOC396524 |
| DMC | 23 | 5262520 | 349 | exons | 051370:DAVISGALT005 | DAVISGALT00513700 | AVISGALG0000513 | AVISGALG00005137 |
| DMC | 23 | 5262520 | 349 | exons | 8226:XM_015297839:e | XM_015297839 | RIMS3 | LOC428226 |
| DMC | 23 | 5262520 | 349 | exons | 8226:XM_040651790:e | XM_040651790 | RIMS3 | LOC428226 |
| DMC | 23 | 5262520 | 349 | exons | 8226:XR_006932058:ex | XR_006932058 | RIMS3 | LOC428226 |
| DMC | 23 | 5262520 | 349 | exons | 8226:XR_006932059:ex | XR_006932059 | RIMS3 | LOC428226 |
| DMC | 23 | 5262520 | 349 | exon1 | 051370:DAVISGALT005 | DAVISGALT00513700 | AVISGALG0000513 | AVISGALG00005137 |
| DMC | 23 | 5262520 | 349 | UTR3 | 26:XM_015297839:3'UT | XM_015297839 | RIMS3 | LOC428226 |
| DMC | 23 | 5262520 | 349 | UTR3 | 26:XM_040651790:3'UT | XM_040651790 | RIMS3 | LOC428226 |
| DMC | 23 | 5353716 | 1010 | promoter | 0477:XM_015297827:pr | XM_015297827 | COL16A1 | LOC430477 |
| DMC | 23 | 5353716 | 1010 | promoter | 0477:XM_015297828:pr | XM_015297828 | COL16A1 | LOC430477 |
| DMC | 23 | 5353716 | 1010 | promoter | 0477:XM_015297829:pr | XM_015297829 | COL16A1 | LOC430477 |
| DMC | 23 | 5353716 | 1010 | promoter | 0477:XM_015297830:pr | XM_015297830 | COL16A1 | LOC430477 |
| DMC | 23 | 5353716 | 1010 | promoter | 0477:XM_015297831:pr | XM_015297831 | COL16A1 | LOC430477 |
| DMC | 23 | 5353716 | 1010 | exons | 163:XM_015297826:ex | XM_015297826 | ADGRB2 | LOC426163 |
| DMC | 23 | 5353716 | 1010 | exons | 163:XM_040651780:ex | XM_040651780 | ADGRB2 | LOC426163 |
| DMC | 23 | 5353716 | 1010 | UTR3 | 63:XM_015297826:3'UT | XM_015297826 | ADGRB2 | LOC426163 |
| DMC | 23 | 5353716 | 1010 | UTR3 | 63:XM_040651780:3'UT | XM_040651780 | ADGRB2 | LOC426163 |
| DMC | 24 | 252858 | 2482 | introns | 067:NM_001398296:int | NM_001398296 | PKNOX2 | LOC374067 |
| DMC | 24 | 252858 | 2482 | introns | 067:NM_001398297:int | NM_001398297 | PKNOX2 | LOC374067 |
| DMC | 24 | 252858 | 2482 | introns | 74067:NM_204226:intr | NM_204226 | PKNOX2 | LOC374067 |
| DMC | 24 | 252858 | 2482 | introns | 067:XM_015297935:int | XM_015297935 | PKNOX2 | LOC374067 |
| DMC | 24 | 267843 | 567 | downstream | 704:XR_006932079:dow | XR_006932079 | SLC37A2 | LOC419704 |
| DMC | 24 | 267843 | 567 | downstream | 04:XM_015298088:dov | XM_015298088 | SLC37A2 | LOC419704 |
| DMC | 24 | 267843 | 567 | downstream | 04:XM_040652268:dov | XM_040652268 | SLC37A2 | LOC419704 |
| DMC | 24 | 267843 | 567 | downstream | 04:XM_040652270:dov | XM_040652270 | SLC37A2 | LOC419704 |
| DMC | 24 | 267843 | 567 | downstream | 04:XM_046903783:dov | XM_046903783 | SLC37A2 | LOC419704 |
| DMC | 24 | 267843 | 567 | downstream | 704:XR_006932078:dow | XR_006932078 | SLC37A2 | LOC419704 |
| DMC | 24 | 403924 | -146 | introns | 710:XM_046903791:int | XM_046903791 | CDON | LOC419710 |
| DMC | 24 | 2431104 | -8248 | other | NA | NA | NA | NA |
| DMC | 25 | 2038202 | 214 | promoter | 07583:XM_040652618: | XM_040652618 | ZBTB7B | LOC121107583 |
| DMC | 25 | 2038202 | 214 | promoter | 07583:XM_046904042: | XM_046904042 | ZBTB7B | LOC121107583 |
| DMC | 25 | 2038202 | 214 | exons | 530287:XM_046904045 | XM_046904045 | LOC112530287 | LOC112530287 |
| DMC | 25 | 2038202 | 214 | exons | 530287:XM_025143466 | XM_025143466 | LOC112530287 | LOC112530287 |
| DMC | 25 | 2038202 | 214 | cds | 530287:XM_046904045 | XM_046904045 | LOC112530287 | LOC112530287 |
| DMC | 25 | 2038202 | 214 | cds | 530287:XM_025143466 | XM_025143466 | LOC112530287 | LOC112530287 |
| DMC | 25 | 2421366 | 25 | promoter | 07593:XM_040652644: | XM_040652644 | LOC121107593 | LOC121107593 |
| DMC | 25 | 2421366 | 25 | exons | 356:XM_015298671:ex | XM_015298671 | CADM3 | LOC772356 |
| DMC | 25 | 2421366 | 25 | exons | 356:XM_015298672:ex | XM_015298672 | CADM3 | LOC772356 |
| DMC | 25 | 2421366 | 25 | exons | 356:XM_015298673:ex | XM_015298673 | CADM3 | LOC772356 |
| DMC | 25 | 2421366 | 25 | exons | 2356:XM_015298674:e | XM_015298674 | CADM3 | LOC772356 |
| DMC | 25 | 2421366 | 25 | exons | 356:XM_040652640:ex | XM_040652640 | CADM3 | LOC772356 |
| DMC | 25 | 2421366 | 25 | exons | 356:XM_040652642:ex | XM_040652642 | CADM3 | LOC772356 |
| DMC | 25 | 2421366 | 25 | exons | 2356:XM_040652643:ex | XM_040652643 | CADM3 | LOC772356 |
| DMC | 25 | 2421366 | 25 | exons | 356:XM_040652641:ex | XM_040652641 | CADM3 | LOC772356 |
| DMC | 25 | 2421366 | 25 | cds | 356:XM_015298671:ex | XM_015298671 | CADM3 | LOC772356 |
| DMC | 25 | 2421366 | 25 | cds | 356:XM_015298672:ex | XM_015298672 | CADM3 | LOC772356 |
| DMC | 25 | 2421366 | 25 | cds | 356:XM_015298673:ex | XM_015298673 | CADM3 | LOC772356 |
| DMC | 25 | 2421366 | 25 | cds | 2356:XM_015298674:e | XM_015298674 | CADM3 | LOC772356 |
| DMC | 25 | 2421366 | 25 | cds | 356:XM_040652640:ex | XM_040652640 | CADM3 | LOC772356 |
| DMC | 25 | 2421366 | 25 | cds | 356:XM_040652642:ex | XM_040652642 | CADM3 | LOC772356 |
| DMC | 25 | 2421366 | 25 | cds | 2356:XM_040652643:e | XM_040652643 | CADM3 | LOC772356 |
| DMC | 25 | 2421366 | 25 | cds | 356:XM_040652641:ex | XM_040652641 | CADM3 | LOC772356 |
| DMC | 25 | 2421366 | 25 | downstream | 2141:TAGAGALT00000 | GAGALT0000002344 | GAGALG000000042 | GAGALG0000000421 |
| DMC | 25 | 2421366 | 25 | downstream | 2141:TAGAGALT00000 | GAGALT0000002344 | GAGALG000000042 | GAGALG0000000421 |
| DMC | 25 | 2439336 | 17 | introns | 454:XM_040652639:int | XM_040652639 | IGSF9 | LOC425454 |
| DMC | 25 | 2506847 | -55 | exons | 51238:XM_025143604: | XM_025143604 | PPOX | LOC107051238 |
| DMC | 25 | 2506847 | -55 | cds | 51238:XM_025143604: | XM_025143604 | PPOX | LOC107051238 |
| DMC | 25 | 2593686 | -127 | introns | 59441:XM_003642666:i | XM_003642666 | CERS2 | LOC100859441 |
| DMC | 25 | 2593686 | -127 | introns | 59441:XM_025143643:i | XM_025143643 | CERS2 | LOC100859441 |
| DMC | 25 | 2593686 | -127 | introns | 59441:XM_046903963:i | XM_046903963 | CERS2 | LOC100859441 |
| DMC | 25 | 2608624 | 6 | promoter | 5847:XM_046903971:pr | XM_046903971 | MLLT11 | LOC395847 |
| DMC | 25 | 2608624 | 6 | promoter | 5847:XR_005841846:pr | XR_005841846 | MLLT11 | LOC395847 |
| DMC | 25 | 2608624 | 6 | exons | 57323:NM_001396641 | NM_001396641 | CDC42SE1 | LOC100857323 |

| | | | | | | | | |
|---|---|---|---|---|---|---|---|---|
| DMC | 25 | 2608624 | 6 | exons | 357323:XM_040652658 | XM_040652658 | CDC42SE1 | LOC100857323 |
| DMC | 25 | 2608624 | 6 | exons | 357323:XR_005841847: | XR_005841847 | CDC42SE1 | LOC100857323 |
| DMC | 25 | 2608624 | 6 | cds | 57323:NM_001396641 | NM_001396641 | CDC42SE1 | LOC100857323 |
| DMC | 25 | 2608624 | 6 | cds | 357323:XM_040652658 | XM_040652658 | CDC42SE1 | LOC100857323 |
| DMC | 25 | 2626556 | -130 | introns | 07596:XR_005841848:i | XR_005841848 | LOC121107596 | LOC121107596 |
| DMC | 25 | 2626556 | -130 | introns | 59892:XM_040652653:i | XM_040652653 | SEMA6C | LOC100859892 |
| DMC | 25 | 2626556 | -130 | introns | 9892:XM_025143612:ir | XM_025143612 | SEMA6C | LOC100859892 |
| DMC | 25 | 2626556 | -130 | introns | 9892:XM_046903966:ir | XM_046903966 | SEMA6C | LOC100859892 |
| DMC | 25 | 2626556 | -130 | introns | 9892:XM_015280041:ir | XM_015280041 | SEMA6C | LOC100859892 |
| DMC | 25 | 2626556 | -130 | intron1 | 07596:XR_005841848:i | XR_005841848 | LOC121107596 | LOC121107596 |
| DMC | 25 | 2637225 | -128 | introns | 95547:NM_204774:intr | NM_204774 | TMOD4 | LOC395547 |
| DMC | 25 | 2639051 | -96 | promoter | 9983:NM_001142440:pr | NM_001142440 | PIP5K1A | LOC429983 |
| DMC | 25 | 2639051 | -96 | promoter | 95547:NM_204774:pro | NM_204774 | TMOD4 | LOC395547 |
| DMC | 25 | 2639051 | -96 | exons | 425660:XM_423393:exo | XM_423393 | VPS72 | LOC425660 |
| DMC | 25 | 2639051 | -96 | cds | 425660:XM_423393:exo | XM_423393 | VPS72 | LOC425660 |
| DMC | 25 | 2639369 | -23 | promoter | 9983:NM_001142440:pr | NM_001142440 | PIP5K1A | LOC429983 |
| DMC | 25 | 2639369 | -23 | promoter | 95547:NM_204774:pro | NM_204774 | TMOD4 | LOC395547 |
| DMC | 25 | 2639369 | -23 | introns | 25660:XM_423393:intr | XM_423393 | VPS72 | LOC425660 |
| DMC | 25 | 2662686 | -135 | promoter | 9831:XM_025143610:pr | XM_025143610 | ZNF687 | LOC429831 |
| DMC | 25 | 2662686 | -135 | promoter | 29831:XM_427387:pro | XM_427387 | ZNF687 | LOC429831 |
| DMC | 25 | 2662686 | -135 | introns | 16364:NM_001142390: | NM_001142390 | PSMD4 | LOC100216364 |
| DMC | 25 | 2663056 | -46 | promoter | 9831:XM_025143610:pr | XM_025143610 | ZNF687 | LOC429831 |
| DMC | 25 | 2663056 | -46 | promoter | 29831:XM_427387:pro | XM_427387 | ZNF687 | LOC429831 |
| DMC | 25 | 2663056 | -46 | exons | 16364:NM_001142390: | NM_001142390 | PSMD4 | LOC100216364 |
| DMC | 25 | 2663056 | -46 | cds | 16364:NM_001142390: | NM_001142390 | PSMD4 | LOC100216364 |
| DMC | 25 | 2699544 | -238 | promoter | 57912:XM_003642698: | XM_003642698 | RFX5 | LOC100857912 |
| DMC | 25 | 2699544 | -238 | introns | 662:XM_025143601:int | XM_025143601 | LOC425662 | LOC425662 |
| DMC | 25 | 2699544 | -238 | introns | 662:XM_025143603:int | XM_025143603 | LOC425662 | LOC425662 |
| DMC | 25 | 2699544 | -238 | introns | 662:XM_025143602:int | XM_025143602 | LOC425662 | LOC425662 |
| DMC | 25 | 2699544 | -238 | downstream | 9407:INRAGALT000000 | IRAGALT0000001979 | RAGALG000000094 | IRAGALG0000000940 |
| DMC | 25 | 2701419 | -77 | introns | 662:XM_025143601:int | XM_025143601 | LOC425662 | LOC425662 |
| DMC | 25 | 2701419 | -77 | introns | 662:XM_025143603:int | XM_025143603 | LOC425662 | LOC425662 |
| DMC | 25 | 2701419 | -77 | introns | 662:XM_025143602:int | XM_025143602 | LOC425662 | LOC425662 |
| DMC | 25 | 2772551 | -320 | promoter | 2466695:NR_105488:pr | NR_105488 | MIR6620 | LOC102466695 |
| DMC | 25 | 2772551 | -320 | promoter | :unassigned_transcript | ssigned_transcript_1 | MIR6620 | LOC102466695 |
| DMC | 25 | 2772551 | -320 | exons | 665:NM_001347391:ex | NM_001347391 | CGN | LOC425665 |
| DMC | 25 | 2772551 | -320 | cds | 665:NM_001347391:ex | NM_001347391 | CGN | LOC425665 |
| DMC | 26 | 505285 | 1901 | introns | 173:XM_003642732:int | XM_003642732 | LGR6 | LOC421173 |
| DMC | 26 | 505285 | 1901 | introns | 173:XM_004934825:int | XM_004934825 | LGR6 | LOC421173 |
| DMC | 26 | 505285 | 1901 | introns | 173:XM_004934826:int | XM_004934826 | LGR6 | LOC421173 |
| DMC | 26 | 513602 | 733 | promoter | 09431:INRAGALT00000 | IRAGALT0000001983 | RAGALG000000094 | IRAGALG0000000943 |
| DMC | 26 | 513602 | 733 | introns | .73:XM_003642732:intr | XM_003642732 | LGR6 | LOC421173 |
| DMC | 26 | 513602 | 733 | introns | .73:XM_004934825:intr | XM_004934825 | LGR6 | LOC421173 |
| DMC | 26 | 513602 | 733 | introns | .73:XM_004934826:intr | XM_004934826 | LGR6 | LOC421173 |
| DMC | 26 | 513646 | 689 | promoter | 09431:INRAGALT00000 | IRAGALT0000001983 | RAGALG000000094 | IRAGALG0000000943 |
| DMC | 26 | 513646 | 689 | introns | .73:XM_003642732:intr | XM_003642732 | LGR6 | LOC421173 |
| DMC | 26 | 513646 | 689 | introns | .73:XM_004934825:intr | XM_004934825 | LGR6 | LOC421173 |
| DMC | 26 | 513646 | 689 | introns | .73:XM_004934826:intr | XM_004934826 | LGR6 | LOC421173 |
| DMC | 26 | 535915 | 29 | promoter | .171:XM_040652763:pr | XM_040652763 | SKIV2L | LOC421171 |
| DMC | 26 | 535915 | 29 | exons | 59071:XM_015298767: | XM_015298767 | NELFE | LOC100859071 |
| DMC | 26 | 535915 | 29 | cds | 59071:XM_015298767: | XM_015298767 | NELFE | LOC100859071 |
| DMC | 26 | 645548 | -323 | introns | .66:XM_025143929:intr | XM_025143929 | LZTR1 | LOC421166 |
| DMC | 26 | 645548 | -323 | introns | .66:XM_040652791:intr | XM_040652791 | LZTR1 | LOC421166 |
| DMC | 26 | 663159 | 1436 | UTR5 | 62:XM_040652757:5'UT | XM_040652757 | NAV1 | LOC430162 |
| DMC | 26 | 663159 | 1436 | UTR5 | 62:XM_040652758:5'UT | XM_040652758 | NAV1 | LOC430162 |
| DMC | 26 | 663159 | 1436 | UTR5 | 62:XM_040652759:5'UT | XM_040652759 | NAV1 | LOC430162 |
| DMC | 26 | 663159 | 1436 | UTR5 | 62:XM_040652760:5'UT | XM_040652760 | NAV1 | LOC430162 |
| DMC | 26 | 663159 | 1436 | UTR5 | 62:XM_040652761:5'UT | XM_040652761 | NAV1 | LOC430162 |
| DMC | 26 | 663159 | 1436 | exons | 0162:XM_040652757:ex | XM_040652757 | NAV1 | LOC430162 |
| DMC | 26 | 663159 | 1436 | exons | 0162:XM_040652758:ex | XM_040652758 | NAV1 | LOC430162 |
| DMC | 26 | 663159 | 1436 | exons | 0162:XM_040652759:ex | XM_040652759 | NAV1 | LOC430162 |
| DMC | 26 | 663159 | 1436 | exons | 0162:XM_040652760:ex | XM_040652760 | NAV1 | LOC430162 |
| DMC | 26 | 663159 | 1436 | exons | 0162:XM_040652761:ex | XM_040652761 | NAV1 | LOC430162 |
| DMC | 26 | 663159 | 1436 | exon1 | 0162:XM_040652757:ex | XM_040652757 | NAV1 | LOC430162 |
| DMC | 26 | 663159 | 1436 | exon1 | 0162:XM_040652758:ex | XM_040652758 | NAV1 | LOC430162 |
| DMC | 26 | 663159 | 1436 | exon1 | 0162:XM_040652759:ex | XM_040652759 | NAV1 | LOC430162 |
| DMC | 26 | 663159 | 1436 | exon1 | 0162:XM_040652760:ex | XM_040652760 | NAV1 | LOC430162 |

| | | | | | | | | |
|---|---|---|---|---|---|---|---|---|
| DMC | 26 | 663159 | 1436 | exon1 | 0162:XM_040652761:ex | XM_040652761 | NAV1 | LOC430162 |
| DMC | 26 | 663159 | 1436 | introns | 162:XM_040652762:int | XM_040652762 | NAV1 | LOC430162 |
| DMC | 26 | 663159 | 1436 | introns | 162:XM_046904162:int | XM_046904162 | NAV1 | LOC430162 |
| DMC | 26 | 663159 | 1436 | intron1 | 162:XM_040652762:int | XM_040652762 | NAV1 | LOC430162 |
| DMC | 26 | 663159 | 1436 | intron1 | 162:XM_046904162:int | XM_046904162 | NAV1 | LOC430162 |
| DMC | 26 | 820746 | 211 | UTR5 | 0283:XM_003642734:5'U | XM_003642734 | PHLDA3 | LOC100859283 |
| DMC | 26 | 820746 | 211 | exons | 359283:XM_003642734 | XM_003642734 | PHLDA3 | LOC100859283 |
| DMC | 26 | 820746 | 211 | exon1 | 359283:XM_003642734 | XM_003642734 | PHLDA3 | LOC100859283 |
| DMC | 26 | 836104 | 876 | introns | 160:XM_040652816:int | XM_040652816 | LAD1 | LOC421160 |
| DMC | 26 | 836104 | 876 | introns | 160:XM_025143836:int | XM_025143836 | LAD1 | LOC421160 |
| DMC | 26 | 859885 | 274 | promoter | 027768:ENSGALT00010( | NSGALT0001006729 | NSGALG000100277 | NSGALG0001002776 |
| DMC | 26 | 859885 | 274 | promoter | 027832:ENSGALT00010( | NSGALT0001006744 | NSGALG000100278 | NSGALG0001002783 |
| DMC | 26 | 859885 | 274 | promoter | 027832:ENSGALT00010( | NSGALT0001006744 | NSGALG000100278 | NSGALG0001002783 |
| DMC | 26 | 859885 | 274 | promoter | 027832:ENSGALT00010( | NSGALT0001006744 | NSGALG000100278 | NSGALG0001002783 |
| DMC | 26 | 859885 | 274 | introns | 027832:ENSGALT00010( | NSGALT0001006744 | NSGALG000100278 | NSGALG0001002783 |
| DMC | 26 | 859885 | 274 | introns | 027832:ENSGALT00010( | NSGALT0001006744 | NSGALG000100278 | NSGALG0001002783 |
| DMC | 26 | 859885 | 274 | introns | 027832:ENSGALT00010( | NSGALT0001006744 | NSGALG000100278 | NSGALG0001002783 |
| DMC | 26 | 859885 | 274 | intron1 | 027832:ENSGALT00010( | NSGALT0001006744 | NSGALG000100278 | NSGALG0001002783 |
| DMC | 26 | 859885 | 274 | intron1 | 027832:ENSGALT00010( | NSGALT0001006744 | NSGALG000100278 | NSGALG0001002783 |
| DMC | 26 | 859885 | 274 | intron1 | 027832:ENSGALT00010( | NSGALT0001006744 | NSGALG000100278 | NSGALG0001002783 |
| DMC | 26 | 1056445 | 110 | promoter | 034608:TAGAGALT0000( | GAGALT0000002362 | GAGALG000000034 | GAGALG0000000346 |
| DMC | 26 | 1056445 | 110 | promoter | 009090:NONGGAT0145 | NONGGAT014541 | NONGGAG009090 | NONGGAG009090 |
| DMC | 26 | 1056445 | 110 | exons | 154:NM_001031028:ex | NM_001031028 | MYBPHL | LOC421154 |
| DMC | 26 | 1056445 | 110 | cds | 154:NM_001031028:ex | NM_001031028 | MYBPHL | LOC421154 |
| DMC | 27 | 2054956 | 741 | promoter | 9950:XM_046904387:pr | XM_046904387 | MRC2 | LOC419950 |
| DMC | 27 | 2054956 | 741 | promoter | 9950:XM_046904388:pr | XM_046904388 | MRC2 | LOC419950 |
| DMC | 27 | 2651262 | -1425 | UTR5 | 32:XM_025143962:5'UT | XM_025143962 | SLC4A1 | LOC396532 |
| DMC | 27 | 2651262 | -1425 | exons | 6532:XM_025143962:ex | XM_025143962 | SLC4A1 | LOC396532 |
| DMC | 27 | 2651262 | -1425 | exon1 | 6532:XM_025143962:ex | XM_025143962 | SLC4A1 | LOC396532 |
| DMC | 27 | 2651262 | -1425 | introns | 532:XM_025143963:int | XM_025143963 | SLC4A1 | LOC396532 |
| DMC | 27 | 2651262 | -1425 | introns | 532:XM_025143964:int | XM_025143964 | SLC4A1 | LOC396532 |
| DMC | 27 | 2651262 | -1425 | introns | 532:XM_025143965:int | XM_025143965 | SLC4A1 | LOC396532 |
| DMC | 27 | 2651262 | -1425 | introns | 532:XM_025143966:int | XM_025143966 | SLC4A1 | LOC396532 |
| DMC | 27 | 2651262 | -1425 | introns | 532:XM_046904342:int | XM_046904342 | SLC4A1 | LOC396532 |
| DMC | 27 | 2651262 | -1425 | introns | 532:NM_001290554:int | NM_001290554 | SLC4A1 | LOC396532 |
| DMC | 27 | 2651262 | -1425 | intron1 | 532:XM_025143963:int | XM_025143963 | SLC4A1 | LOC396532 |
| DMC | 27 | 2651262 | -1425 | intron1 | 532:XM_025143964:int | XM_025143964 | SLC4A1 | LOC396532 |
| DMC | 27 | 2651262 | -1425 | intron1 | 532:XM_025143965:int | XM_025143965 | SLC4A1 | LOC396532 |
| DMC | 27 | 2651262 | -1425 | intron1 | 532:XM_025143966:int | XM_025143966 | SLC4A1 | LOC396532 |
| DMC | 27 | 2651262 | -1425 | intron1 | 532:XM_046904342:int | XM_046904342 | SLC4A1 | LOC396532 |
| DMC | 27 | 2651262 | -1425 | intron1 | 532:NM_001290554:int | NM_001290554 | SLC4A1 | LOC396532 |
| DMC | 27 | 2653657 | 103 | promoter | 6532:XM_040653146:pr | XM_040653146 | SLC4A1 | LOC396532 |
| DMC | 27 | 2653657 | 103 | introns | 532:XM_025143962:int | XM_025143962 | SLC4A1 | LOC396532 |
| DMC | 27 | 2653657 | 103 | introns | 532:XM_025143963:int | XM_025143963 | SLC4A1 | LOC396532 |
| DMC | 27 | 2653657 | 103 | introns | 532:XM_025143964:int | XM_025143964 | SLC4A1 | LOC396532 |
| DMC | 27 | 2653657 | 103 | introns | 532:XM_025143965:int | XM_025143965 | SLC4A1 | LOC396532 |
| DMC | 27 | 2653657 | 103 | introns | 532:XM_025143966:int | XM_025143966 | SLC4A1 | LOC396532 |
| DMC | 27 | 2653657 | 103 | introns | 532:XM_046904342:int | XM_046904342 | SLC4A1 | LOC396532 |
| DMC | 27 | 2653657 | 103 | introns | 532:NM_001290554:int | NM_001290554 | SLC4A1 | LOC396532 |
| DMC | 27 | 2668544 | -962 | promoter | 59792:XM_025144215: | XM_025144215 | UBTF | LOC100859792 |
| DMC | 27 | 2668544 | -962 | promoter | 59792:XM_025144211: | XM_025144211 | UBTF | LOC100859792 |
| DMC | 27 | 2668544 | -962 | promoter | 59792:XM_025144212: | XM_025144212 | UBTF | LOC100859792 |
| DMC | 27 | 2668544 | -962 | promoter | 59792:XM_025144217: | XM_025144217 | UBTF | LOC100859792 |
| DMC | 27 | 2668544 | -962 | promoter | 59792:XM_040653266: | XM_040653266 | UBTF | LOC100859792 |
| DMC | 27 | 2668544 | -962 | promoter | 019748:INRAGALT00000 | NRAGALT0000004362 | RAGALG000000197 | RAGALG000001974 |
| DMC | 27 | 2668544 | -962 | UTR5 | 0792:XM_046904412:5'U | XM_046904412 | UBTF | LOC100859792 |
| DMC | 27 | 2668544 | -962 | exons | 532:XM_025143962:ex | XM_025143962 | SLC4A1 | LOC396532 |
| DMC | 27 | 2668544 | -962 | exons | 532:XM_025143963:ex | XM_025143963 | SLC4A1 | LOC396532 |
| DMC | 27 | 2668544 | -962 | exons | 532:XM_025143964:ex | XM_025143964 | SLC4A1 | LOC396532 |
| DMC | 27 | 2668544 | -962 | exons | 532:XM_025143965:ex | XM_025143965 | SLC4A1 | LOC396532 |
| DMC | 27 | 2668544 | -962 | exons | 532:XM_025143966:ex | XM_025143966 | SLC4A1 | LOC396532 |
| DMC | 27 | 2668544 | -962 | exons | 532:XM_046904342:ex | XM_046904342 | SLC4A1 | LOC396532 |
| DMC | 27 | 2668544 | -962 | exons | 532:NM_001290554:ex | NM_001290554 | SLC4A1 | LOC396532 |
| DMC | 27 | 2668544 | -962 | exons | 532:XM_040653146:ex | XM_040653146 | SLC4A1 | LOC396532 |
| DMC | 27 | 2668544 | -962 | exons | 59792:XM_046904412: | XM_046904412 | UBTF | LOC100859792 |
| DMC | 27 | 2668544 | -962 | exon1 | 59792:XM_046904412: | XM_046904412 | UBTF | LOC100859792 |
| DMC | 27 | 2668544 | -962 | UTR3 | 2:XM_025143962:3'UT | XM_025143962 | SLC4A1 | LOC396532 |

| | | | | | | | | |
|---|---|---|---|---|---|---|---|---|
| DMC | 27 | 2668544 | -962 | UTR3 | ;2:XM_025143963:3'UT | XM_025143963 | SLC4A1 | LOC396532 |
| DMC | 27 | 2668544 | -962 | UTR3 | ;2:XM_025143964:3'UT | XM_025143964 | SLC4A1 | LOC396532 |
| DMC | 27 | 2668544 | -962 | UTR3 | ;2:XM_025143965:3'UT | XM_025143965 | SLC4A1 | LOC396532 |
| DMC | 27 | 2668544 | -962 | UTR3 | ;2:XM_025143966:3'UT | XM_025143966 | SLC4A1 | LOC396532 |
| DMC | 27 | 2668544 | -962 | UTR3 | ;2:XM_046904342:3'UT | XM_046904342 | SLC4A1 | LOC396532 |
| DMC | 27 | 2668544 | -962 | UTR3 | 2:NM_001290554:3'UT | NM_001290554 | SLC4A1 | LOC396532 |
| DMC | 27 | 2668544 | -962 | UTR3 | ;2:XM_040653146:3'UT | XM_040653146 | SLC4A1 | LOC396532 |
| DMC | 27 | 2682933 | 86 | introns | 9792:XM_046904412:ir | XM_046904412 | UBTF | LOC100859792 |
| DMC | 27 | 2682933 | 86 | introns | 9792:XM_025144215:ir | XM_025144215 | UBTF | LOC100859792 |
| DMC | 27 | 2682933 | 86 | introns | 9792:XM_025144211:ir | XM_025144211 | UBTF | LOC100859792 |
| DMC | 27 | 2682933 | 86 | introns | 9792:XM_025144212:ir | XM_025144212 | UBTF | LOC100859792 |
| DMC | 27 | 2682933 | 86 | introns | 9792:XM_025144217:ir | XM_025144217 | UBTF | LOC100859792 |
| DMC | 27 | 2682933 | 86 | introns | 9792:XM_040653266:ir | XM_040653266 | UBTF | LOC100859792 |
| DMC | 27 | 2682933 | 86 | introns | 9792:XM_025144213:ir | XM_025144213 | UBTF | LOC100859792 |
| DMC | 27 | 2682933 | 86 | introns | 9792:XM_040653265:ir | XM_040653265 | UBTF | LOC100859792 |
| DMC | 27 | 2682933 | 86 | introns | 9792:XM_040653268:ir | XM_040653268 | UBTF | LOC100859792 |
| DMC | 27 | 2682933 | 86 | introns | 9792:XM_025144214:ir | XM_025144214 | UBTF | LOC100859792 |
| DMC | 27 | 2699906 | -939 | promoter | )06543:NONGGAT0113! | NONGGAT011390 | NONGGAG006543 | NONGGAG006543 |
| DMC | 27 | 2699906 | -939 | promoter | 49570:NM_001361182: | NM_001361182 | LOC107049570 | LOC107049570 |
| DMC | 27 | 2699906 | -939 | downstream | 9387:XM_003643812:d | XM_003643812 | TMEM101 | LOC100859387 |
| DMC | 27 | 2699906 | -939 | downstream | 4934:TAGAGALT00000( | GAGALT0000002456 | GAGALG000000014 | GAGALG0000000149 |
| DMC | 27 | 2742932 | -2 | exons | 532:NM_001396622:ex | NM_001396622 | COL1A1 | LOC395532 |
| DMC | 27 | 2742932 | -2 | cds | 532:NM_001396622:ex | NM_001396622 | COL1A1 | LOC395532 |
| DMC | 27 | 3140593 | 528 | promoter | 95533:NM_204765:pro | NM_204765 | MEOX1 | LOC395533 |
| DMC | 27 | 3275156 | 6864 | introns | 95953:NM_205071:intro | NM_205071 | IGF2BP1 | LOC395953 |
| DMC | 27 | 3839725 | 357 | promoter | 57564:XM_040653320: | XM_040653320 | LOC107057564 | LOC107057564 |
| DMC | 27 | 3839725 | 357 | promoter | 57564:XM_040653321: | XM_040653321 | LOC107057564 | LOC107057564 |
| DMC | 27 | 3839725 | 357 | promoter | 57564:XM_040653322: | XM_040653322 | LOC107057564 | LOC107057564 |
| DMC | 27 | 3839725 | 357 | promoter | 57564:XM_040653323: | XM_040653323 | LOC107057564 | LOC107057564 |
| DMC | 27 | 3839725 | 357 | promoter | 57564:XR_005841980:p | XR_005841980 | LOC107057564 | LOC107057564 |
| DMC | 27 | 3839725 | 357 | promoter | 57564:XR_005841981:p | XR_005841981 | LOC107057564 | LOC107057564 |
| DMC | 27 | 3839725 | 357 | promoter | 55290:XM_040653234: | XM_040653234 | GPR179 | LOC107055290 |
| DMC | 27 | 3839725 | 357 | exons | 55290:XM_040653234: | XM_040653234 | GPR179 | LOC107055290 |
| DMC | 27 | 3839725 | 357 | exon1 | 55290:XM_040653234: | XM_040653234 | GPR179 | LOC107055290 |
| DMC | 27 | 3839725 | 357 | cds | 55290:XM_040653234: | XM_040653234 | GPR179 | LOC107055290 |
| DMC | 27 | 4125355 | -126 | promoter | .869:XM_004948580:pr | XM_004948580 | FBXL20 | LOC771869 |
| DMC | 27 | 4125355 | -126 | promoter | .869:XM_040653336:pr | XM_040653336 | FBXL20 | LOC771869 |
| DMC | 27 | 4125355 | -126 | UTR5 | 59:XM_004948580:5'UT | XM_004948580 | FBXL20 | LOC771869 |
| DMC | 27 | 4125355 | -126 | UTR5 | 59:XM_040653336:5'UT | XM_040653336 | FBXL20 | LOC771869 |
| DMC | 27 | 4125355 | -126 | exons | .869:XM_004948580:ex | XM_004948580 | FBXL20 | LOC771869 |
| DMC | 27 | 4125355 | -126 | exons | .869:XM_040653336:ex | XM_040653336 | FBXL20 | LOC771869 |
| DMC | 27 | 4125355 | -126 | exon1 | .869:XM_004948580:ex | XM_004948580 | FBXL20 | LOC771869 |
| DMC | 27 | 4125355 | -126 | exon1 | .869:XM_040653336:ex | XM_040653336 | FBXL20 | LOC771869 |
| DMC | 27 | 4125355 | -126 | introns | 869:XM_015299535:int | XM_015299535 | FBXL20 | LOC771869 |
| DMC | 27 | 4125355 | -126 | introns | 869:XM_015299533:int | XM_015299533 | FBXL20 | LOC771869 |
| DMC | 27 | 4125355 | -126 | introns | 869:XM_015299534:int | XM_015299534 | FBXL20 | LOC771869 |
| DMC | 27 | 4125355 | -126 | intron1 | 869:XM_015299535:int | XM_015299535 | FBXL20 | LOC771869 |
| DMC | 27 | 4125355 | -126 | intron1 | 869:XM_015299533:int | XM_015299533 | FBXL20 | LOC771869 |
| DMC | 27 | 4125355 | -126 | intron1 | 869:XM_015299534:int | XM_015299534 | FBXL20 | LOC771869 |
| DMC | 27 | 4171972 | 1430 | promoter | )09698:INRAGALT00000 | IRAGALT0000002056 | RAGALG000000096 | RAGALG000000096 |
| DMC | 27 | 4171972 | 1430 | promoter | 30403:XR_003072032:p | XR_003072032 | LOC112530403 | LOC112530403 |
| DMC | 27 | 4171972 | 1430 | promoter | )26357:ENSGALT00010( | NSGALT0001006413 | NSGALG00010026 | NSGALG0001002635 |
| DMC | 27 | 4761042 | 639 | promoter | )03240:NONGGAT0055 | NONGGAT005563 | NONGGAG003240 | NONGGAG003240 |
| DMC | 27 | 4761042 | 639 | exons | )0031:XM_004948613:e | XM_004948613 | ZNF385C | LOC420031 |
| DMC | 27 | 4761042 | 639 | exons | )0031:XM_046904514:e | XM_046904514 | ZNF385C | LOC420031 |
| DMC | 27 | 4761042 | 639 | exons | )0031:XM_046904515:e | XM_046904515 | ZNF385C | LOC420031 |
| DMC | 27 | 4761042 | 639 | UTR3 | 31:XM_004948613:3'U | XM_004948613 | ZNF385C | LOC420031 |
| DMC | 27 | 4761042 | 639 | UTR3 | 31:XM_046904514:3'U | XM_046904514 | ZNF385C | LOC420031 |
| DMC | 27 | 4761042 | 639 | UTR3 | 31:XM_046904515:3'U | XM_046904515 | ZNF385C | LOC420031 |
| DMC | 27 | 4770569 | 428 | introns | )0031:XM_004948613:in | XM_004948613 | ZNF385C | LOC420031 |
| DMC | 27 | 4770569 | 428 | introns | )0031:XM_046904514:in | XM_046904514 | ZNF385C | LOC420031 |
| DMC | 27 | 4770569 | 428 | introns | )0031:XM_046904515:in | XM_046904515 | ZNF385C | LOC420031 |
| DMC | 27 | 4775459 | -219 | exons | )0030:XM_046904511:e | XM_046904511 | LOC420030 | LOC420030 |
| DMC | 27 | 4775459 | -219 | exons | )0030:XM_046904512:e | XM_046904512 | LOC420030 | LOC420030 |
| DMC | 27 | 4775459 | -219 | introns | )0031:XM_004948613:in | XM_004948613 | ZNF385C | LOC420031 |
| DMC | 27 | 4775459 | -219 | introns | )0031:XM_046904514:in | XM_046904514 | ZNF385C | LOC420031 |
| DMC | 27 | 4775459 | -219 | introns | )0031:XM_046904515:in | XM_046904515 | ZNF385C | LOC420031 |

| | | | | | | | | |
|---|---|---|---|---|---|---|---|---|
| DMC | 27 | 4775459 | -219 | intron1 | )031:XM_004948613:in | XM_004948613 | ZNF385C | LOC420031 |
| DMC | 27 | 4775459 | -219 | cds | )030:XM_046904511:e | XM_046904511 | LOC420030 | LOC420030 |
| DMC | 27 | 4775459 | -219 | cds | )030:XM_046904512:e | XM_046904512 | LOC420030 | LOC420030 |
| DMC | 27 | 4810731 | 1916 | introns | )031:XM_046904514:in | XM_046904514 | ZNF385C | LOC420031 |
| DMC | 27 | 4810731 | 1916 | introns | )031:XM_046904515:in | XM_046904515 | ZNF385C | LOC420031 |
| DMC | 27 | 4810731 | 1916 | intron1 | )031:XM_046904514:in | XM_046904514 | ZNF385C | LOC420031 |
| DMC | 27 | 4810731 | 1916 | intron1 | )031:XM_046904515:in | XM_046904515 | ZNF385C | LOC420031 |
| DMC | 27 | 4811439 | 1208 | introns | )031:XM_046904514:in | XM_046904514 | ZNF385C | LOC420031 |
| DMC | 27 | 4811439 | 1208 | introns | )031:XM_046904515:in | XM_046904515 | ZNF385C | LOC420031 |
| DMC | 27 | 4811439 | 1208 | intron1 | )031:XM_046904514:in | XM_046904514 | ZNF385C | LOC420031 |
| DMC | 27 | 4811439 | 1208 | intron1 | )031:XM_046904515:in | XM_046904515 | ZNF385C | LOC420031 |
| DMC | 27 | 4818021 | -467 | promoter | )26466:ENSGALT00010( | NSGALT0001006433 | NSGALG000100264 | NSGALG0001002646 |
| DMC | 27 | 4818021 | -467 | exons | )26466:ENSGALT00010 | NSGALT0001006433 | NSGALG000100264 | NSGALG0001002646 |
| DMC | 27 | 4818021 | -467 | exons | 58653:XM_004948616:e | XM_004948616 | DHX58 | LOC100858653 |
| DMC | 27 | 4818021 | -467 | exon1 | )26466:ENSGALT00010 | NSGALT0001006433 | NSGALG000100264 | NSGALG0001002646 |
| DMC | 27 | 4818021 | -467 | UTR3 | 653:XM_004948616:3'U | XM_004948616 | DHX58 | LOC100858653 |
| DMC | 28 | 1376882 | 53 | promoter | )28654:ENSGALT00010( | NSGALT0001006936 | NSGALG000100286 | NSGALG0001002865 |
| DMC | 28 | 1376882 | 53 | promoter | )53271:DAVISGALT0053 | AVISGALT00532710 | AVISGALG0000532 | AVISGALG00005327 |
| DMC | 28 | 1376882 | 53 | introns | )76:XM_015299784:int | XM_015299784 | MATK | LOC769076 |
| DMC | 28 | 1376882 | 53 | introns | )76:XM_015299785:int | XM_015299785 | MATK | LOC769076 |
| DMC | 28 | 1376882 | 53 | introns | )76:XM_015299786:int | XM_015299786 | MATK | LOC769076 |
| DMC | 28 | 1376882 | 53 | introns | )76:XM_040653622:int | XM_040653622 | MATK | LOC769076 |
| DMC | 28 | 1376882 | 53 | introns | )76:XM_040653623:int | XM_040653623 | MATK | LOC769076 |
| DMC | 28 | 1377212 | 163 | promoter | )28654:ENSGALT00010( | NSGALT0001006936 | NSGALG000100286 | NSGALG0001002865 |
| DMC | 28 | 1377212 | 163 | promoter | )53271:DAVISGALT0053 | AVISGALT00532710 | AVISGALG0000532 | AVISGALG00005327 |
| DMC | 28 | 1377212 | 163 | introns | )76:XM_015299784:int | XM_015299784 | MATK | LOC769076 |
| DMC | 28 | 1377212 | 163 | introns | )76:XM_015299785:int | XM_015299785 | MATK | LOC769076 |
| DMC | 28 | 1377212 | 163 | introns | )76:XM_015299786:int | XM_015299786 | MATK | LOC769076 |
| DMC | 28 | 1377212 | 163 | introns | )76:XM_040653622:int | XM_040653622 | MATK | LOC769076 |
| DMC | 28 | 1377212 | 163 | introns | )76:XM_040653623:int | XM_040653623 | MATK | LOC769076 |
| DMC | 28 | 1377558 | -183 | promoter | )53271:DAVISGALT0053 | AVISGALT00532710 | AVISGALG0000532 | AVISGALG00005327 |
| DMC | 28 | 1377558 | -183 | exons | )076:XM_015299784:ex | XM_015299784 | MATK | LOC769076 |
| DMC | 28 | 1377558 | -183 | exons | )076:XM_015299785:ex | XM_015299785 | MATK | LOC769076 |
| DMC | 28 | 1377558 | -183 | exons | )076:XM_015299786:ex | XM_015299786 | MATK | LOC769076 |
| DMC | 28 | 1377558 | -183 | exons | )076:XM_040653622:ex | XM_040653622 | MATK | LOC769076 |
| DMC | 28 | 1377558 | -183 | exons | )076:XM_040653623:ex | XM_040653623 | MATK | LOC769076 |
| DMC | 28 | 1377558 | -183 | cds | )076:XM_015299784:ex | XM_015299784 | MATK | LOC769076 |
| DMC | 28 | 1377558 | -183 | cds | )076:XM_015299785:ex | XM_015299785 | MATK | LOC769076 |
| DMC | 28 | 1377558 | -183 | cds | )076:XM_015299786:ex | XM_015299786 | MATK | LOC769076 |
| DMC | 28 | 1377558 | -183 | cds | )076:XM_040653622:ex | XM_040653622 | MATK | LOC769076 |
| DMC | 28 | 1377558 | -183 | cds | )076:XM_040653623:ex | XM_040653623 | MATK | LOC769076 |
| DMC | 28 | 1377684 | -309 | promoter | )223:XM_015299787:pr | XM_015299787 | LOC769223 | LOC769223 |
| DMC | 28 | 1377684 | -309 | promoter | )53271:DAVISGALT0053 | AVISGALT00532710 | AVISGALG0000532 | AVISGALG00005327 |
| DMC | 28 | 1377684 | -309 | introns | )76:XM_015299784:int | XM_015299784 | MATK | LOC769076 |
| DMC | 28 | 1377684 | -309 | introns | )76:XM_015299785:int | XM_015299785 | MATK | LOC769076 |
| DMC | 28 | 1377684 | -309 | introns | )76:XM_015299786:int | XM_015299786 | MATK | LOC769076 |
| DMC | 28 | 1377684 | -309 | introns | )76:XM_040653622:int | XM_040653622 | MATK | LOC769076 |
| DMC | 28 | 1377684 | -309 | introns | )76:XM_040653623:int | XM_040653623 | MATK | LOC769076 |
| DMC | 28 | 1378037 | 216 | promoter | )223:XM_015299787:pr | XM_015299787 | LOC769223 | LOC769223 |
| DMC | 28 | 1378037 | 216 | promoter | )53271:DAVISGALT0053 | AVISGALT00532710 | AVISGALG0000532 | AVISGALG00005327 |
| DMC | 28 | 1378037 | 216 | introns | )76:XM_015299784:int | XM_015299784 | MATK | LOC769076 |
| DMC | 28 | 1378037 | 216 | introns | )76:XM_015299785:int | XM_015299785 | MATK | LOC769076 |
| DMC | 28 | 1378037 | 216 | introns | )76:XM_015299786:int | XM_015299786 | MATK | LOC769076 |
| DMC | 28 | 1378037 | 216 | introns | )76:XM_040653622:int | XM_040653622 | MATK | LOC769076 |
| DMC | 28 | 1378037 | 216 | introns | )76:XM_040653623:int | XM_040653623 | MATK | LOC769076 |
| DMC | 28 | 1381156 | 53 | promoter | )223:XM_015299787:pr | XM_015299787 | LOC769223 | LOC769223 |
| DMC | 28 | 1381156 | 53 | promoter | 3893:XM_046904519:pr | XM_046904519 | RAX2 | LOC373893 |
| DMC | 28 | 1381156 | 53 | UTR5 | 23:XM_015299787:5'UT | XM_015299787 | LOC769223 | LOC769223 |
| DMC | 28 | 1381156 | 53 | exons | 9223:XM_015299787:e | XM_015299787 | LOC769223 | LOC769223 |
| DMC | 28 | 1381156 | 53 | exon1 | 9223:XM_015299787:e | XM_015299787 | LOC769223 | LOC769223 |
| DMC | 28 | 1381156 | 53 | downstream | 76:XM_015299784:dov | XM_015299784 | MATK | LOC769076 |
| DMC | 28 | 1381156 | 53 | downstream | 76:XM_015299785:dov | XM_015299785 | MATK | LOC769076 |
| DMC | 28 | 1381156 | 53 | downstream | 76:XM_015299786:dov | XM_015299786 | MATK | LOC769076 |
| DMC | 28 | 1381156 | 53 | downstream | 76:XM_040653622:dov | XM_040653622 | MATK | LOC769076 |
| DMC | 28 | 1381156 | 53 | downstream | 76:XM_040653623:dov | XM_040653623 | MATK | LOC769076 |
| DMC | 28 | 1390707 | -146 | exons | 3893:XM_046904519:e | XM_046904519 | RAX2 | LOC373893 |
| DMC | 28 | 1390707 | -146 | exons | 73893:NM_204104:exo | NM_204104 | RAX2 | LOC373893 |

| | | | | | | | | |
|---|---|---|---|---|---|---|---|---|
| DMC | 28 | 1390707 | -146 | cds | 3893:XM_046904519:e | XM_046904519 | RAX2 | LOC373893 |
| DMC | 28 | 1390707 | -146 | cds | 73893:NM_204104:exc | NM_204104 | RAX2 | LOC373893 |
| DMC | 28 | 1390707 | -146 | downstream | 51:NM_001031574:dov | NM_001031574 | PCASP2 | LOC429451 |
| DMC | 28 | 1403439 | -52 | exons | 20069:XM_418188:exo | XM_418188 | APBA3 | LOC420069 |
| DMC | 28 | 1403439 | -52 | cds | 20069:XM_418188:exo | XM_418188 | APBA3 | LOC420069 |
| DMC | 28 | 1415434 | 172 | promoter | 0072:XM_015299765:pr | XM_015299765 | PIP5K1C | LOC420072 |
| DMC | 28 | 1415434 | 172 | promoter | 0072:NM_001319021:pr | NM_001319021 | PIP5K1C | LOC420072 |
| DMC | 28 | 1415434 | 172 | promoter | 0072:XM_015299762:pr | XM_015299762 | PIP5K1C | LOC420072 |
| DMC | 28 | 1415434 | 172 | promoter | 0072:XM_015299766:pr | XM_015299766 | PIP5K1C | LOC420072 |
| DMC | 28 | 1415434 | 172 | promoter | 0072:XM_040653530:pr | XM_040653530 | PIP5K1C | LOC420072 |
| DMC | 28 | 1415434 | 172 | promoter | 0072:XR_006932216:pr | XR_006932216 | PIP5K1C | LOC420072 |
| DMC | 28 | 1415434 | 172 | promoter | 0070:XM_046904621:pr | XM_046904621 | TJP3 | LOC420070 |
| DMC | 28 | 1415434 | 172 | promoter | 0070:XM_015299757:pr | XM_015299757 | TJP3 | LOC420070 |
| DMC | 28 | 1415434 | 172 | promoter | 0070:XM_015299758:pr | XM_015299758 | TJP3 | LOC420070 |
| DMC | 28 | 1415434 | 172 | promoter | 0070:XM_015299759:pr | XM_015299759 | TJP3 | LOC420070 |
| DMC | 28 | 1415434 | 172 | promoter | 0070:XM_015299760:pr | XM_015299760 | TJP3 | LOC420070 |
| DMC | 28 | 1415434 | 172 | promoter | 0070:XM_040653624:pr | XM_040653624 | TJP3 | LOC420070 |
| DMC | 28 | 1415434 | 172 | UTR5 | 70:XM_015299759:5'UT | XM_015299759 | TJP3 | LOC420070 |
| DMC | 28 | 1415434 | 172 | UTR5 | 70:XM_015299760:5'UT | XM_015299760 | TJP3 | LOC420070 |
| DMC | 28 | 1415434 | 172 | exons | 0070:XM_015299759:ex | XM_015299759 | TJP3 | LOC420070 |
| DMC | 28 | 1415434 | 172 | exons | 0070:XM_015299760:ex | XM_015299760 | TJP3 | LOC420070 |
| DMC | 28 | 1415434 | 172 | exon1 | 0070:XM_015299759:ex | XM_015299759 | TJP3 | LOC420070 |
| DMC | 28 | 1415434 | 172 | exon1 | 0070:XM_015299760:ex | XM_015299760 | TJP3 | LOC420070 |
| DMC | 28 | 1415434 | 172 | introns | 070:XM_015299757:int | XM_015299757 | TJP3 | LOC420070 |
| DMC | 28 | 1415434 | 172 | introns | 070:XM_015299758:int | XM_015299758 | TJP3 | LOC420070 |
| DMC | 28 | 1415434 | 172 | introns | 070:XM_040653624:int | XM_040653624 | TJP3 | LOC420070 |
| DMC | 28 | 1415434 | 172 | intron1 | 070:XM_015299757:int | XM_015299757 | TJP3 | LOC420070 |
| DMC | 28 | 1415434 | 172 | intron1 | 070:XM_015299758:int | XM_015299758 | TJP3 | LOC420070 |
| DMC | 28 | 1415434 | 172 | intron1 | 070:XM_040653624:int | XM_040653624 | TJP3 | LOC420070 |
| DMC | 28 | 1427834 | 1263 | introns | 072:XM_015299765:int | XM_015299765 | PIP5K1C | LOC420072 |
| DMC | 28 | 1427834 | 1263 | introns | 072:NM_001319021:int | NM_001319021 | PIP5K1C | LOC420072 |
| DMC | 28 | 1427834 | 1263 | introns | 072:XM_015299762:int | XM_015299762 | PIP5K1C | LOC420072 |
| DMC | 28 | 1427834 | 1263 | introns | 072:XM_015299766:int | XM_015299766 | PIP5K1C | LOC420072 |
| DMC | 28 | 1427834 | 1263 | introns | 072:XM_040653530:int | XM_040653530 | PIP5K1C | LOC420072 |
| DMC | 28 | 1427834 | 1263 | introns | 072:XR_006932216:intı | XR_006932216 | PIP5K1C | LOC420072 |
| DMC | 28 | 1427834 | 1263 | introns | 09118:NONGGAT01457 | NONGGAT014572 | NONGGAG009118 | NONGGAG009118 |
| DMC | 28 | 1427834 | 1263 | intron1 | 072:XM_015299765:int | XM_015299765 | PIP5K1C | LOC420072 |
| DMC | 28 | 1427834 | 1263 | intron1 | 072:NM_001319021:int | NM_001319021 | PIP5K1C | LOC420072 |
| DMC | 28 | 1427834 | 1263 | intron1 | 072:XM_015299762:int | XM_015299762 | PIP5K1C | LOC420072 |
| DMC | 28 | 1427834 | 1263 | intron1 | 072:XM_015299766:int | XM_015299766 | PIP5K1C | LOC420072 |
| DMC | 28 | 1427834 | 1263 | intron1 | 072:XM_040653530:int | XM_040653530 | PIP5K1C | LOC420072 |
| DMC | 28 | 1427834 | 1263 | intron1 | 072:XR_006932216:intı | XR_006932216 | PIP5K1C | LOC420072 |
| DMC | 28 | 1427834 | 1263 | downstream | 3259:NONGGAT005591 | NONGGAT005591 | NONGGAG003259 | NONGGAG003259 |
| DMC | 28 | 1427834 | 1263 | downstream | 3259:NONGGAT005592 | NONGGAT005592 | NONGGAG003259 | NONGGAG003259 |
| DMC | 28 | 1525120 | -393 | promoter | 31951:TAGAGALT0000 | GAGALT0000002515 | GAGALG00000003 | GAGALG0000000319 |
| DMC | 28 | 1525120 | -393 | promoter | 28192:ENSGALT00010 | NSGALT0001006832 | NSGALG00010028 | NSGALG0001002819 |
| DMC | 28 | 1525120 | -393 | introns | 208:XM_015299672:int | XM_015299672 | NFIC | LOC396208 |
| DMC | 28 | 1525120 | -393 | introns | 208:XM_015299673:int | XM_015299673 | NFIC | LOC396208 |
| DMC | 28 | 1525120 | -393 | introns | 208:XM_015299675:in | XM_015299675 | NFIC | LOC396208 |
| DMC | 28 | 1525120 | -393 | introns | 208:XM_015299676:in | XM_015299676 | NFIC | LOC396208 |
| DMC | 28 | 1525120 | -393 | introns | 208:XM_015299677:in | XM_015299677 | NFIC | LOC396208 |
| DMC | 28 | 1525120 | -393 | introns | 208:XM_015299678:in | XM_015299678 | NFIC | LOC396208 |
| DMC | 28 | 1525120 | -393 | introns | 208:NM_001397300:int | NM_001397300 | NFIC | LOC396208 |
| DMC | 28 | 1525120 | -393 | introns | 208:NM_001397304:in | NM_001397304 | NFIC | LOC396208 |
| DMC | 28 | 1525120 | -393 | introns | 06208:NM_205271:intrc | NM_205271 | NFIC | LOC396208 |
| DMC | 28 | 1525120 | -393 | introns | 208:NM_001397302:int | NM_001397302 | NFIC | LOC396208 |
| DMC | 28 | 1525120 | -393 | introns | 208:NM_001397303:int | NM_001397303 | NFIC | LOC396208 |
| DMC | 28 | 1525120 | -393 | introns | 208:NM_001397306:in | NM_001397306 | NFIC | LOC396208 |
| DMC | 28 | 1525120 | -393 | introns | 208:NM_001397307:in | NM_001397307 | NFIC | LOC396208 |
| DMC | 28 | 1525120 | -393 | downstream | 5895:NONGGAT010118 | NONGGAT010118 | NONGGAG005895 | NONGGAG005895 |
| DMC | 28 | 1820553 | 90 | promoter | 55335:XM_046904635: | XM_046904635 | TMIGD2 | LOC107055335 |
| DMC | 28 | 1820553 | 90 | promoter | 51263:XM_025144384: | XM_025144384 | LOC107051263 | LOC107051263 |
| DMC | 28 | 1820553 | 90 | introns | 55335:XM_015299828: | XM_015299828 | TMIGD2 | LOC107055335 |
| DMC | 28 | 1820553 | 90 | introns | 55335:XM_015299829: | XM_015299829 | TMIGD2 | LOC107055335 |
| DMC | 28 | 1820553 | 90 | introns | 55335:XM_025144387: | XM_025144387 | TMIGD2 | LOC107055335 |
| DMC | 28 | 1820553 | 90 | introns | 55335:XM_046904636: | XM_046904636 | TMIGD2 | LOC107055335 |
| DMC | 28 | 1820553 | 90 | introns | 55335:XM_046904635: | XM_046904635 | TMIGD2 | LOC107055335 |

| | | | | | | | | |
|---|---|---|---|---|---|---|---|---|
| DMC | 28 | 1820553 | 90 | intron1 | 55335:XM_015299828: | XM_015299828 | TMIGD2 | LOC107055335 |
| DMC | 28 | 1820553 | 90 | intron1 | 55335:XM_015299829: | XM_015299829 | TMIGD2 | LOC107055335 |
| DMC | 28 | 1820553 | 90 | intron1 | 55335:XM_025144387: | XM_025144387 | TMIGD2 | LOC107055335 |
| DMC | 28 | 1820553 | 90 | intron1 | 55335:XM_046904636: | XM_046904636 | TMIGD2 | LOC107055335 |
| DMC | 28 | 1820553 | 90 | intron1 | 55335:XM_046904635: | XM_046904635 | TMIGD2 | LOC107055335 |
| DMC | 28 | 1820553 | 90 | downstream | 8417:ENSGALT0001006 | NSGALT0001006883 | NSGALG000100284 | NSGALG0001002841 |
| DMC | 28 | 2965916 | 178 | introns | 302:NM_001317087:int | NM_001317087 | HDGFRP2 | LOC772302 |
| DMC | 28 | 2965916 | 178 | introns | 302:XM_001235457:int | XM_001235457 | HDGFRP2 | LOC772302 |
| DMC | 28 | 2965916 | 178 | introns | 302:XM_004948799:int | XM_004948799 | HDGFRP2 | LOC772302 |
| DMC | 28 | 2965916 | 178 | introns | 302:XM_004948800:int | XM_004948800 | HDGFRP2 | LOC772302 |
| DMC | 28 | 2965916 | 178 | intron1 | 302:NM_001317087:int | NM_001317087 | HDGFRP2 | LOC772302 |
| DMC | 28 | 2965916 | 178 | intron1 | 302:XM_001235457:int | XM_001235457 | HDGFRP2 | LOC772302 |
| DMC | 28 | 2965916 | 178 | intron1 | 302:XM_004948799:int | XM_004948799 | HDGFRP2 | LOC772302 |
| DMC | 28 | 2965916 | 178 | intron1 | 302:XM_004948800:int | XM_004948800 | HDGFRP2 | LOC772302 |
| DMC | 28 | 2982729 | -116 | promoter | 5943:XM_015300019:pr | XM_015300019 | PLIN5 | LOC426943 |
| DMC | 28 | 2982729 | -116 | exons | 358066:XM_003642865 | XM_003642865 | LRG1 | LOC100858066 |
| DMC | 28 | 2982729 | -116 | exon1 | 358066:XM_003642865 | XM_003642865 | LRG1 | LOC100858066 |
| DMC | 28 | 2982729 | -116 | cds | 358066:XM_003642865 | XM_003642865 | LRG1 | LOC100858066 |
| DMC | 28 | 2985520 | -295 | promoter | 58066:XM_003642865: | XM_003642865 | LRG1 | LOC100858066 |
| DMC | 28 | 2985520 | -295 | exons | 330:NM_001397911:ex | NM_001397911 | SEMA6B | LOC428330 |
| DMC | 28 | 2985520 | -295 | exons | 330:XM_015300013:ex | XM_015300013 | SEMA6B | LOC428330 |
| DMC | 28 | 2985520 | -295 | exons | 330:XM_015300014:ex | XM_015300014 | SEMA6B | LOC428330 |
| DMC | 28 | 2985520 | -295 | cds | 330:NM_001397911:ex | NM_001397911 | SEMA6B | LOC428330 |
| DMC | 28 | 2985520 | -295 | cds | 330:XM_015300013:ex | XM_015300013 | SEMA6B | LOC428330 |
| DMC | 28 | 2985520 | -295 | cds | 330:XM_015300014:ex | XM_015300014 | SEMA6B | LOC428330 |
| DMC | 28 | 3044906 | -219 | promoter | 96349:NM_205388:pro | NM_205388 | MAP2K2 | LOC396349 |
| DMC | 28 | 3044906 | -219 | promoter | 5349:XM_015299918:pr | XM_015299918 | MAP2K2 | LOC396349 |
| DMC | 28 | 3044906 | -219 | introns | 3333:XM_004948816:in | XM_004948816 | CREB3L3 | LOC428333 |
| DMC | 28 | 3044906 | -219 | introns | 3333:XM_015300011:in | XM_015300011 | CREB3L3 | LOC428333 |
| DMC | 28 | 3479723 | -1180 | promoter | 07749:XM_040653720: | XM_040653720 | LOC121107749 | LOC121107749 |
| DMC | 28 | 3479723 | -1180 | promoter | 07749:XR_005842157:p | XR_005842157 | LOC121107749 | LOC121107749 |
| DMC | 28 | 3479723 | -1180 | promoter | 049998:TAGAGALT0000 | GAGALT0000002548 | GAGALG000000049 | GAGALG0000000499 |
| DMC | 28 | 3479723 | -1180 | promoter | 049998:TAGAGALT0000 | GAGALT0000002548 | GAGALG000000049 | GAGALG0000000499 |
| DMC | 28 | 3553302 | -5930 | introns | 95831:NM_204983:intr | NM_204983 | EFNA2 | LOC395831 |
| DMC | 28 | 3553302 | -5930 | intron1 | 95831:NM_204983:intr | NM_204983 | EFNA2 | LOC395831 |
| DMC | 28 | 3569842 | 5852 | introns | 95831:NM_204983:intr | NM_204983 | EFNA2 | LOC395831 |
| DMC | 28 | 3569842 | 5852 | intron1 | 95831:NM_204983:intr | NM_204983 | EFNA2 | LOC395831 |
| DMC | 28 | 3572252 | 3442 | introns | 95831:NM_204983:intr | NM_204983 | EFNA2 | LOC395831 |
| DMC | 28 | 3572252 | 3442 | intron1 | 95831:NM_204983:intr | NM_204983 | EFNA2 | LOC395831 |
| DMC | 28 | 3592223 | 1395 | promoter | 7267:XM_046904674:pr | XM_046904674 | CAMK4L | LOC427267 |
| DMC | 28 | 3592223 | 1395 | promoter | 7267:XM_025144406:pr | XM_025144406 | CAMK4L | LOC427267 |
| DMC | 28 | 3592223 | 1395 | promoter | 7267:XM_046904675:pr | XM_046904675 | CAMK4L | LOC427267 |
| DMC | 34 | 622361 | 226 | promoter | 5888:XM_015300251:pr | XM_015300251 | RAPGEF3 | LOC426888 |
| DMC | 34 | 622361 | 226 | promoter | 5888:XM_040654829:pr | XM_040654829 | RAPGEF3 | LOC426888 |
| DMC | 34 | 859075 | -63 | promoter | 77886:NR_031504:pro | NR_031504 | MIR196A2 | LOC777886 |
| DMC | 34 | 859075 | -63 | promoter | nassigned_transcript_1 | ssigned_transcript_1 | MIR196A2 | LOC777886 |
| DMC | 34 | 859075 | -63 | introns | 0471:NM_001305260:in | NM_001305260 | HOXC10 | LOC770471 |
| DMC | 34 | 859075 | -63 | intron1 | 0471:NM_001305260:in | NM_001305260 | HOXC10 | LOC770471 |
| DMC | 34 | 1178938 | 147 | promoter | 49457:XM_015273043: | XM_015273043 | LOC107049457 | LOC107049457 |
| DMC | 34 | 1178938 | 147 | promoter | 49457:XM_046905008: | XM_046905008 | LOC107049457 | LOC107049457 |
| DMC | 34 | 1178938 | 147 | promoter | 49457:XM_046905009: | XM_046905009 | LOC107049457 | LOC107049457 |
| DMC | 34 | 1178938 | 147 | promoter | 5211:XM_015273045:pr | XM_015273045 | ZNF853 | LOC776211 |
| DMC | 34 | 1178938 | 147 | promoter | 5211:XM_046905010:pr | XM_046905010 | ZNF853 | LOC776211 |
| DMC | 34 | 1178938 | 147 | promoter | 5183:NM_001012936:pr | NM_001012936 | ASB8 | LOC426183 |
| DMC | 34 | 1178938 | 147 | promoter | 5183:XM_025145291:pr | XM_025145291 | ASB8 | LOC426183 |
| DMC | 34 | 1178938 | 147 | UTR5 | 9457:XM_046905009:5' | XM_046905009 | LOC107049457 | LOC107049457 |
| DMC | 34 | 1178938 | 147 | exons | 049457:XM_046905009 | XM_046905009 | LOC107049457 | LOC107049457 |
| DMC | 34 | 1178938 | 147 | exon1 | 049457:XM_046905009 | XM_046905009 | LOC107049457 | LOC107049457 |
| DMC | 34 | 1178938 | 147 | introns | 49457:XM_015273043: | XM_015273043 | LOC107049457 | LOC107049457 |
| DMC | 34 | 1178938 | 147 | introns | 49457:XM_046905008: | XM_046905008 | LOC107049457 | LOC107049457 |
| DMC | 34 | 1178938 | 147 | intron1 | 49457:XM_015273043: | XM_015273043 | LOC107049457 | LOC107049457 |
| DMC | 34 | 1178938 | 147 | intron1 | 49457:XM_046905008: | XM_046905008 | LOC107049457 | LOC107049457 |
| DMC | 34 | 1791995 | -236 | promoter | 022693:INRAGALT0000 | NRAGALT0000005417 | RAGALG000000226 | RAGALG0000002269 |
| DMC | 34 | 1791995 | -236 | exons | 022693:INRAGALT0000 | NRAGALT0000005417 | RAGALG000000226 | RAGALG0000002269 |
| DMC | 34 | 1791995 | -236 | exon1 | 022693:INRAGALT0000 | NRAGALT0000005417 | RAGALG000000226 | RAGALG0000002269 |
| DMC | 34 | 1796445 | -296 | promoter | 069425:TAGAGALT0000 | GAGALT0000003078 | GAGALG000000069 | GAGALG0000000694 |
| DMC | 34 | 1796445 | -296 | downstream | 9246:XM_025145635:d | XM_025145635 | GDF11 | LOC107049246 |

| | | | | | | | | |
|---|---|---|---|---|---|---|---|---|
| DMC | 35 | 453293 | 136 | exons | 023963:INRAGALT0000 | RAGALT000000580 | RAGALG000000239 | RAGALG000000239 |
| DMC | 35 | 453293 | 136 | exons | 023963:INRAGALT0000 | RAGALT000000580 | RAGALG000000239 | RAGALG000000239 |
| DMC | 35 | 453293 | 136 | exons | 31360:XM_025146363 | XM_025146363 | LOC112531360 | LOC112531360 |
| DMC | 35 | 453293 | 136 | exons | 31360:XM_025146364 | XM_025146364 | LOC112531360 | LOC112531360 |
| DMC | 35 | 453293 | 136 | exon1 | 023963:INRAGALT0000 | RAGALT000000580 | RAGALG000000239 | RAGALG000000239 |
| DMC | 35 | 453293 | 136 | exon1 | 023963:INRAGALT0000 | RAGALT000000580 | RAGALG000000239 | RAGALG000000239 |
| DMC | 35 | 453293 | 136 | cds | 31360:XM_025146363 | XM_025146363 | LOC112531360 | LOC112531360 |
| DMC | 35 | 453293 | 136 | cds | 31360:XM_025146364 | XM_025146364 | LOC112531360 | LOC112531360 |
| DMC | 35 | 453293 | 136 | downstream | 1359:XM_040655002:d | XM_040655002 | OXA1L | LOC112531359 |
| DMC | 35 | 453293 | 136 | downstream | 1359:XM_040655003:d | XM_040655003 | OXA1L | LOC112531359 |
| DMC | JSK0100002 | 50218 | -22 | exons | 33535:XM_040656642:e | XM_040656642 | LOC112533535 | LOC112533535 |
| DMC | JSK0100002 | 50218 | -22 | cds | 33535:XM_040656642:e | XM_040656642 | LOC112533535 | LOC112533535 |
| DMC | JSK0100005 | 112357 | -93 | promoter | 00480:ENSGALT00010 | NSGALT0001000100 | NSGALG00010004 | NSGALG0001000048 |
| DMC | JSK0100005 | 112357 | -93 | promoter | 00480:ENSGALT00010 | NSGALT0001000100 | NSGALG00010004 | NSGALG0001000048 |
| DMC | JSK0100005 | 112357 | -93 | exons | 08908:XM_040657196:e | XM_040657196 | WRAP53 | LOC121108908 |
| DMC | JSK0100005 | 112357 | -93 | exons | .08908:XM_040657197 | XM_040657197 | WRAP53 | LOC121108908 |
| DMC | JSK0100005 | 112357 | -93 | cds | 08908:XM_040657196:e | XM_040657196 | WRAP53 | LOC121108908 |
| DMC | JSK0100005 | 112357 | -93 | cds | .08908:XM_040657197 | XM_040657197 | WRAP53 | LOC121108908 |
| DMC | JSK0100005 | 911 | 294 | promoter | 96046:NM_205141:pro | NM_205141 | AP4M1 | LOC396046 |
| DMC | JSK0100005 | 911 | 294 | introns | 08911:XM_040657207: | XM_040657207 | LOC121108911 | LOC121108911 |
| DMC | JSK0100005 | 7387 | -76 | promoter | 50649:XM_040657204: | XM_040657204 | LOC107050649 | LOC107050649 |
| DMC | JSK0100005 | 7387 | -76 | exons | 6046:NM_205141:exor | NM_205141 | AP4M1 | LOC396046 |
| DMC | JSK0100005 | 7387 | -76 | cds | 6046:NM_205141:exor | NM_205141 | AP4M1 | LOC396046 |
| DMC | JSK0100005 | 24379 | -57 | exons | 00075:ENSGALT00010 | NSGALT0001000012 | NSGALG00010000 | NSGALG0001000007 |
| DMC | JSK0100005 | 24379 | -57 | cds | 00075:ENSGALT00010 | NSGALT0001000012 | NSGALG00010000 | NSGALG0001000007 |
| DMC | JSK0100006 | 1700 | -90 | exons | 49770:XM_040657265:e | XM_040657265 | LOC107049770 | LOC107049770 |
| DMC | JSK0100006 | 1700 | -90 | cds | 49770:XM_040657265:e | XM_040657265 | LOC107049770 | LOC107049770 |
| DMC | JSK0100006 | 5789 | -116 | promoter | 08963:XM_040657324: | XM_040657324 | LOC121108963 | LOC121108963 |
| DMC | JSK0100006 | 5789 | -116 | promoter | 00310:ENSGALT00010 | NSGALT0001000059 | NSGALG00010003 | NSGALG0001000031 |
| DMC | JSK0100006 | 5789 | -116 | exons | 000310:ENSGALT00010 | NSGALT0001000058 | NSGALG00010003 | NSGALG0001000031 |
| DMC | JSK0100006 | 5789 | -116 | exons | 000310:ENSGALT00010 | NSGALT0001000058 | NSGALG00010003 | NSGALG0001000031 |
| DMC | JSK0100006 | 5789 | -116 | exons | 000310:ENSGALT00010 | NSGALT0001000059 | NSGALG00010003 | NSGALG0001000031 |
| DMC | JSK0100006 | 5789 | -116 | exons | 000310:ENSGALT00010 | NSGALT0001000059 | NSGALG00010003 | NSGALG0001000031 |
| DMC | JSK0100006 | 5789 | -116 | exons | 000310:ENSGALT00010 | NSGALT0001000059 | NSGALG00010003 | NSGALG0001000031 |
| DMC | JSK0100006 | 5789 | -116 | exon1 | 000310:ENSGALT00010 | NSGALT0001000058 | NSGALG00010003 | NSGALG0001000031 |
| DMC | JSK0100006 | 5789 | -116 | exon1 | 000310:ENSGALT00010 | NSGALT0001000058 | NSGALG00010003 | NSGALG0001000031 |
| DMC | JSK0100006 | 5789 | -116 | exon1 | 000310:ENSGALT00010 | NSGALT0001000059 | NSGALG00010003 | NSGALG0001000031 |
| DMC | JSK0100006 | 5789 | -116 | exon1 | 000310:ENSGALT00010 | NSGALT0001000059 | NSGALG00010003 | NSGALG0001000031 |
| DMC | JSK0100006 | 5789 | -116 | exon1 | 000310:ENSGALT00010 | NSGALT0001000059 | NSGALG00010003 | NSGALG0001000031 |
| DMC | MU179264.1 | 66370 | -323 | promoter | 006373:TAGAGALT0000 | GAGALT000002417 | GAGALG000000006 | GAGALG0000000063 |
| DMC | MU179264.1 | 66370 | -323 | promoter | 51057:XM_040656843: | XM_040656843 | MAZ | LOC107051057 |
| DMC | MU179264.1 | 66370 | -323 | promoter | 51057:XM_040656842: | XM_040656842 | MAZ | LOC107051057 |
| DMC | MU179264.1 | 66370 | -323 | exons | 006373:TAGAGALT0000 | GAGALT000002417 | GAGALG000000006 | GAGALG0000000063 |
| DMC | MU179264.1 | 66370 | -323 | exon1 | 006373:TAGAGALT0000 | GAGALT000002417 | GAGALG000000006 | GAGALG0000000063 |
| DMC | MU179273.1 | 174183 | -1465 | introns | 272:XM_040657094:intr | XM_040657094 | MYO7L2 | LOC429272 |
| DMC | MU179279.1 | 25860 | 1207 | promoter | 070380:TAGAGALT0000 | GAGALT000002935 | GAGALG000000070 | GAGALG0000000703 |
| DMC | MU179279.1 | 25860 | 1207 | promoter | 070380:TAGAGALT0000 | GAGALT000002935 | GAGALG000000070 | GAGALG0000000703 |
| DMC | MU179279.1 | 25860 | 1207 | exons | 024382:TAGAGALT0000 | GAGALT000002935 | GAGALG000000024 | GAGALG0000000243 |
| DMC | MU179279.1 | 25860 | 1207 | exons | 024382:TAGAGALT0000 | GAGALT000002935 | GAGALG000000024 | GAGALG0000000243 |
| DMC | MU179279.1 | 25860 | 1207 | exons | 024382:TAGAGALT0000 | GAGALT000002935 | GAGALG000000024 | GAGALG0000000243 |
| DMC | MU179279.1 | 25860 | 1207 | introns | 30992:XM_040657236: | XM_040657236 | MECP2 | LOC112530992 |
| DMC | MU179279.1 | 25860 | 1207 | intron1 | 30992:XM_040657236: | XM_040657236 | MECP2 | LOC112530992 |
| DMC | MU179279.1 | 37263 | -1574 | exons | 30992:XM_040657236 | XM_040657236 | MECP2 | LOC112530992 |
| DMC | MU179279.1 | 37263 | -1574 | UTR3 | 0992:XM_040657236:3' | XM_040657236 | MECP2 | LOC112530992 |
